# Differential Roles of Longevity Assurance Genes *LAG1* and *LAC1* in Regulating Endoplasmic Reticulum stress and Lipid Homeostasis in *Saccharomyces cerevisiae*

**DOI:** 10.1101/2024.11.03.621711

**Authors:** Arul Mathivanan, Vasanthi Nachiappan

## Abstract

Yeast ceramide synthesis is regulated by two homologous genes, *LAC1* and *LAG1,* each with distinct physiological roles, especially under ER stress conditions. This study examines their specific functions in lipid homeostasis and the ER stress response. Deleting *LAG1* enhances cell growth and survival under ER stress, whereas *LAC1* deletion does not provide similar resistance. In contrast, *LAG1* deletion significantly impacts phospholipid and neutral lipid metabolism without altering the expression of key ER stress response genes. In comparison, *LAG1* overexpression, unlike *LAC1* overexpression, severely impairs cell growth and viability, induces ER stress responses, disrupts phospholipid biosynthesis, alters membrane morphology, modifies neutral lipid synthesis, and reduces lipid droplet numbers. Overall, *LAG1* uniquely regulates ER stress and lipid homeostasis, independent of its function in ceramide synthesis. Understanding the specific contributions of LAG1 to lipid homeostasis and ER stress provides valuable insight into cellular stress mechanisms.

## 1) INTRODUCTION

Saccharomyces cerevisiae, also known as baker yeast, is an efficient and tractable model organism for studying eukaryotic lipid metabolism, genetics, biochemistry, and cell biology[1]. This yeast exhibits a significant degree of homology with mammals, providing valuable insights into the origins of lipid-related diseases[2].

The Endoplasmic Reticulum (ER) plays a crucial role in synthesizing proteins and the three major classes of membrane lipids: glycolipids, sterols, and sphingolipids[3]. Proper maintenance of ER morphology is essential for overall cellular homeostasis. ER stress arises from the accumulation of unfolded proteins and can be exacerbated by lipid bilayer stress, leading to prolonged and unresolved ER stress even when protective mechanisms like ER-associated degradation (ERAD) and the unfolded protein response (UPR) are active[4]. This stress can result in abnormal ER elongation and disrupt proteostasis. While substantial evidence links phospholipid imbalances to alterations in ER structure and stress, the specific role of ceramides in maintaining ER function remains poorly understood.

Ceramides are N-acyl amide derivatives of sphingolipid bases, and sphingolipids constitute 10– 15% of membrane lipids[5]. Our understanding of ceramide synthesis has improved significantly with the identification of two highly homologous ceramide synthase genes, *LAG1* and *LAC1*, in yeast. Systematic screening of the yeast mutation library to assess replicative lifespan (RLS) revealed that deletion of *LAG1* results in a 50% increase in RLS, marking it as the first longevity assurance gene (LAG). In contrast, *LAC1*, a homolog of *LAG1*, does not affect RLS, suggesting it has a different role in cellular physiology[6].

The ORM family proteins (Orm1p and Orm2p) reside in the ER and are key regulators of sphingolipid metabolism and proteostasis[7]. Although the interactions between the ceramide-synthesizing genes (*LAG1* and *LAC1*) and ORM genes (*ORM1* and *ORM2*) have been extensively studied, further research is necessary to elucidate their roles in regulating ER stress and lipid metabolism[8], [9]. ORM genes have functional counterparts in higher organisms, with ORMDL3 linked to asthma responses. The TOR pathway, which senses nutrient depletion, activates *ORM1* via the TAP42 phosphatase complex, leading to the negative regulation of sphingolipid synthesis. Further research on the interplay between ORM and LAG genes is essential, as it holds significant potential for advancing our understanding of cellular responses to stress and the pathogenesis of various human diseases[10].

This study specifically investigates the role of LAG genes, particularly LAG1, in modulating lipid homeostasis and influencing ER morphology. Our findings provide compelling evidence for the distinct functions of *LAC1* and *LAG1*, highlighting their specific contributions to cellular processes.

## 2) MATERIALS AND METHODS

### 2.1. Chemicals

The media utilized in this study comprised YPD (Yeast extract, peptone, and dextrose), Yeast Nitrogen Base (YNB) with ammonium sulfate, yeast dropout media, and uracil-free media, all sourced from HiMedia and Sigma. Tunicamycin, Aureobasidin, Myriocin, and Fumonisin were also acquired from Sigma. Silica gel 160F254 TLC plates and solvents were obtained from Merck, while lipid standards were sourced from Avanti Polar Lipids (Alabaster, AL). For molecular biology applications, Trizol for RNA isolation, oligo dT primers, a cDNA synthesis kit, SYBR Green PCR master mix, Nile red, and DiOC6 were purchased from Invitrogen unless otherwise specified. The anti-Pgk1 and anti-GFP antibodies were sourced from Invitrogen. Additionally, the anti-Kar2 antibody was generously provided by Prof. J. L. Brodsky from the Department of Biological Sciences at the University of Pittsburgh, PA, USA, while the anti-Cpy1 antibody was a gift from Prof. Neta Dean, Department of Biochemistry and Cell Biology at Stony Brook University, New York, NY, USA.

### 2.2. Growth conditions

Cells were cultivated in YPD media containing 1% yeast extract powder, 2% peptone, and 2% glucose and incubated at 30°C and 180 rpm until the mid-log phase was reached. In subsequent experiments, 2 OD (A_600nm_) units of cells from this starter culture were transferred into SC media, which consisted of a 2% synthetic complete mixture, 0.67% yeast nitrogen base, and 2% glucose, and allowed to grow under the same conditions until reaching the mid-log phase. Complementation strains transformed with the -pRS416 plasmid were cultivated in SC-Ura medium lacking uracil, also with 2% glucose, until they reached the mid-log phase. Cells expressing the pYES2 plasmid were grown in SC-Ura medium supplemented with 2% galactose under the same conditions. For chemical treatment experiments, cells were cultured in SC media 6 hours before reaching the mid-log phase and treated with either 1, 2, or 4 µg/ml of tunicamycin (Tm), 0.25 µg/ml of myriocin (Myr), 0.3 mM of Fumonisin B1 (FB1), or 0.8 µg/ml of Aureobasidin A (AbA). After treatment, the cells grew for an additional 6 hours. The yeast strains used in this study are listed in Table S1 in the supplementary section.

### 2.3. Growth Analysis

Cells were grown in appropriate SC media to analyze growth curves, with optical density (OD) measurements taken every 6 hours over 72 hours to monitor cell proliferation. The resulting data were then plotted to illustrate the growth trends.For the spot test analysis, cells were initially cultivated in YPD media until they reached the mid-log phase, after which the cell density was adjusted to 3.0 OD at A600 nm. The cells were diluted ten-fold with water, and 3 µl of each dilution was plated onto agar plates containing either SC or SC-Ura (2% agar), supplemented with 2% glucose or galactose. The appropriate concentrations of chemicals (tunicamycin, AbA, FB1, and myriocin) were added to the media before preparing the plates. After conducting the spot test, the plates were incubated at 30°C for 48 hours, and the results were photographed afterward.Cell suspensions derived from the YPD-grown starter culture were adjusted for the chemical sensitivity growth assay to an OD600 nm of 0.1. These suspensions were added to 200 µl of the corresponding SC culture containing the designated concentrations of the aforementioned chemicals in the wells of a 96-well plate. The cells were cultured at 30°C with shaking at 180 rpm, with OD600 nm readings taken every 2 hours using an Alta ADX-150 ELISA plate reader. Growth curves were generated from three independent experiments, and the average results from these replicates were presented.

### 2.4. Yeast transformation using Lithium acetate (LiOAc) method

To prepare yeast cells for transformation, 2.0 OD of cells were cultured in YPD media until the OD reached between 0.8 and 1.0/ml, typically taking 3 to 4 hours. Once the desired OD was achieved, the cells were harvested, washed, and resuspended in 500 µl of a Tris-EDTA-LiOAc solution. Following two additional washes, the cells were resuspended in a Tris-LiOAc-Polyethylene Glycol solution. 5 µl of single-stranded carrier DNA and the relevant plasmid (pRS416 or pYES2) were added to this cell suspension. The mixture was incubated on a shaker at 30 °C and 180 rpm for 30 minutes. After incubation, 20 µl of DMSO was added, and the cells underwent a heat shock at 42 °C for 15 minutes, followed by immediate cooling in an ice bath. The cells were then pelleted, resuspended in 0.1 ml of sterile water, and plated onto SC-Ura plates using a sterile L-rod. The plates were incubated at 30°C until the transformed cells appeared, typically within 3 to 4 days. Finally, the transformed cells were cultured for further analysis[11].

### 2.5. Cell viability assay

Cells from a starter inoculum were washed with sterile water and then transferred to the appropriate synthetic complete (SC) media at a density of 2.0 OD units. They were allowed to grow for 8 or 5 days, with samples taken every 24 hours during the growth period. To evaluate cell viability, 1,000 cells from each time point were counted using a hemocytometer and plated on appropriate SC or SC-Ura plates, with and without the specified chemicals. The plates were incubated at 30°C for 2 days to facilitate colony formation. The plates were photographed after incubation, and colony-forming units (CFU) were counted manually. The results were presented in graphical form, with cell viability calculated as a percentage of the CFU observed on day 1, which was set as 100% for the wild-type (WT) or the respective vector control sample.

### 2.6. Lipid extraction and Thin-layer chromatography

Cells were cultured, and equal optical density (OD) of cells was utilized for lipid extraction following the Bligh and Dyer method[12]. Total lipids were extracted using a mixture of chloroform, ethanol, and 2% ortho-phosphoric acid in a ratio of 2:1:1 (V/V). For the separation of lipids, phospholipids (PL) were loaded onto a TLC plate and separated using a solvent mixture of chloroform, ethanol, acetic acid, and distilled water (85:15:10:3.5, V/V). In contrast, non-polar neutral lipids (NL) were separated using a solvent of petroleum ether, diethyl ether, and acetic acid (70:30:1, V/V). Lipids were visualized on the TLC plates after exposure to iodine vapor, and individual PL and NL were identified by comparing their Rf values to standard Rf values. Phospholipid quantification involves estimating phosphorus content. Scraped PL spots were digested using perchloric acid to release phosphate ions, which formed a colored “phosphomolybdate complex.” The absorbance of this complex was proportional to the phosphorus content, allowing for the quantification of phospholipids. Subsequently, the TLC plates were scanned, and Image J software was used to quantify the neutral lipids based on the intensity of the bands observed on the plates.

### 2.8. Microscopic analysis

After culturing the yeast cells as previously described, they were collected at the mid-log phase and fixed with a 2% formaldehyde solution. The cells were then washed with phosphate-buffered saline (1X PBS). For membrane staining, 5 µl of DiOC6 (1 mg/ml) was added to 1 ml of cell suspension, which was incubated in the dark for 10 minutes, followed by three washes with 1X PBS. The fixed cells were then placed on a slide for imaging. Images were acquired with excitation and emission wavelengths of 482 nm and 504 nm, respectively.

For lipid droplet analysis, Nile Red (20 µg/ml) was added to the cells and incubated at room temperature in the dark for 15 minutes. The excitation and emission wavelengths for Nile Red were 480 nm and 510 nm, respectively. Following incubation, the cells were washed three times with 1X PBS. Cells were transformed with the pRS314-UPRE-GFP plasmid using the LiOAc method. This plasmid contains a GFP reporter gene driven by the UPRE promoter, activated during ER stress and UPR induction. The cells were grown in SC-Trp (tryptophan) medium with 2% glucose for 24 hours. After incubation, the cells were harvested and washed with 1X PBS. Subsequently, images were captured using a laser scanning confocal microscope with excitation and emission wavelengths set at 488 and 509 nm, respectively. Fluorescent images were captured using a Zeiss LSM 710 confocal microscope with a 100x/1.40 oil objective and an AxioCam camera.

### 2.9. RNA isolation, cDNA construction, and Real-Time PCRD

Yeast cultures were harvested, and cells were lysed using TRIzol reagent. RNA was extracted from the upper aqueous phase after phase separation and centrifugation. For cDNA synthesis, 2 µg of total RNA was used with the High-Capacity cDNA Reverse Transcription Kit from Applied Biosystems. PCR amplification was performed on an Applied Biosystems Step One Plus™ Real-Time PCR machine, with fluorescence continuously monitored for target gene quantification. Actin was the loading control, and gene expression levels were quantified using the 2^(-ΔΔCt)^ method[13], [14]. The gene primers used in this study are listed in Table S2 of the supplementary section.

### 2.10. *HAC1* mRNA splicing assay

Following RNA isolation from cultured cells as previously described, reverse transcription was performed using the SuperScript II kit to generate cDNA according to the manufacturer’s instructions. The resulting cDNA served as a template for PCR amplification with specific primers and Taq DNA polymerase. The PCR products were then analyzed using an agarose gel for further examination[15]. Details of the gene primers used in this study are provided in Table S2 of the supplementary section.

### 2.11. Protein isolation and Western blotting

Yeast cells were cultured as previously described. The cells were then collected and pelleted by centrifugation. Glass beads were added to the pellet for bead beating, and an extraction buffer containing Tris-HCl, EDTA, and PMSF was used to lyse the cells. The cell mixture was vortexed for 30 seconds and then placed on ice for 30 seconds, repeating this cycle 30 times. The resulting lysate was centrifuged at 10,000 rpm at 4°C to collect the upper layer containing the proteins. Protein concentration was determined using the Bradford assay, with BSA as the standard, to confirm sufficient protein for downstream applications. Equal amounts of protein were loaded onto an SDS-PAGE gel and transferred onto a nitrocellulose membrane for protein blotting. The blots were incubated with primary antibodies under gentle rocking for 12 hours, followed by incubation with secondary antibodies for 2 hours before development with BCIP/NBT substrate. Pgk1p was used as the loading control to normalize protein levels. The anti-Kar2p primary antibody was used at a 1:5000 dilution, with a 2.5:5000 dilution of the anti-mouse secondary antibody, both resolved on a 12% gel. The anti-GFP primary antibody was used at a 2.5:5000 dilution, paired with the anti-mouse secondary antibody, and resolved on a 12% gel. For anti-Cpy1p detection, the primary antibody was diluted to 2:5000, and an anti-rabbit secondary antibody was used at a 2.5:5000 dilution, resolved on a 10% gel.

### 2.12. Data quantification and Statistical analysis

Data quantification was performed using ImageJ software, and statistical analysis was conducted with PRISM software version 9.0. One-way or two-way ANOVA was applied depending on the dataset. Error bars represent the mean ± standard deviation (SD) from three independent experiments. Significance levels in the figures are indicated as * for p < 0.05, ** for p < 0.01, and *** for p < 0.005.

## 3. RESULTS

### Role of *LAG1* and *LAC1* in modulating ER Stress resistance and cell survival

#### 3.1. *LAG1* deletion enhances ER stress resistance in cells

To examine the roles of *LAC1* and *LAG1* in the cellular response to ER stress, growth curve analyses were performed on wild-type (WT), *lac1*Δ, and *lag1*Δ strains. Under normal conditions, both *lac1*Δ and *lag1*Δ cells exhibited growth rates comparable to WT cells (Fig.1A). However, when exposed to tunicamycin. This compound induces ER stress by disrupting N-glycosylation and causing the accumulation of misfolded proteins[16]. *lag1*Δ cells displayed enhanced growth relative to WT and *lac1*Δ cells (Fig.1B).

**Figure 1.**
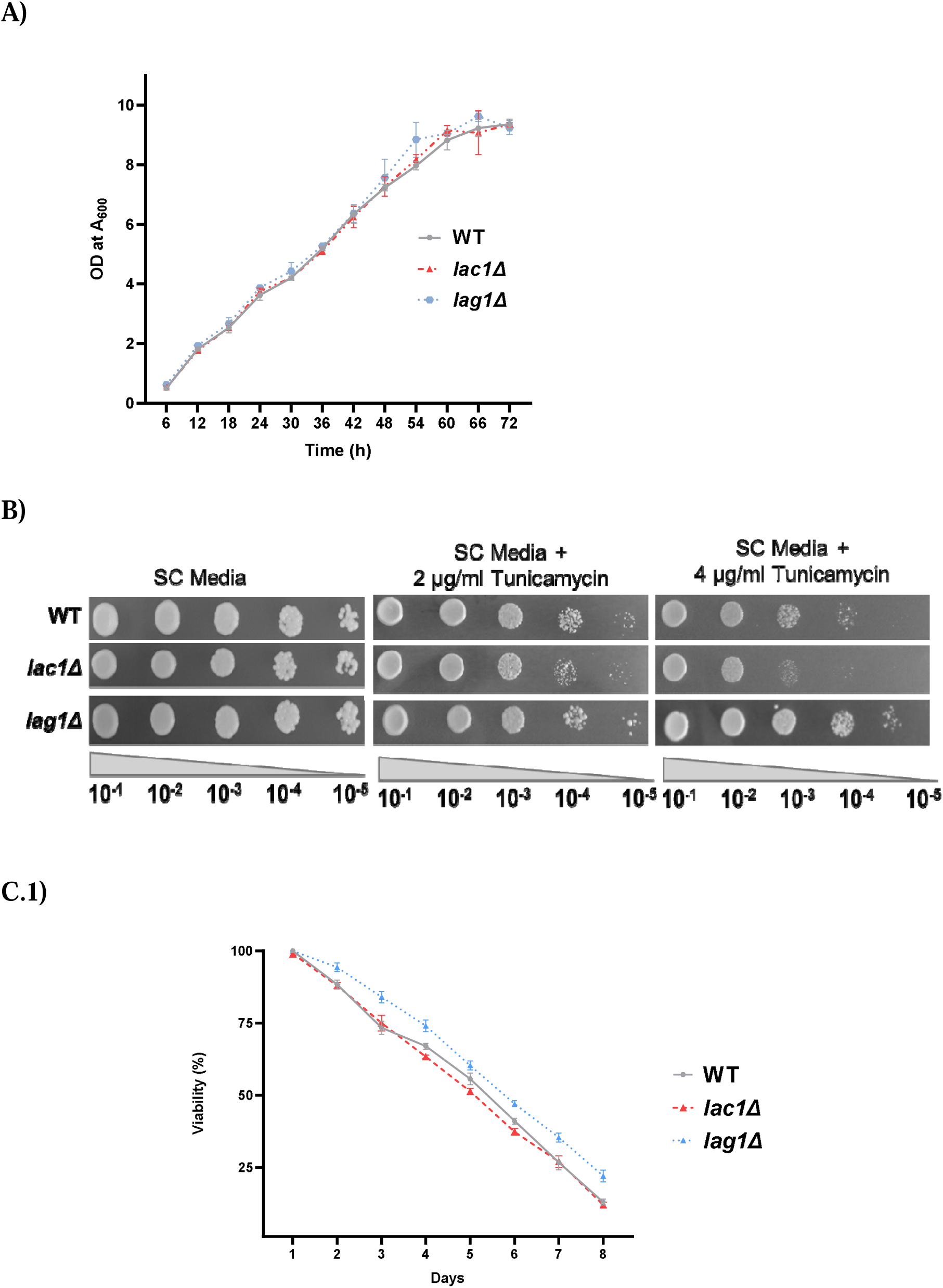
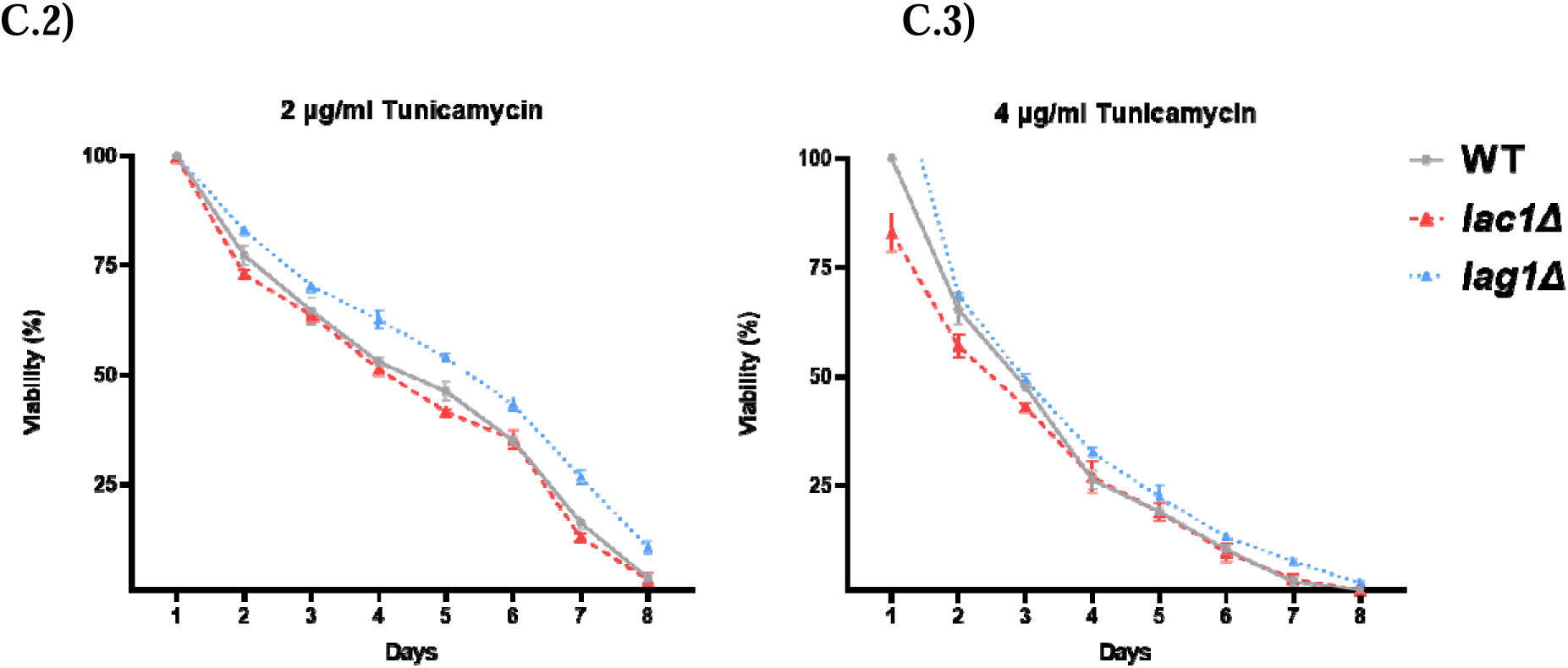
The deletion of *LAG1* enhances cell growth, viability, and resistance under ER stress conditions. (A) Growth curves of wild-type (WT), *lac1*Δ, and *lag1*Δ cells were plotted by measuring the optical density at 600 nm (OD□□□) at regular intervals over 72 hours. (B) For spot growth assays, cells adjusted to the same OD□□□ of 3.0 were serially diluted 10-fold and spotted onto SC agar plates with or without tunicamycin. The plates were incubated at 30 °C for 48 hours. (C) Cells were cultured in SC media, and equal numbers of cells (10^3^) were collected and grown at varying intervals from 1 to 8 days. Using a hemocytometer, the cells were counted and plated using a sterile, autoclaved L-rod onto SC media containing 2% dextrose (+ or - tunicamycin). The plated cells were incubated at 30□°C for 48 hours to allow growth. Cell viability of the *LAC1* and *LAG1* deletion strains was calculated relative to the colony-forming units (CFUs) of wild-type cells, which were set at 100% viability. Data represent the mean ± SD from three independent experiments (*P□<□0.05).

Cells were cultured in tunicamycin concentrations ranging from 0.5 µg/ml to 6 µg/ml (data not shown) to determine the optimal level for detecting growth differences among the strains. Significant growth differences were observed at 4 µg/ml, where *lag1*Δ cells exhibited notable resistance to ER stress-induced growth inhibition, affecting both WT and *lac1*Δ cells. A concentration of 1 µg/ml effectively triggered an ER stress response, while higher concentrations were used solely further to differentiate the responses of *LAG1* deletion cells. These findings suggest that *LAG1* deletion resists ER stress-related growth defects in WT and *lac1*Δ strains.

To further investigate the role of LAG1 in ER stress resistance, cells were treated with dithiothreitol (DTT), which induces ER stress by reducing protein disulfide bonds and promoting misfolded protein accumulation[17]. Under these conditions, *lag1*Δ cells again showed enhanced resistance to DTT-induced ER stress compared to WT and *lac1*Δ cells (Fig.S1), indicating lag1 deletion showed a protective effect over ER stress and misfolded protein accumulation to an extent.

For conformation, *lag1*Δ cells were complemented by expressing *LAG1* from the yeast centromere vector pRS416 and subjected to ER stress. This complementation reversed the ER stress resistance associated with *LAG1* deletion (Fig.S2), reinforcing the relationship between *LAG1* loss and ER stress resilience. Cell viability assays further revealed that *lag1*Δ cells had higher survival rates than WT and *lac1*Δ cells (Fig.1C.1). Notably, at tunicamycin concentrations of 2 and 4 µg/ml, *lag1*Δ cells exhibited significantly increased viability compared to WT and *lac1*Δ cells (Fig.1C.2 and 1C.3). Together, these findings confirm that *LAG1* deletion enhances cell survival under conditions of severe ER stress.

#### 3.2. *LAC1* and *LAG1* are upregulated under ER stress, each serving distinct, non-overlapping roles

Besides their role in promoting longevity, *LAC1* and *LAG1* are essential for synthesizing ceramides from sphingosine[18]. While their functions in ceramide synthesis were previously considered redundant, recent studies have revealed distinct substrate specificities for these genes[19]. Notably, only *LAG1* plays a crucial role in resisting ER stress. Gene and protein expression levels were assessed in WT cells to analyze the expression patterns of *LAG1* and *LAC1* under ER stress. *LAG1* and *LAC1* showed a significant increase in expression following ER stress induction (Fig.2A and 2B). Despite the increased *LAC1* expression, it did not contribute to countering ER stress (Fig.1B and 2B).

**Figure 2.**
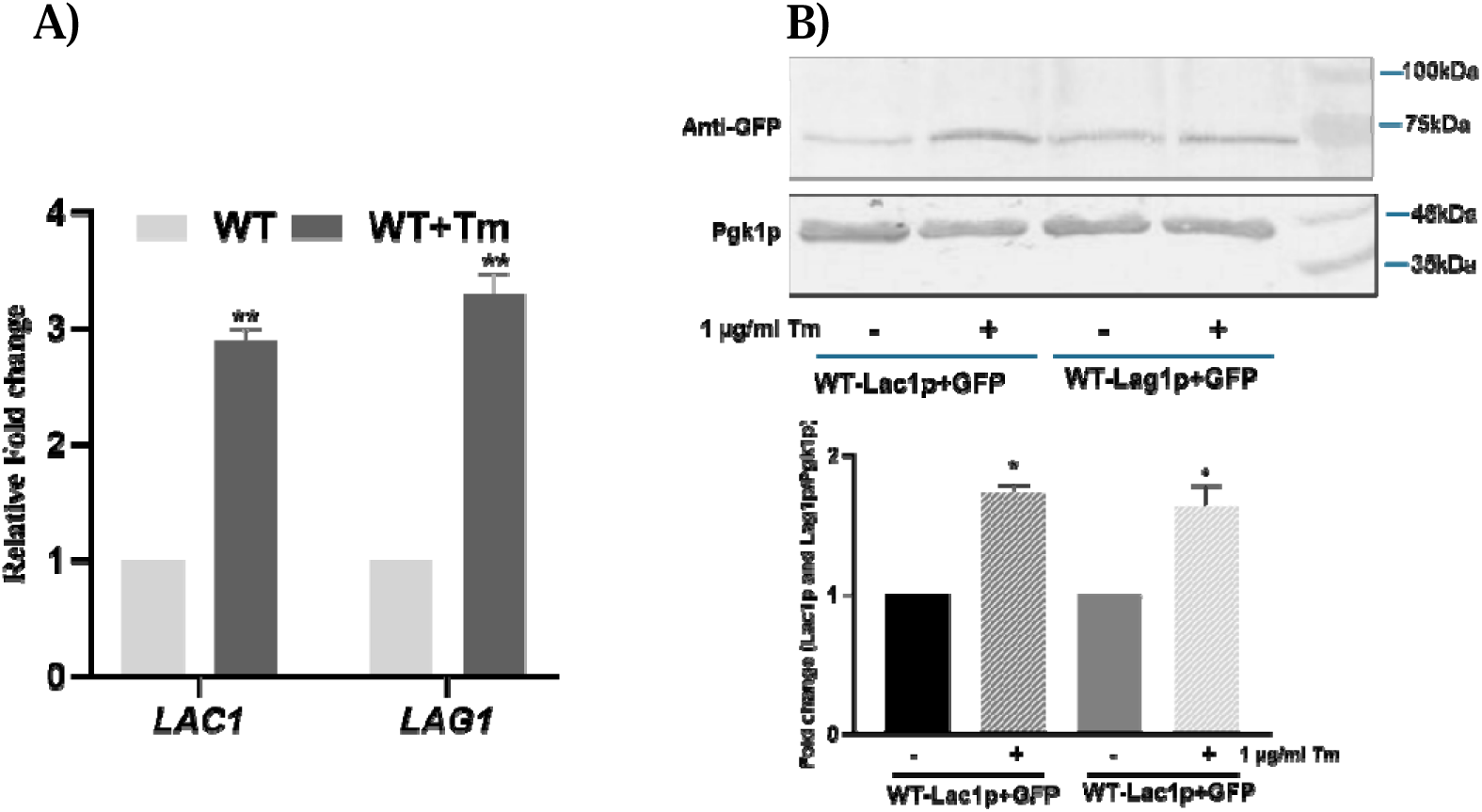
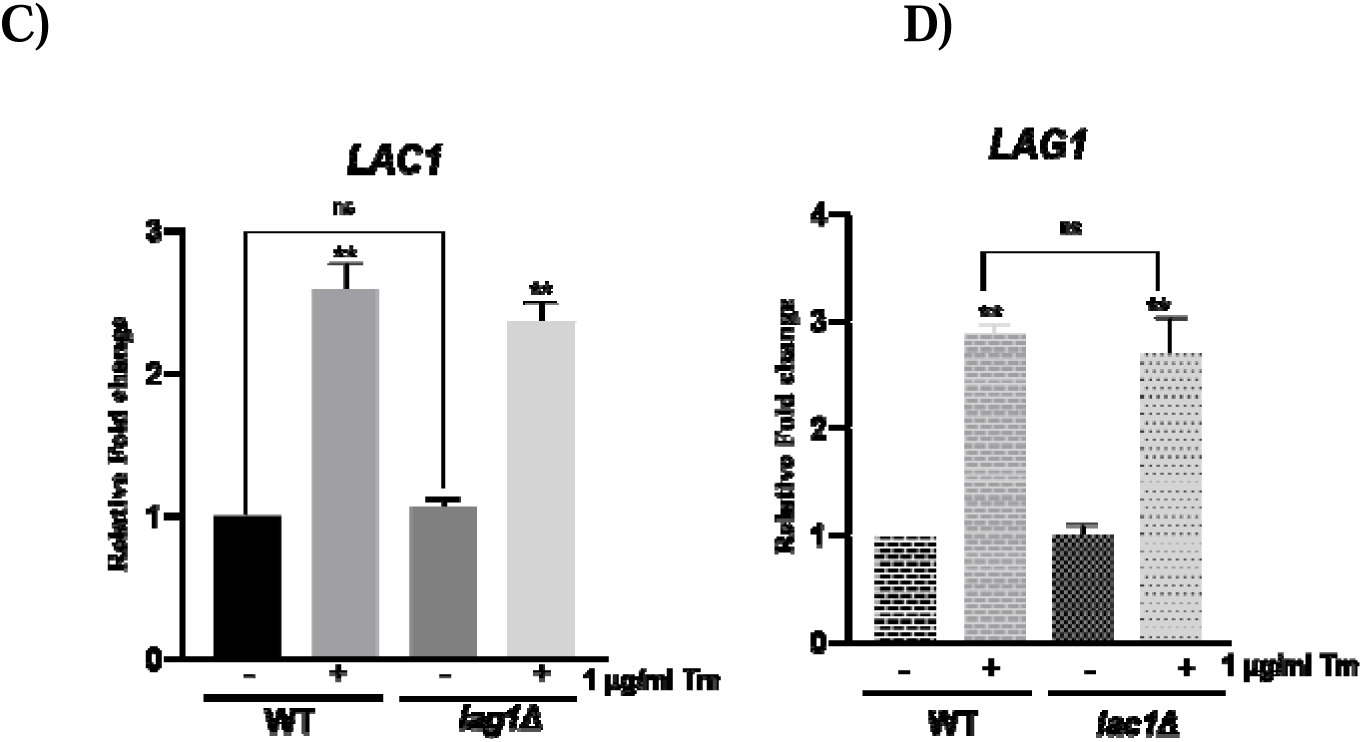
Tunicamycin-induced ER stress upregulates the expression of LAG genes, and the presence of *LAG1* or *LAC1* does not compensate for the absence of the other. Cells were grown in SC media until the mid-log phase (+ or – tunicamycin), after which mRNA and proteins were extracted for analysis. (A) The expression levels of *LAC1* and *LAG1* genes were assessed in WT cells using qRT PCR, and 2^-ΔΔCT^ values were calculated and plotted. (B) Equal amounts of protein from WT cells with endogenous GFP-tagged *LAC1*, regulated by its native promoter, and from WT cells with GFP-tagged *LAG1*, also under the control of its native promoter, were separated on a 12% SDS-PAGE gel. Pgk1p was used as a loading control, and an anti-GFP antibody was the primary detection antibody. (C) *LAC1* gene expression was analyzed in WT and *lag1*Δ cells. (D) *LAG1* gene expression was analyzed in WT and *lac1*Δ cells. Data represent the mean□±□SD from three independent experiments (*P□<□0.05).

To determine whether *LAG1* and *LAC1* have overlapping roles, the expression of the *LAC1* gene in both WT and *lag1*Δ cells was examined under ER stress. *LAC1* expression remained unchanged in both cell types, even under stress conditions (Fig.2C). Similarly, *LAG1* expression in wild-type and *lac1*Δ cells did not show significant alterations under ER stress (Fig.2D). The lack of substantial changes in expression for both genes under stress conditions indicates that *LAG1* and *LAC1* may have distinct functions and regulatory mechanisms that operate independently in response to ER stress. These findings underscore the unique and non-redundant roles of *LAG1* and *LAC1* in this context.

### *LAG1* and *LAC1* deletion altered phospholipid and neutral lipid homeostasis without affecting the ER Stress response

#### 3.3. Deletion of *LAG1* had no significant effects in altering ER stress response

Proteins are predominantly synthesized in the rough endoplasmic reticulum (RER) and require proper folding, post-translational modifications, and delivery to their designated subcellular locations to perform their functions[20]. When misfolded proteins accumulate unchecked, ER stress responses are activated to promote the proper folding of these proteins. Misfolded proteins must either be correctly refolded or degraded if refolding fails[21].

The unfolded protein response (UPR) is a cellular mechanism that promotes proper protein folding in response to misfolded or unfolded proteins in the ER. A key regulator of the UPR is *IRE1*, an ER transmembrane protein that activates UPR signaling by initiating the splicing of *HAC1* mRNA. Through this process, *IRE1* splices the uninduced, unspliced (*HAC1*^u^) mRNA by removing an intron in the cytoplasm via an unconventional splicing mechanism. The exons are then joined by tRNA ligase to produce the spliced, induced (*HAC1*^i^) mRNA. This mRNA is translated into the active form of Hac1p[15], [22]. This active transcription factor then translocates to the nucleus, binds to unfolded protein response elements (UPRE), and activates the expression of UPR target genes (see Fig. 3B).

**Figure 3.**
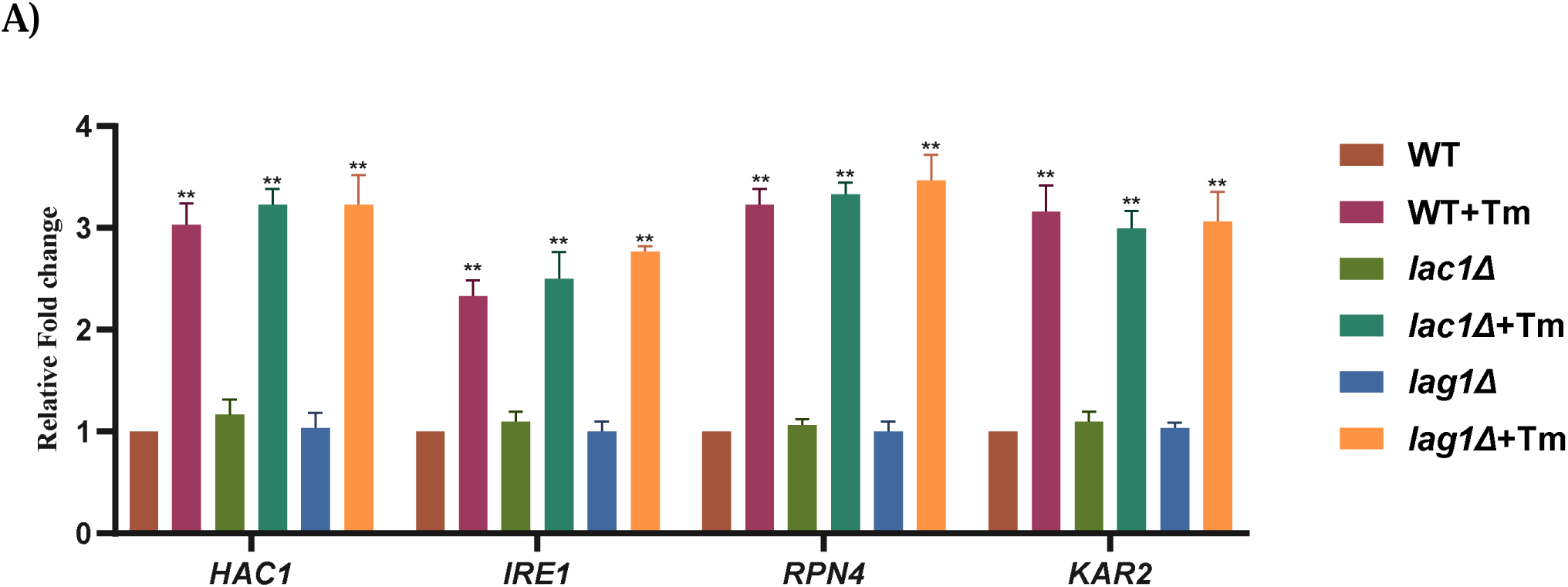
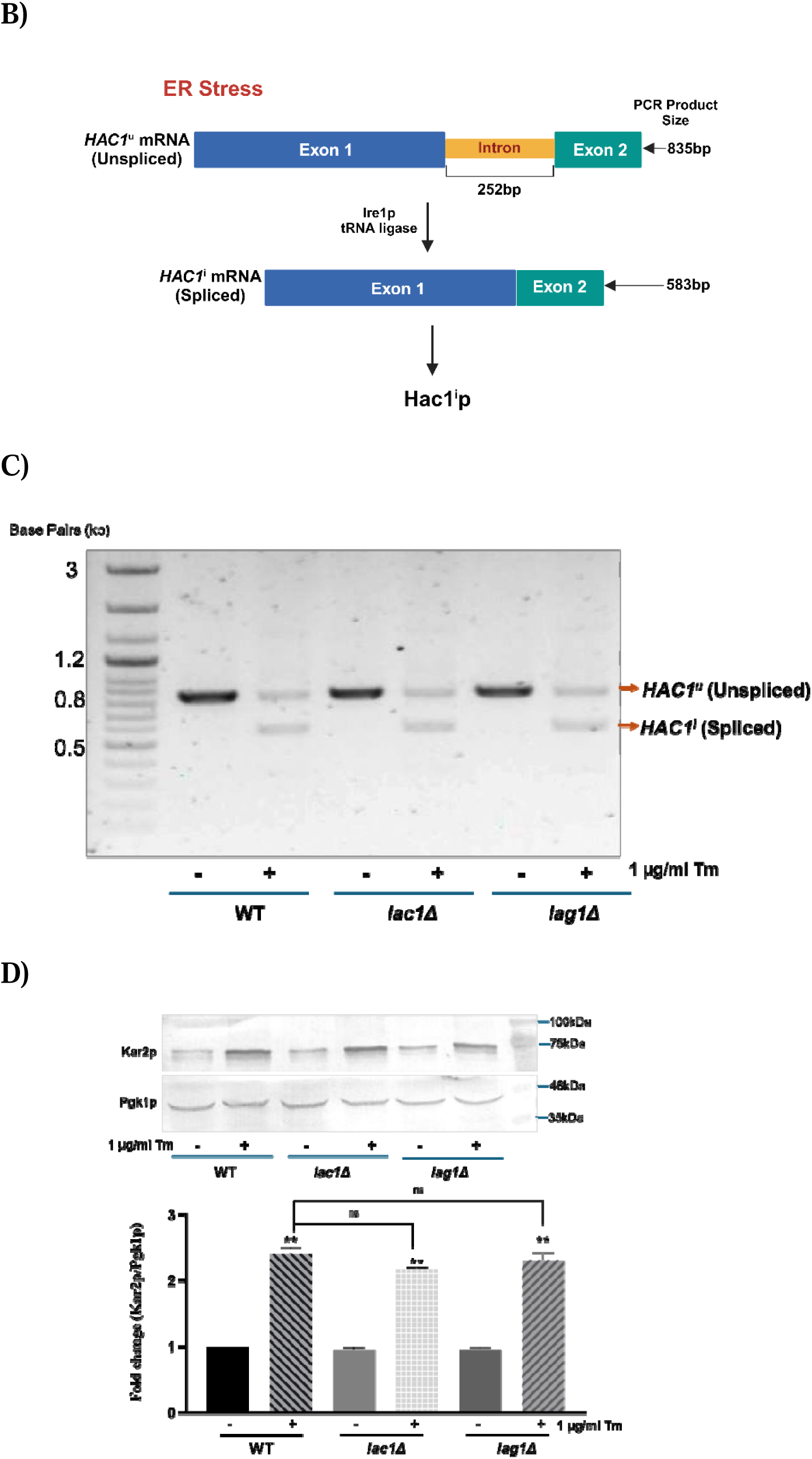
The ER stress response remains unchanged regardless of the presence or absence of the LAG genes. (A) Expression levels of ER stress response genes were analyzed using qRT PCR. (B) A schematic model of *HAC1* splicing. (C) The *HAC1* mRNA splicing assay was performed using Superscript II reverse transcriptase, and the PCR products were analyzed on a 1% agarose gel. (D) Equal amounts of protein were loaded onto a 12% SDS-PAGE gel for each sample. Pgk1p served as the loading control, and anti-Kar2p was the primary antibody. Data represent the mean□±□SD from three independent experiments (*P□<□0.05).

Beyond the UPR, ER-specific sorting signals on newly synthesized proteins are recognized by *KAR2*, an ER luminal chaperone[23]. *KAR2* facilitates the translocation of these proteins to appropriate subcellular compartments and assists in refolding misfolded proteins, a function dependent on its interaction with *IRE1*. Thus, *IRE1* is essential for maintaining the chaperone activities of *KAR2*[23]. Despite mechanisms ensuring correct protein folding, proteins that fail to achieve their proper conformation are directed toward proteasomal degradation. *RPN4*, a transcription factor, regulates the expression of proteasomal subunits, thereby controlling the degradation of misfolded or unwanted proteins[24].

Three experiments were performed to investigate ER stress induction: qRT-PCR analysis of ER stress response genes, *HAC1* splicing assay, and Kar2p expression analysis using western blotting. For this purpose, WT, *lac1*Δ, and *lag1*Δ cells were treated with tunicamycin to induce ER stress and analyzed alongside untreated controls. The results were as follows:

qRT-PCR analysis of the ER stress response genes *IRE1, HAC1, KAR2,* and *RPN4* revealed that, under ER stress, gene expression levels increased consistently across WT, *lac1*Δ, and *lag1*Δ cells, while in non-stress conditions, expression levels remained comparable among all three cell types (Fig. 3A). To investigate further HAC1 splicing, primers targeting the intron-flanking region of HAC1 mRNA were used. Reverse transcription semi-quantitative PCR in WT, *lac1*Δ, and *lag1*Δ cells showed a 252-base-pair size difference in the mRNA of ER stress-induced cells (*HAC1*^i^), indicating the activation of the unfolded protein response (UPR). No *HAC1*(*HAC1*^u^) splicing was observed in any cell type under non-stress conditions (Fig. 3C).

Finally, western blotting to analyze Kar2p expression demonstrated that, under ER stress, Kar2p levels were similarly elevated in all cell types, with *lac1*Δ and *lag1*Δ cells exhibiting expression patterns comparable to WT cells in non-stress conditions. While the loss of *LAG1* provides a protective effect against ER stress, as shown in Figure 1, it does not alter the cellular mechanisms involved in the ER stress response. This indicates that *LAG1* loss offers protection without disrupting the fundamental response mechanisms to ER stress.

#### 3.4. Loss of LAG genes modified the phospholipid levels, expression of its synthesizing genes, and membrane morphology

To examine the role of *LAG* genes (*LAG1* and *LAC1*) in phospholipid (PL) metabolism, phospholipids were isolated and assessed through thin-layer chromatography (TLC). The primary phospholipids, phosphatidylcholine (PC) and phosphatidylethanolamine (PE), crucial structural components of the cellular lipid bilayer[25], were quantified in deletion strains. In *lac1*Δ cells, PC and PE levels were diminished relative to WT cells (Fig. 4A and B). In contrast, *lag1*Δ cells displayed a more substantial reduction in both PC and PE, emphasizing the critical role of *LAG1* in sustaining these phospholipid levels. Under ER stress conditions, PL levels were elevated across all strains; however, deletion strains showed a less pronounced increase than WT cells. Notably, the stress-induced increase in PC was significantly lower in *lag1*Δ cells compared to *lac1*Δ cells, while the increase in PE under stress was comparable between *lag1*Δ and *lac1*Δ strains.

**Figure 4.**
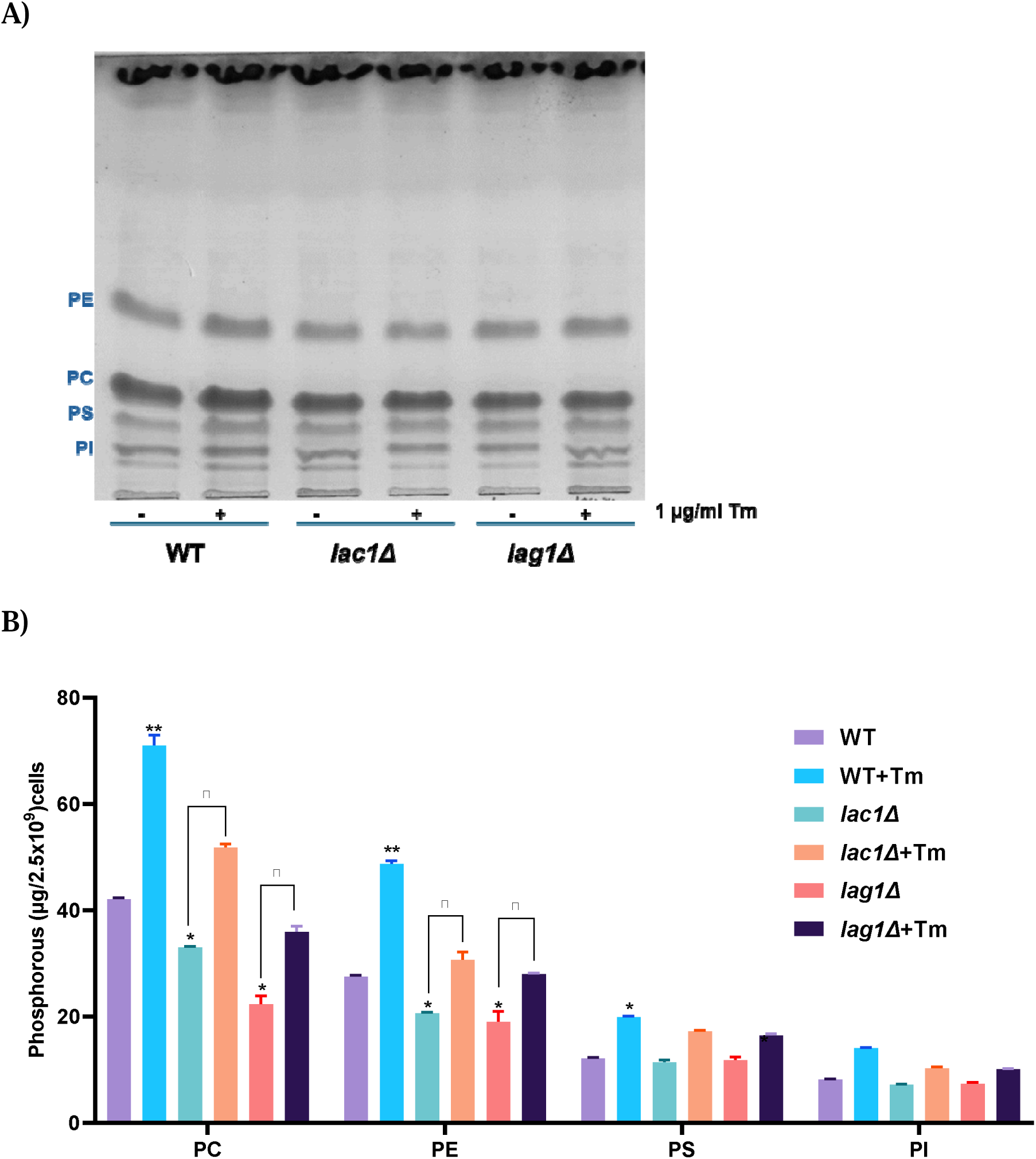
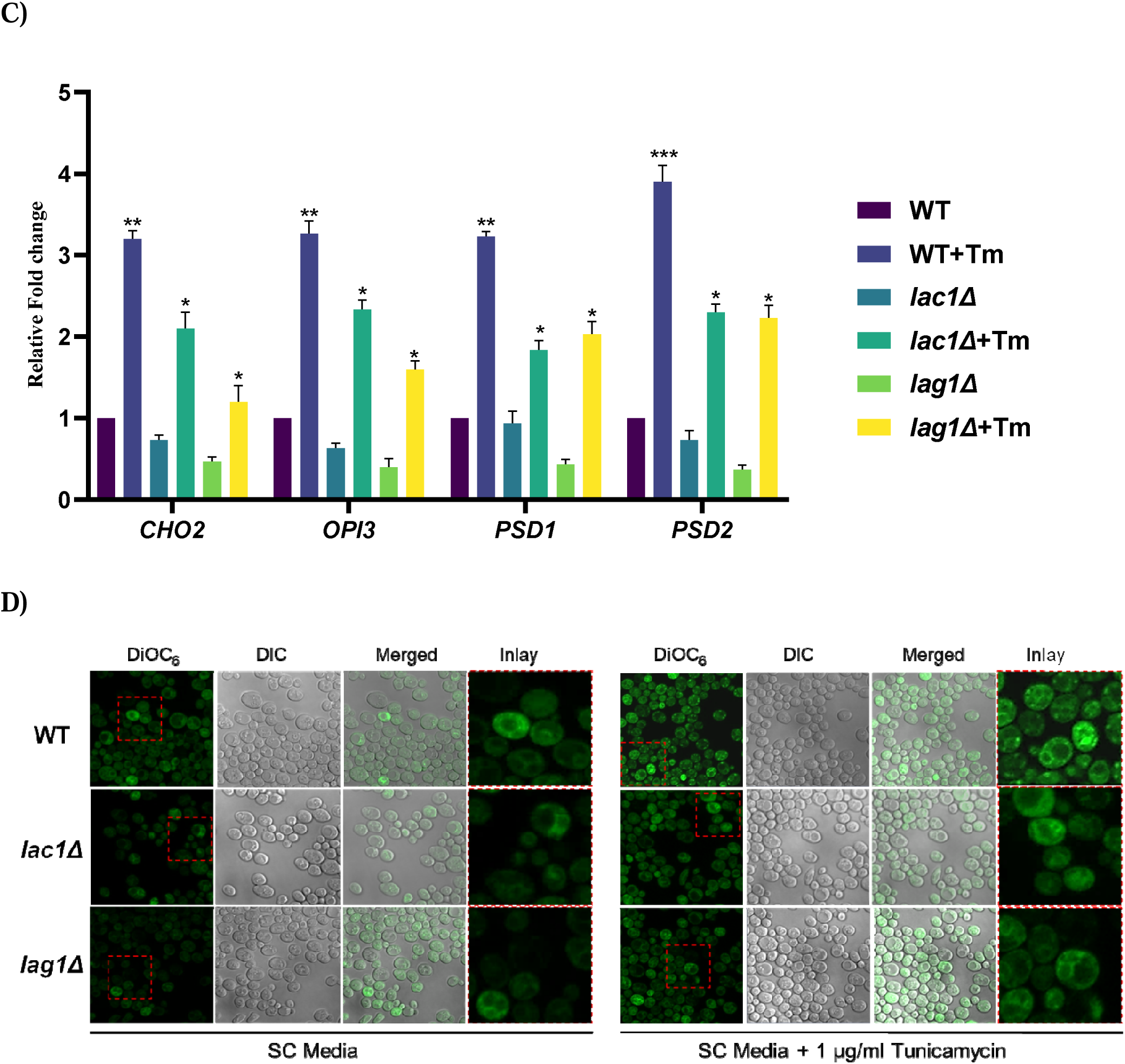
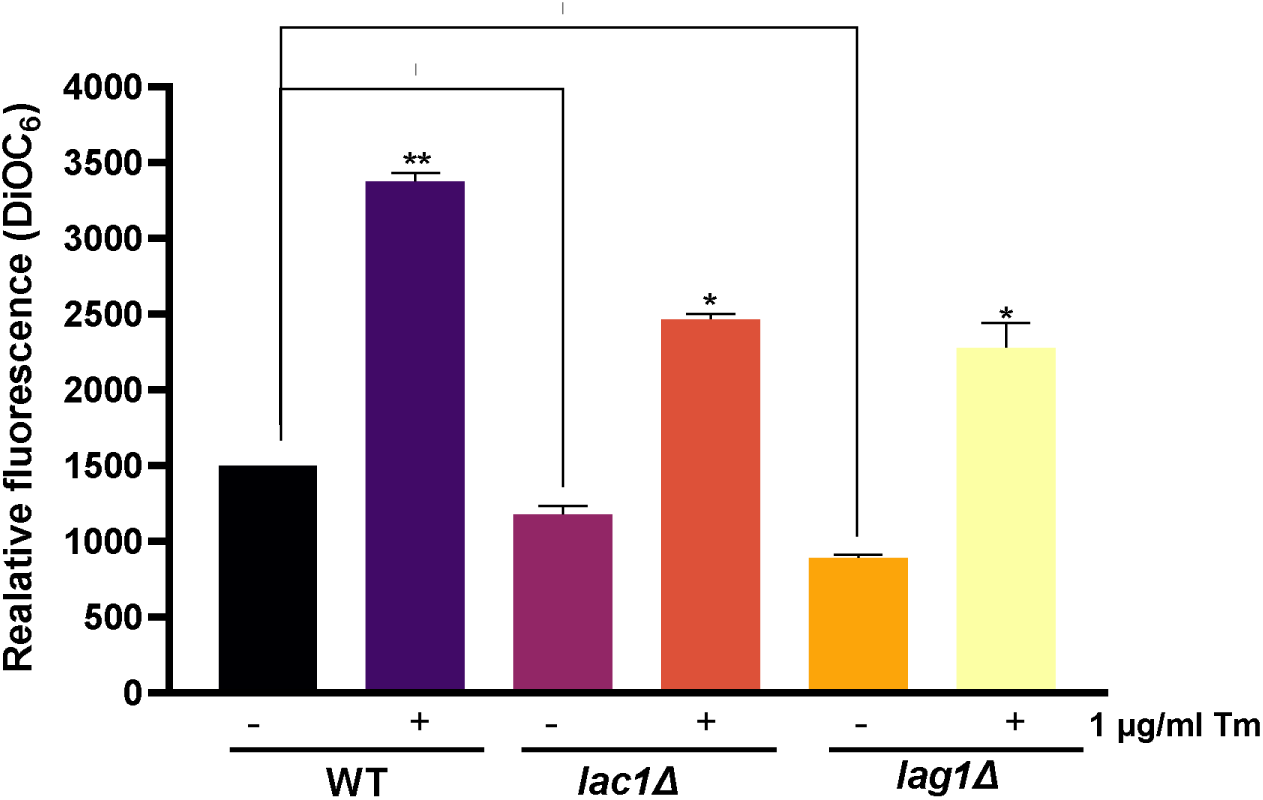
Deletion of LAG1 significantly reduces phospholipid levels and modifies membrane morphology. (A) Cells were grown in SC media with or without tunicamycin until they reached the mid-log phase. Equal numbers of cells were then harvested for lipid extraction. Thin-layer chromatography (TLC) was performed to separate phospholipids (PL). (B) Phosphorus quantification was conducted to measure PL levels, and plotted (C) mRNA was extracted and subjected to qRT PCR (D). Membrane morphology was examined using DiOC_6_ staining. Images were captured using a Zeiss LSM 710 laser-scanning confocal microscope, with a scale bar representing 10 microns and excitation/emission wavelengths of 482/504 nm. The fluorescence intensity of DiOC_6_ was recorded using a Hitachi F-4500 fluorescence spectrometer at excitation/emission wavelengths of 482/504 nm, and the results were plotted. Data represent the mean□±□SD from three independent experiments (*P□<□0.05).

Gene expression analysis of key enzymes involved in the de novo synthesis of PC and PE was also conducted. The decarboxylases *PSD1* and *PSD2* convert phosphatidylserine (PS) to PE, while *CHO2* and *OPI3* methyltransferases convert PE to PC[26]. In *lac1*Δ cells, expression of these PL-synthesizing genes was slightly reduced, except for *PSD1*, which maintained expression levels similar to WT (Fig. 4C). By comparison, *lag1*Δ cells demonstrated a marked reduction in the expression of all PL synthesizing genes relative to both WT and *lac1*Δ strains. When exposed to ER stress, *lag1*Δ cells exhibited an increase in *PSD1* and *PSD2* comparable to ER stress-induced *LAC1* deletion cells, though the *CHO2* and *OPI3* were still significantly less than that observed in WT and *lac1*Δ cells.

To further support these findings, membrane morphology was assessed using DiOC_6_ staining. In *lac1*Δ cells, fluorescence signals were reduced, indicating potential alterations in membrane structure (Fig. 4D). In contrast, *lag1*Δ cells exhibited a marked decrease in fluorescence intensity, suggesting more pronounced structural changes in the membrane. Under ER stress, DiOC_6_ fluorescence in *lag1*Δ cells remained relatively low compared to tunicamycin-treated WT cells but was similar to that observed in tunicamycin-treated *lac1*Δ cells.

In summary, *LAC1* deletion significantly reduces PC and PE levels, whereas *LAG1 deletion* has a considerably greater impact. Both deletions affect the expression of phospholipid-synthesizing genes, with *LAG1* deletion showing a more severe effect. The pronounced changes in membrane morphology in *lag1*Δ cells highlight the essential role of *LAG1* in preserving phospholipid composition and membrane integrity.

#### 3.5. Deletion of *LAG1* increased the neutral lipid levels, expression of its synthesizing genes, and lipid droplets

Lipid droplets (LDs) serve as storage depots for triacylglycerols (TAG) and sterol esters (SE), which are collectively known as neutral lipids (NLs)[27]. These LDs can be broken down to meet the energy demands of the cell[28]. Under conditions of ER stress, an increase in the accumulation of storage lipids and a rise in the number of LDs have been observed[29]. The role of neutral lipids (NL) in cells lacking *LAG1* and *LAC1* was studied. It was found that *lag1*Δ cells exhibited a marked increase in TAG and SE levels compared to *lac1*Δ and WT cells (Fig. 5A and B).

**Figure 5.**
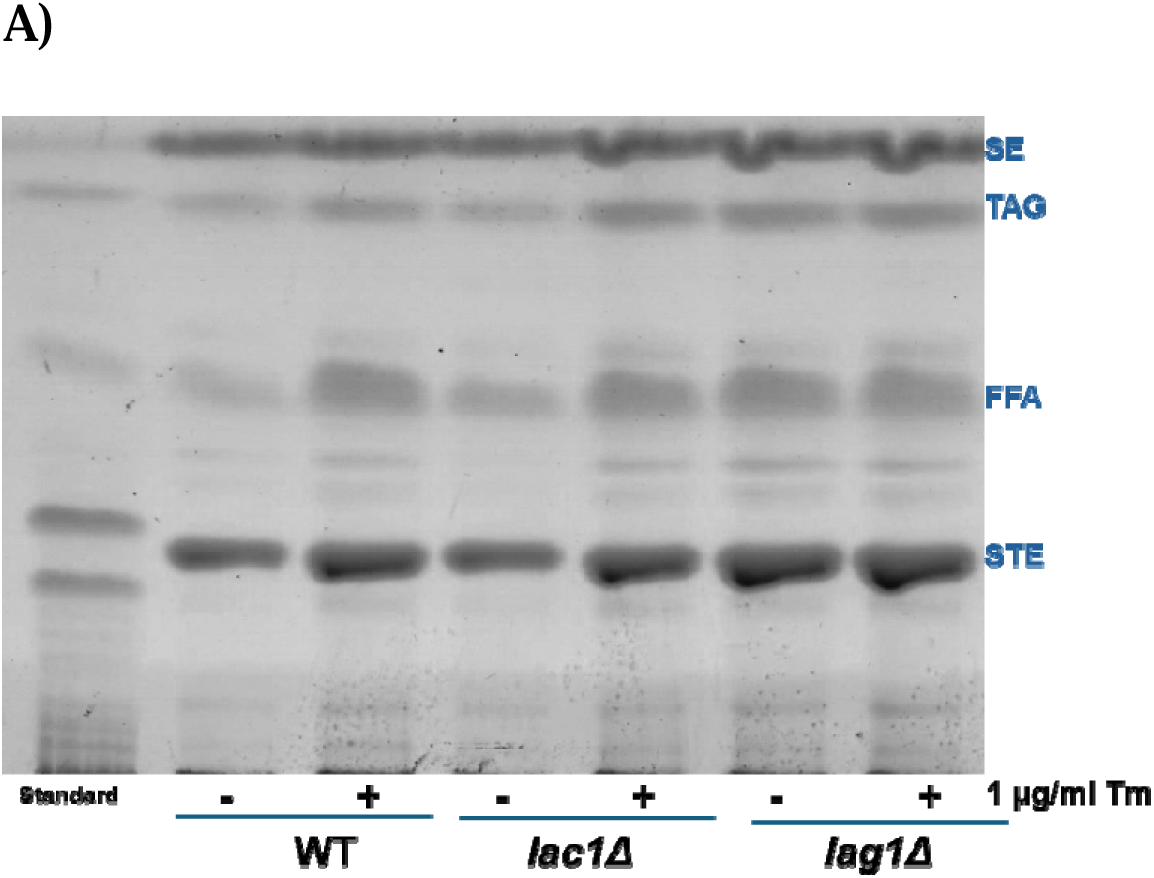
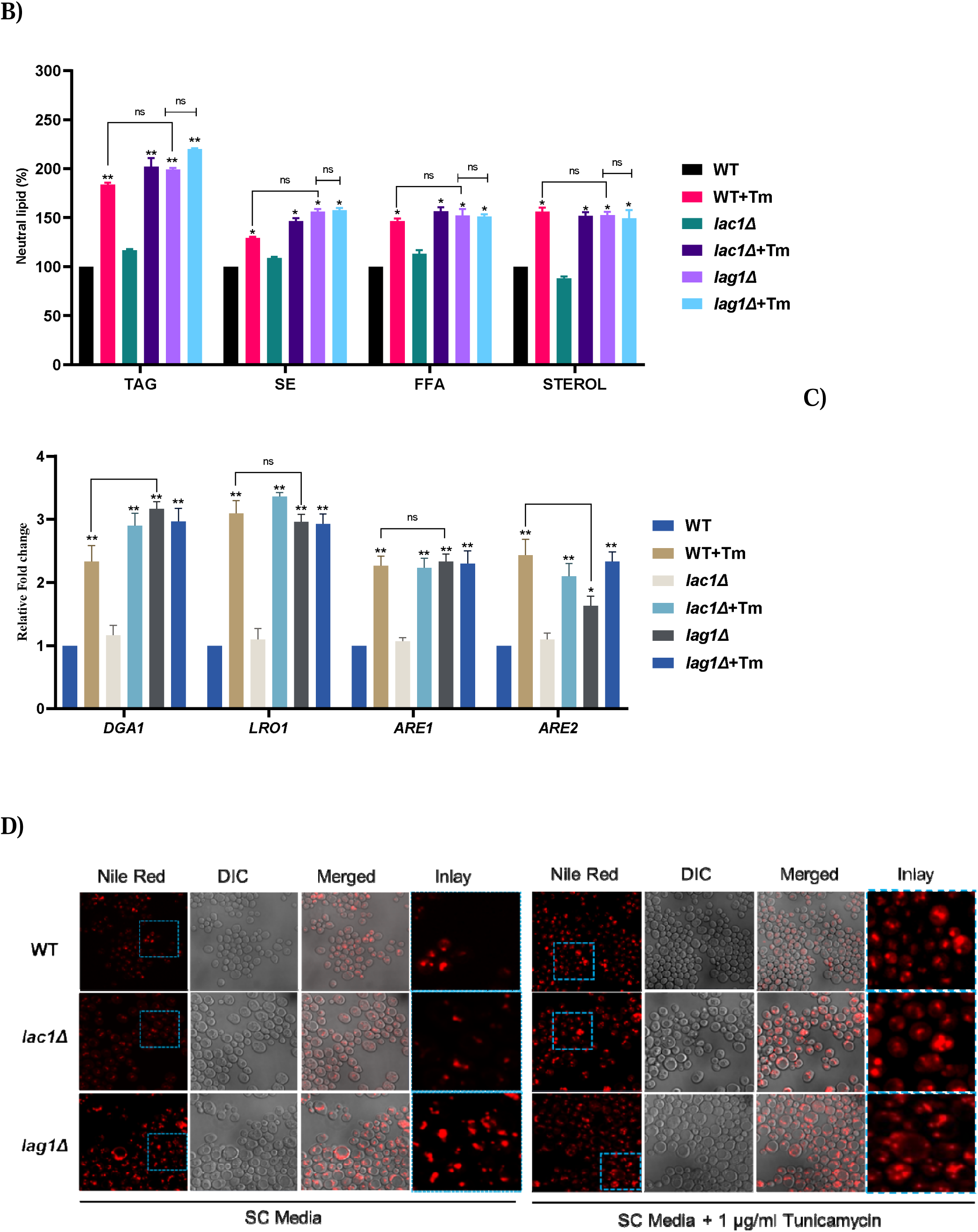
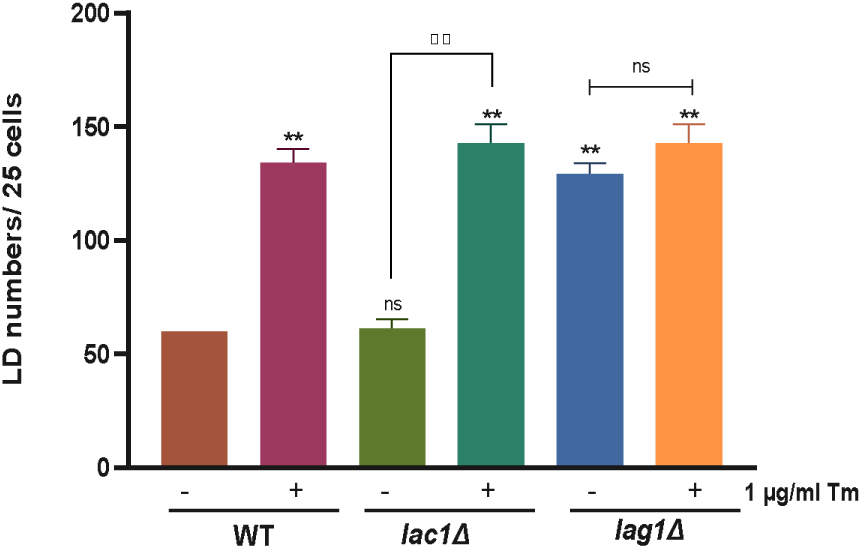
Deletion of LAG1 modifies neutral lipid homeostasis. (A) Neutral lipids (NL) were extracted from cells using protocols described in the methods section, and individual NLs were separated by TLC. (B) The separated NLs were quantified by measuring band intensities with ImageJ software, and the results were plotted. (C) qRT PCR was performed to analyze the expression levels of genes involved in NL synthesis. (D) Lipid droplets (LD) were stained with Nile red, and images were captured using a confocal fluorescence microscope with a 100×/1.40 oil immersion objective. Excitation and emission wavelengths were set at 480 and 510 nm, respectively. LDs were counted in 25 randomly selected cells using ImageJ software, and the results were presented as a graph. Data represent the mean□±□SD from three independent experiments (*P□<□0.05).

TAG biosynthesis involves converting diacylglycerol (DAG) to TAG through the action of two acyltransferases, *DGA1* and *LRO1*. Similarly, the production of SE from sterols is facilitated by the key enzymes *ARE1* and *ARE2*[30]. The findings showed a significant increase in the expression of *LRO1, DGA1, ARE1,* and *ARE2* in *lag1*Δ cells (Fig. 5C). Additionally, staining with Nile red revealed an accumulation of LDs in *lag1*Δ cells compared to *lac1*Δ and WT cells (Fig. 5D).

Interestingly, both *lac1*Δ and WT cells demonstrated increased synthesis of neutral lipids, their synthesizing genes, and higher LD counts during ER stress conditions. In contrast, *lag1*Δ cells showed no change in neutral lipid levels and its synthesizing gene levels and LD numbers during ER stress (Fig. 5). This indicates that the deletion of *LAG1* affects the synthesis of neutral lipids, leading to an increase in LDs. However, this increase does not further escalate under ER stress conditions in *lag1*Δ cells.

### The protective effect conferred by *LAG1* deletion against ER stress relies on PL synthesis and activation of the ER stress response rather than on the synthesis of NL

#### 3.6. A functional ER stress response is required for the protective effects of *LAG1* deletion against ER stress

Although *lac1*Δ altered phospholipid (PL) and neutral lipid (NL) biosynthesis, it did not confer any protective advantage against ER stress. Consequently, further analysis was centered on *lag1*Δ cells. To investigate the protective effects of *LAG1* deletion against ER stress more thoroughly, double deletion strains (*ire1*Δ*lag1*Δ*, hac1*Δ*lag1*Δ, and *rpn4*Δ*lag1*Δ) were generated to assess their growth under both normal and ER stress conditions. The growth patterns of these *LAG1* double deletion strains mirrored those of the corresponding single ER stress response gene deletions, indicating that *LAG1* deletion does not compensate for the loss of ER stress response genes in promoting cell growth (Fig. 6A). This finding suggests that the protective effect of *LAG1* deletion is dependent on an intact ER stress response pathway.

**Figure 6.**
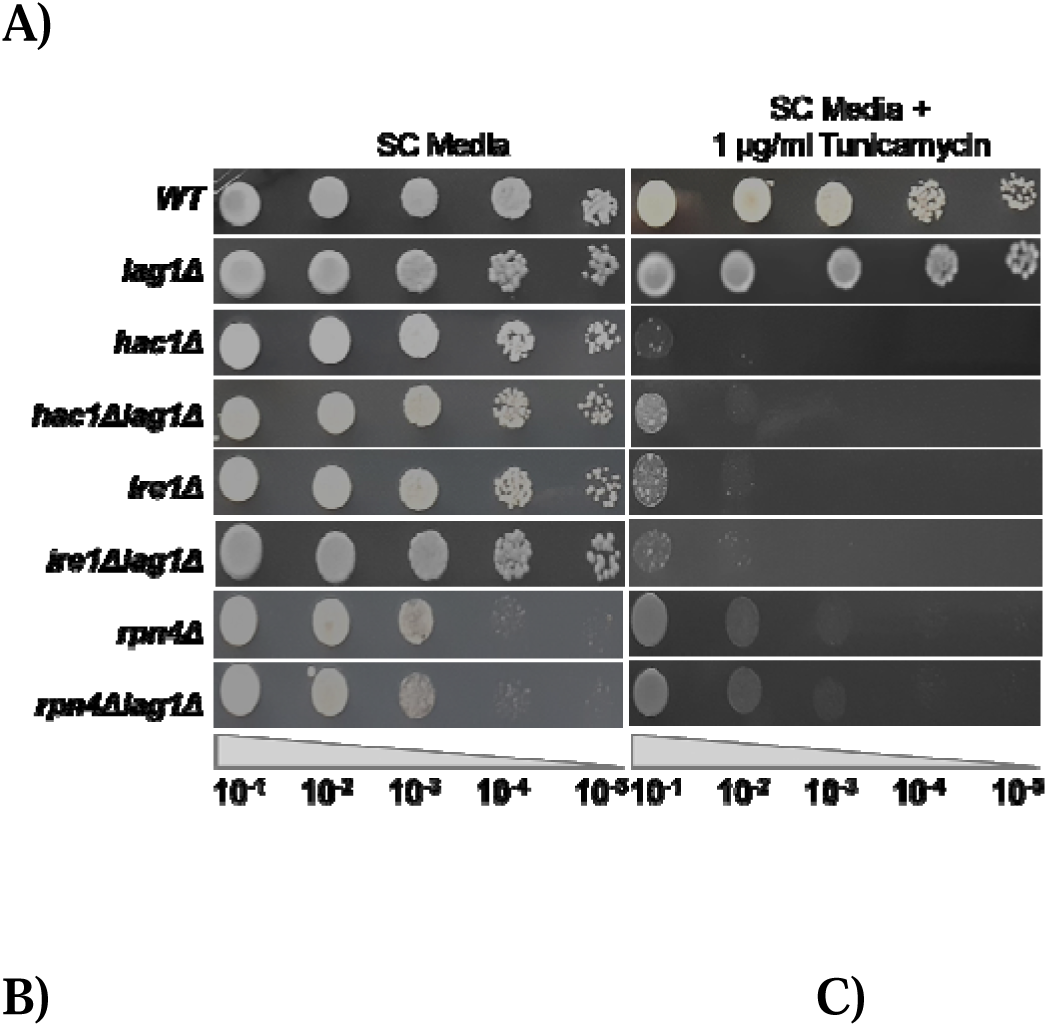
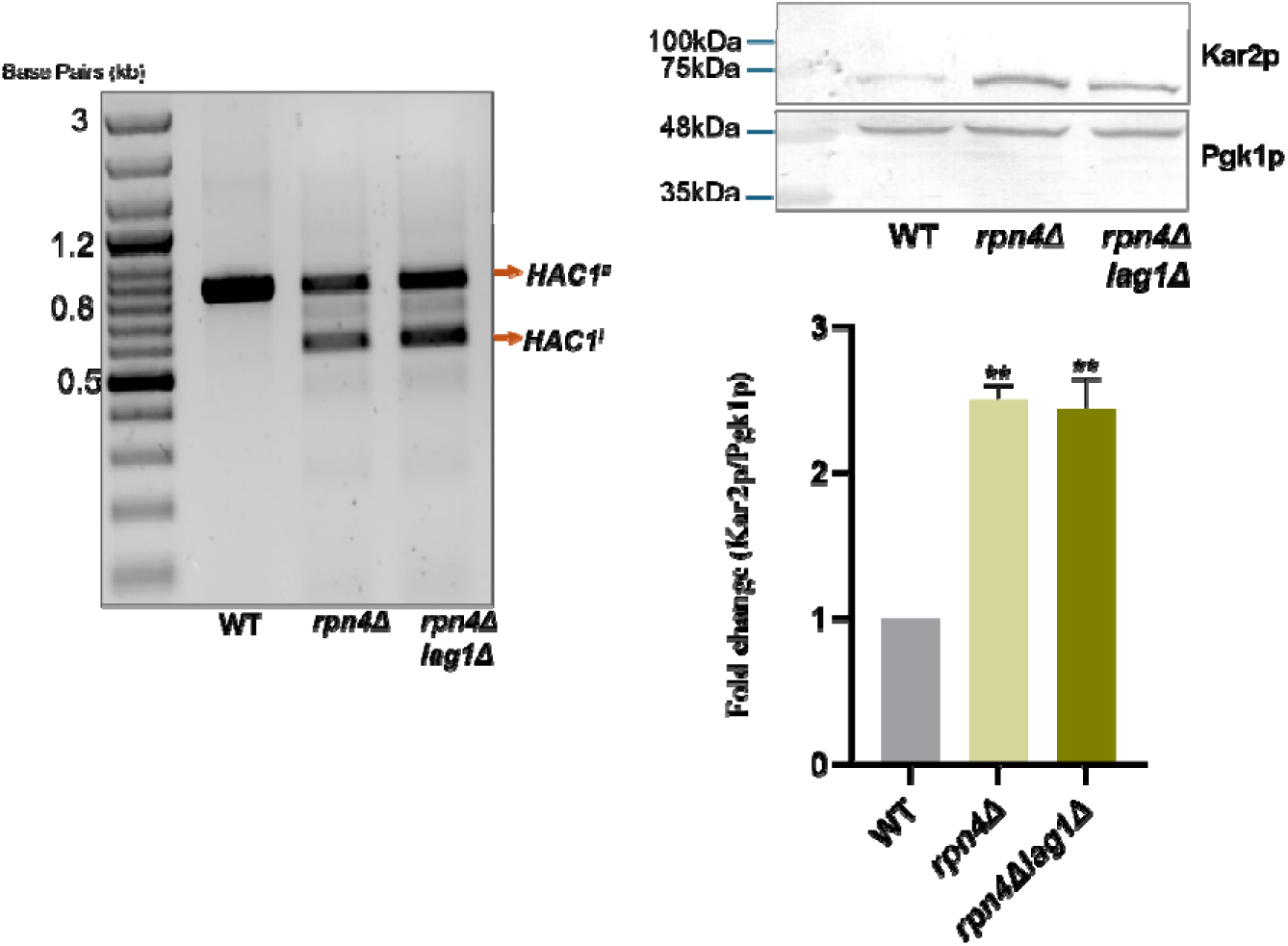
The deletion of LAG1, alongside ER stress response genes, fails to rescue cell growth. (A) A spot growth experiment was conducted to assess cell growth. Cells were cultivated in SC medium at 30° C, and cells with an identical optical density (3.0 OD at A_600 nm_) were subjected to serial dilution (10-fold). These diluted cells were then spotted onto agar plates containing SC medium, with or without tunicamycin, and incubated as prescribed. B) Using the Superscript II reverse transcriptase, we studied the *HAC1* mRNA splicing. A picture of the 1% agarose gel was taken after the acquired PCR products C) Equal amounts of protein from each cell were extracted and put onto a 12% SDS-PAGE gel. Pgk1p was the loading control, while anti-Kar2p was the primary antibody. The data represents the mean ± SD (*P<0.05) from three separate experiments.

As outlined in Section 3.3, *HAC1* splicing and Kar2p expression are absent in *HAC1* and *IRE1* deletion strains[15]. To evaluate these responses, *rpn4*Δ*lag1*Δ cells were analyzed alongside appropriate controls for *HAC1* splicing and Kar2p expression, revealing that *HAC1* splicing remained unchanged while Kar2p expression was elevated (Fig.6B and C). These results underscore that an intact ER stress response pathway is essential for the ER stress mitigation associated with *LAG1* deletion.

#### 3.7. A proper phospholipid synthesis is essential for the ER stress resilience observed in the *LAG1* deletion condition

Phospholipid homeostasis is essential for the normal functioning of subcellular organelles, particularly the ER[31]. Imbalances in PC and PE levels can induce lipid bilayer stress[32], significantly affecting the integrity and function of the ER membrane. Beyond its role in protein quality control, the UPR pathway also transcriptionally regulates phospholipid biosynthesis, thus playing a critical role in mitigating ER stress resulting from lipid dysregulation[21], [31]. Deleting genes involved in phospholipid production, such as *CHO2* or *PSD2*, in *IRE1* mutants and its downstream transcription factor *HAC1* can be lethal to cells[33], [34]. Consequently, the UPR and proper phospholipid biosynthesis are vital for maintaining the integrity of subcellular membrane structures and proteostasis. Studies have indicated that cell growth is impaired under ER stress conditions in cells with defective PC and PE synthesis[35], [36]. In *LAG1* deletion cells, we observed decreased phospholipid synthesis and abnormal membrane structures (Fig. 4). despite these alterations, *LAG1* deletion also demonstrated a protective role against ER stress, as shown in Figures 1 and 2.

To further explore the relationship among phospholipid biosynthesis, ER stress, and *LAG1*, we intentionally disrupted PC and PE synthesis by deleting their respective biosynthetic genes in *LAG1* deletion cells (*cho2*Δ*lag1*Δ*, opi3*Δ*lag1*Δ*, psd1*Δ*lag1*Δ, and *psd2*Δ*lag1*Δ). The objective was to determine whether *LAG1* deletion could alleviate the growth defects caused by impaired phospholipid synthesis, both in the presence and absence of ER stress. The results indicated that cell growth was significantly compromised in cells with double deletions of the PC and PE genes alongside LAG1, resembling the growth defects observed in cells lacking only the PC and PE genes under ER stress (Fig. 7A and B).

**Figure 7.**
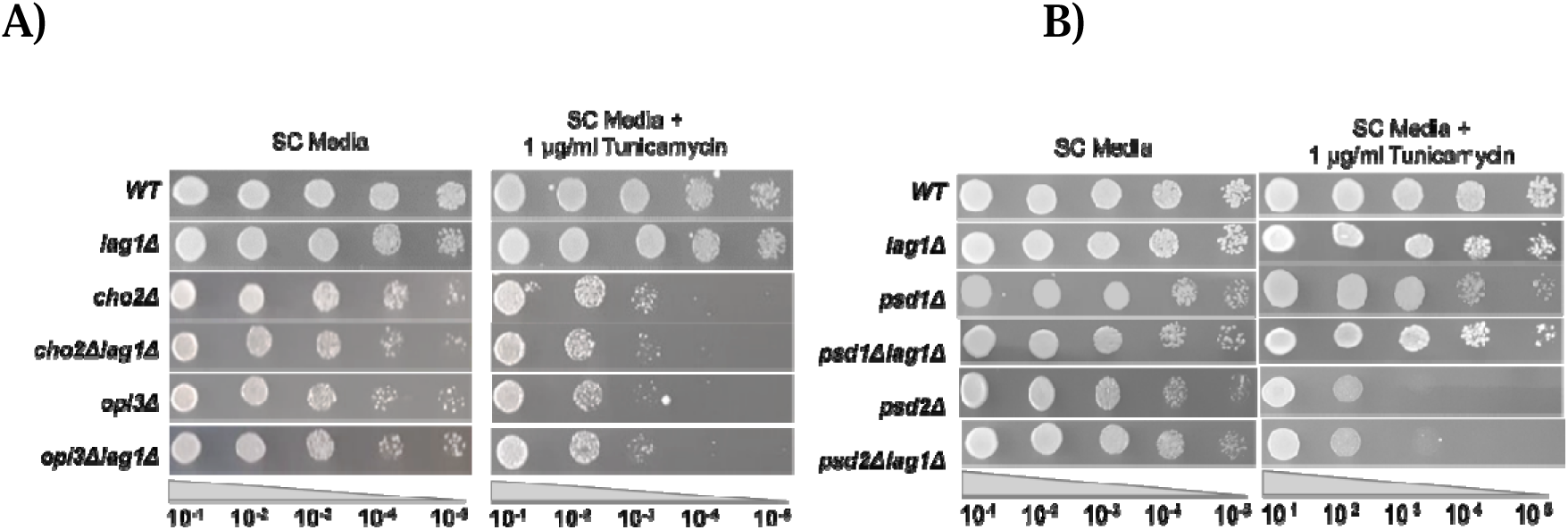
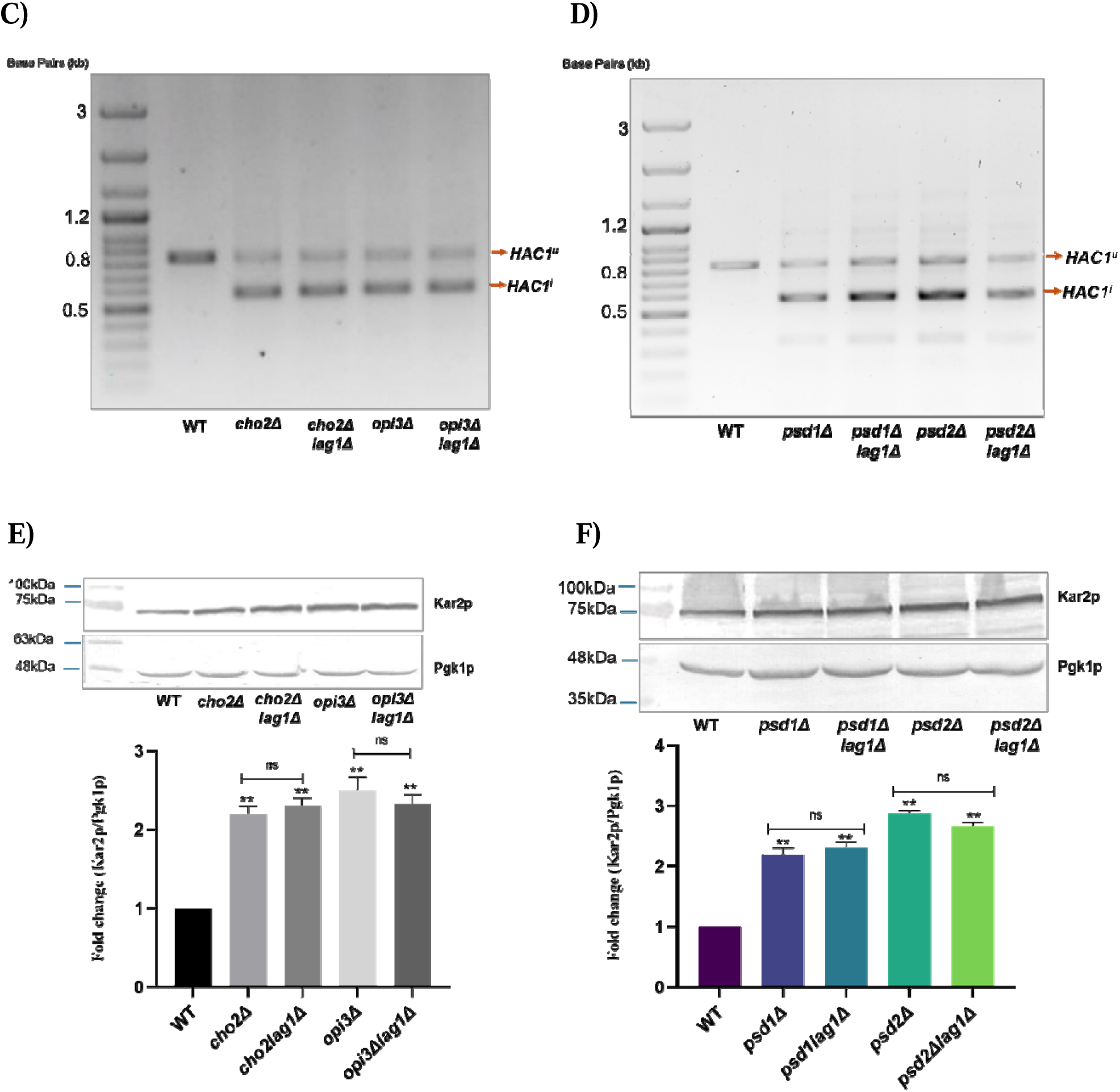
Functional phospholipid biosynthesis is essential for the ER stress-buffering function noted in *LAG1* deletion cells. A and B) A spot growth experiment was conducted to evaluate cell growth. C and D) The *HAC1* mRNA splicing experiment was performed on 1% agarose gel. E and F) An equal amount of protein was extracted from each cell and loaded onto a 12% SDS-PAGE gel. Pgk1p served as the loading control, while the primary antibody was anti-Kar2p. The data represents the mean ± SD (*P<0.05) from three separate experiments

Among the cells lacking PC and PE genes in addition to *LAG1* deletion, *HAC1* splicing was detected (Fig. 7C and D), and the expression of the ER chaperone Kar2p was elevated compared to the WT (Fig. 7E and F). These findings suggest that the ER stress-mitigating function of LAG1 is contingent upon functional phospholipid biosynthesis.

#### 3.8. Neutral lipids are not essential for *LAG1* deletion-mediated cell survival under ER stress

In contrast to phospholipids, prior studies have suggested that neutral lipids are not crucial for maintaining the integrity of the ER and ensuring protein quality control[29], [33]. Notably, increased synthesis of TAGs, SE, and LDs was explicitly observed in *lag1*Δ cells compared to *lac1*Δ and WT cells. To further explore the role of TAGs and SEs in the ER stress-alleviating function of *lag1*Δ cells, double deletion strains were generated: *lro1*Δ*lag1*Δ and *dga1*Δ*lag1*Δ (lacking TAG-synthesizing genes) as well as *are1*Δ*lag1*Δ and *are2*Δ*lag1*Δ (lacking SE-synthesizing genes).

Interestingly, data indicated that the growth of these double deletion strains (*dga1*Δ*lag1*Δ*, lro1*Δ*lag1*Δ*, are1*Δ*lag1*Δ, and *are2*Δ*lag1*Δ) was not significantly altered, exhibiting growth patterns comparable to those of WT cells under both ER stress and non-stress conditions (Fig. 8A and B). Additionally, UPR activation assays and analyses of Kar2 protein expression demonstrated no *HAC1* splicing, indicating a lack of UPR activation (Fig. 8C and D). In contrast, Kar2p expression remained normal in all double deletion strains (Fig. 8E and F). In conclusion, the findings demonstrate that neutral lipid synthesis is not essential for the ER stress protective function of *lag1*Δ cells.

**Figure 8.**
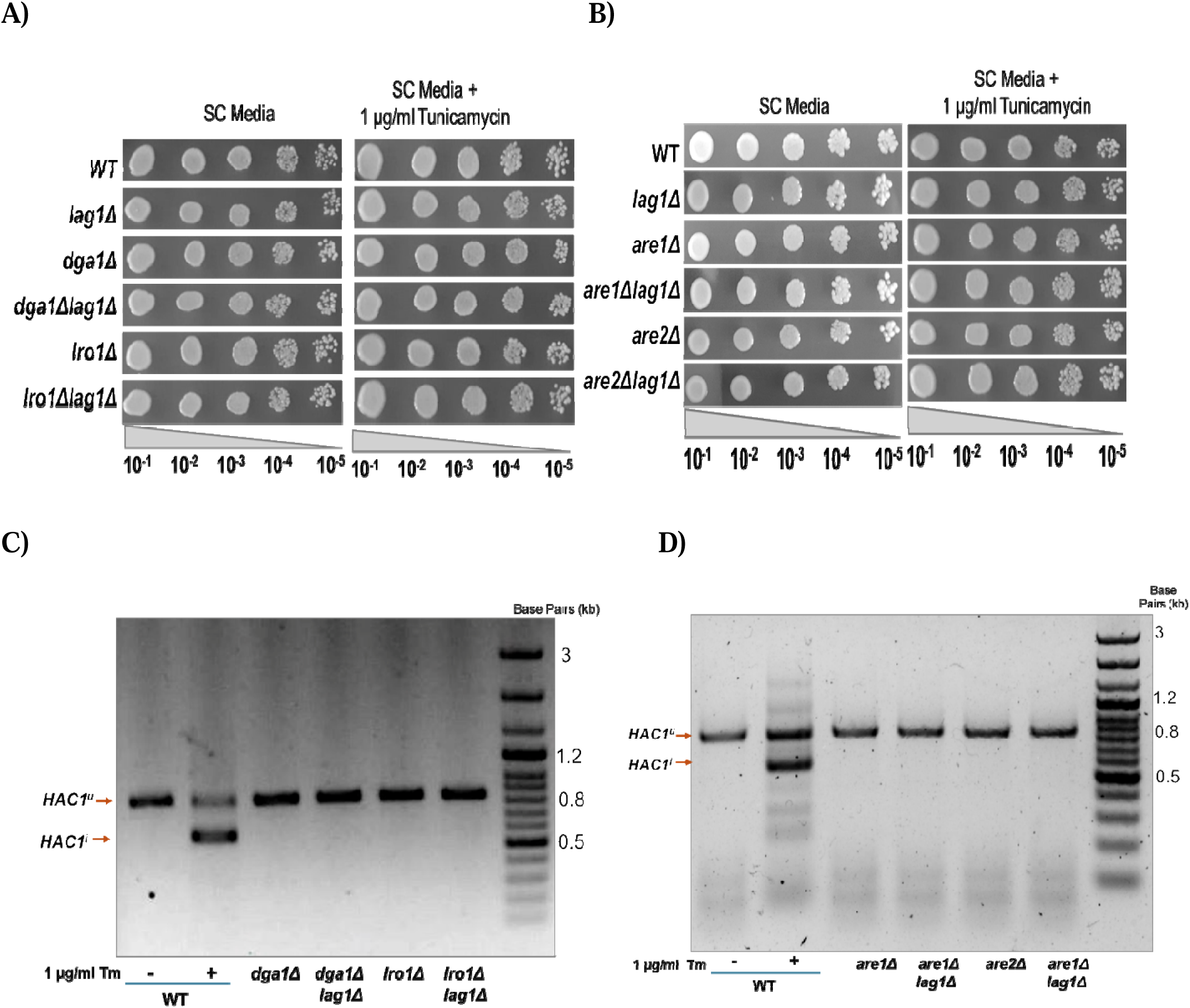
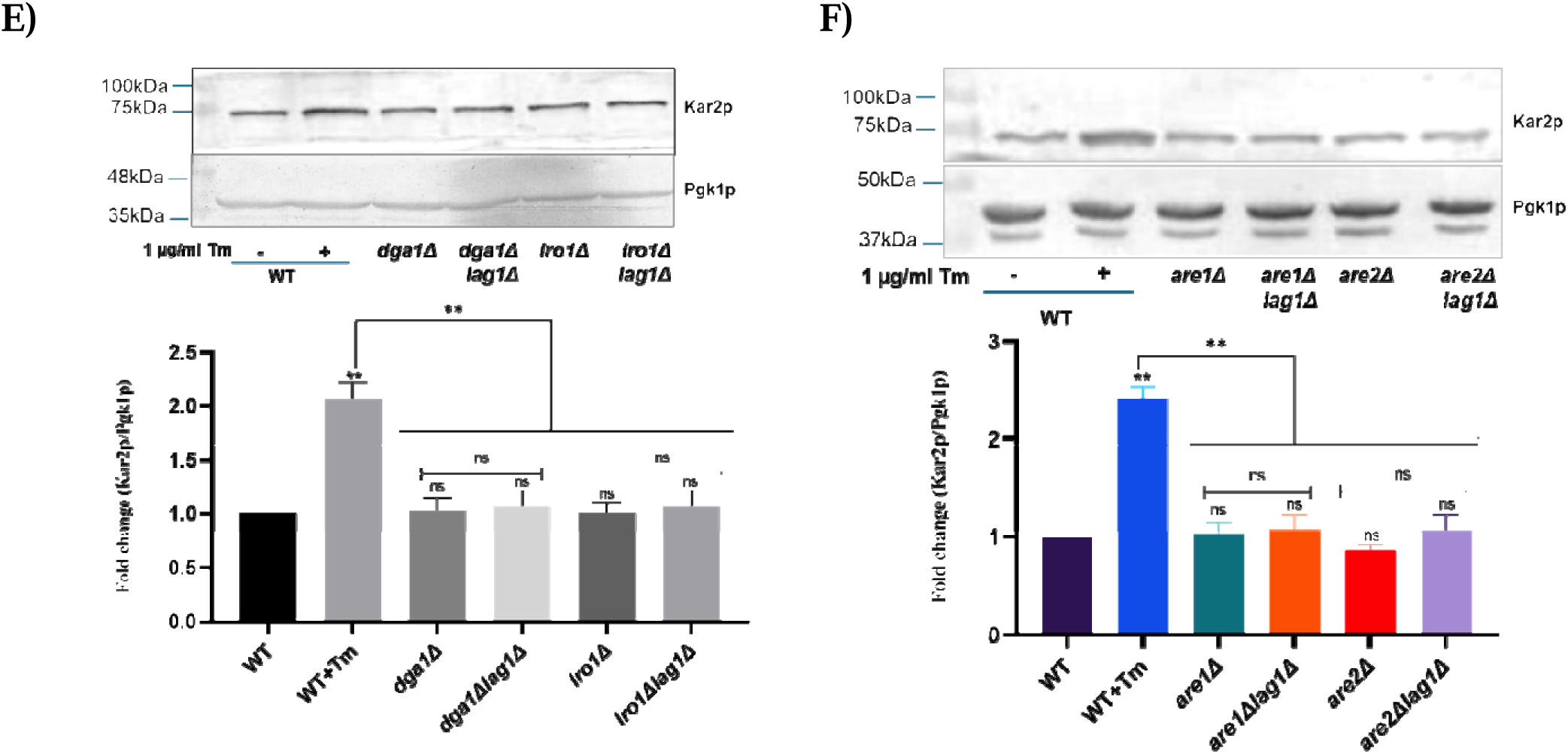
Cell survival under ER stress is not reliant on the NL content in *LAG1* deletion cells. (A and B) A spot growth experiment was conducted following the previously described procedure to assess cell growth. (C and D) The *HAC1* mRNA splicing experiment was carried out, and the resulting PCR products were separated on a 1% agarose gel. The data obtained from the gel electrophoresis was recorded. (E and F) A 12% SDS-PAGE gel was loaded with equal protein concentrations. Anti-Kar2p was the primary antibody employed, while Pgk1p was the loading control. The data represent the means ± SD (*P<0.05) from three experiments.

### *LAG1* overexpression exacerbated ER stress compared to *LAC1* overexpression, and it also altered PL and NL biosynthesis

#### 3.9. Overexpression of LAG1 in wild-type cells showed severe growth defects

*LAC1* and *LAG1* were overexpressed in WT cells using the pYES2 high-copy-number galactose-inducible yeast episomal vector[37]. Analysis of cell growth revealed that the overexpression of *LAG1* (WT+*LAG1*) significantly impaired cell growth in both solid and liquid media (Fig. 9A and B), with this growth reduction exacerbated under ER stress conditions (Fig. 9B). In contrast, overexpression of *LAC1* (WT+*LAC1*) resulted in only a slight decrease in growth in solid media and a significant reduction in liquid media under normal conditions (Fig.9A and B). However, the impact of WT+*LAC1* during ER stress was considerably lesser than that observed with WT+*LAG1* (Fig. 9A and B). These results highlight the more pronounced negative effect of WT+*LAG1* on cellular growth compared to WT+*LAC1* and the vector control (WT-Vec).

**Figure 9.**
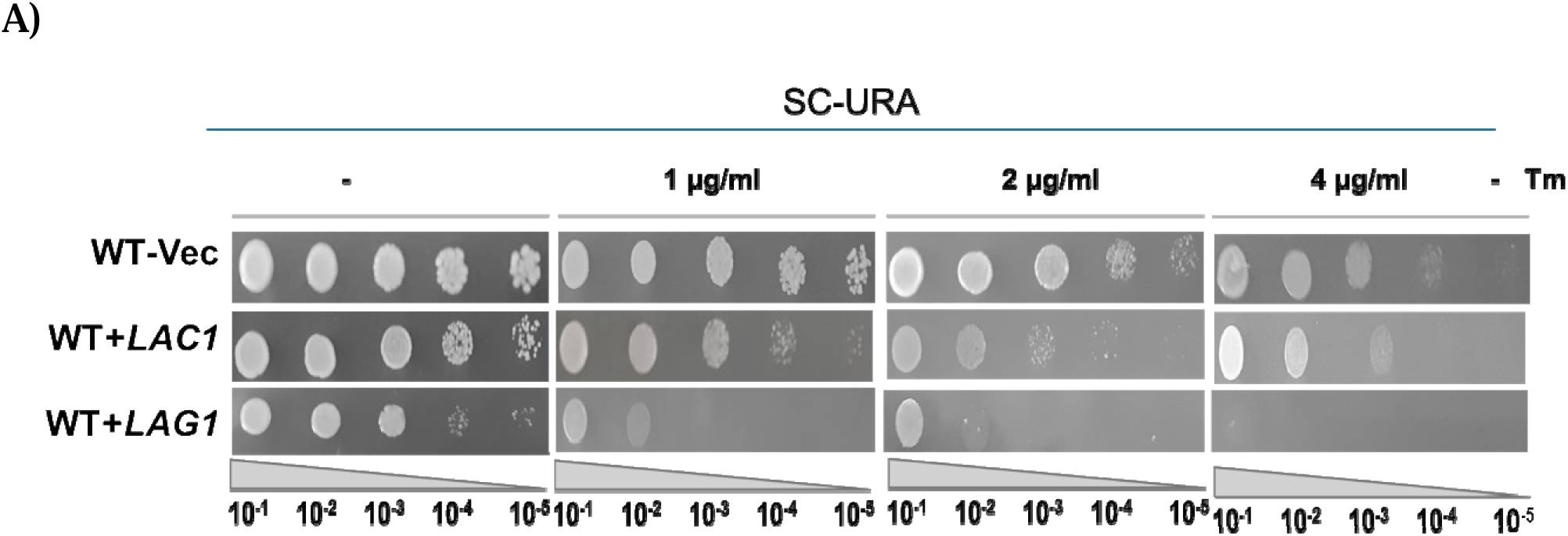
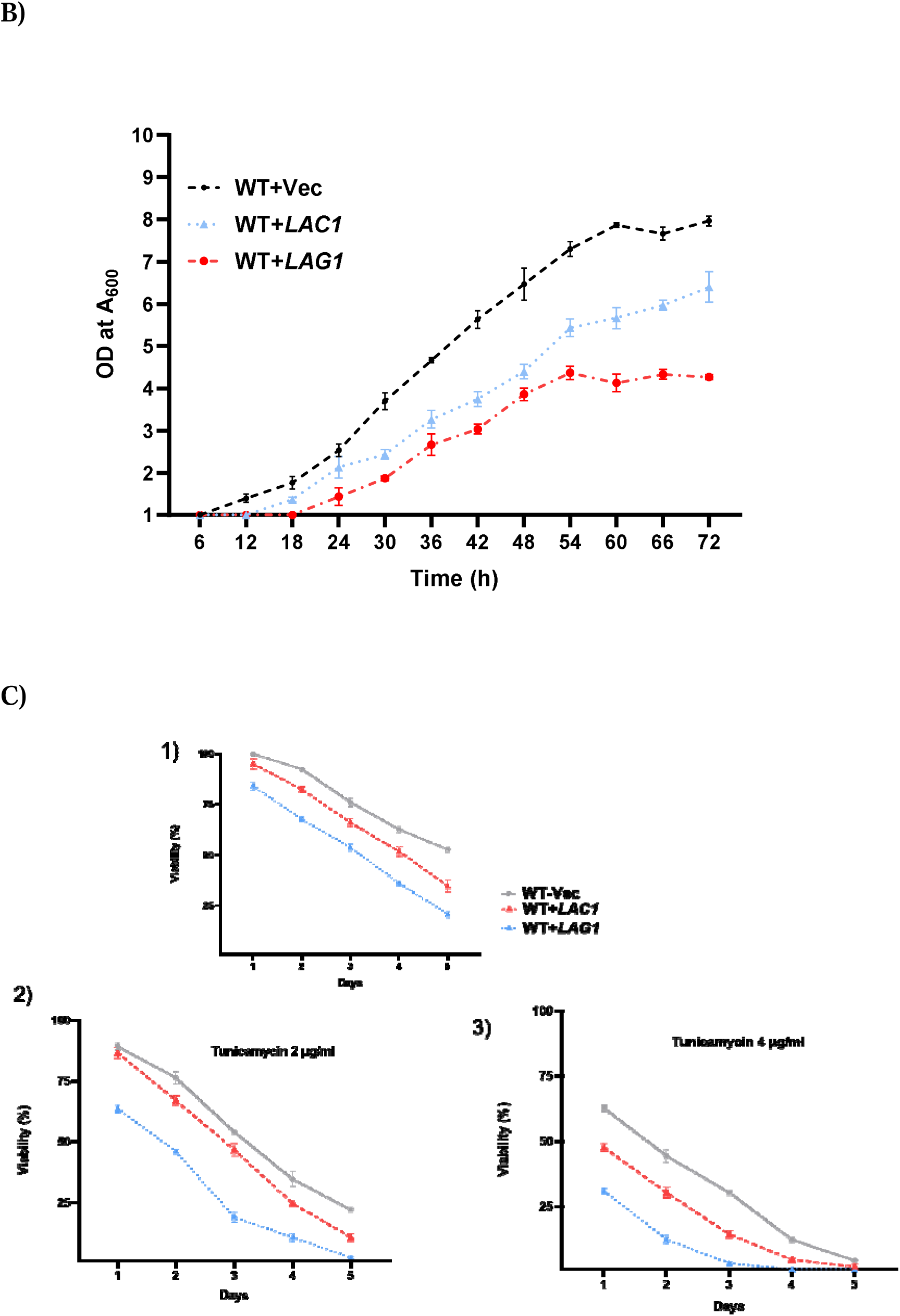
Wild-type cells overexpressing *LAG1* displayed significant growth impairment and reduced cell survival during ER stress. (A) The strains overexpressing the target genes and the vector control were cultured in SC-Ura (SC media devoid of uracil) + 2% galactose. The OD at 600 nm was measured regularly over 72 hours to monitor cell growth. (B) In a separate experiment, the overexpression strains and vector control were initially grown in SC media at 30°C till the mid-log phase. After harvesting the cells, an equal OD (3.0 at A600_nm_) was used to prepare serial dilutions (10-fold). These diluted cells were then spotted onto SC-Ura media supplemented with 2% galactose, with or without Tunicamycin. The agar plates were incubated for two days at 30° C, and images of the plates were captured. (C) *LAG1* and *LAC1* overexpressed cells were cultured in SC-Ura media containing 2% galactose. Afterward, the cells were collected at specific intervals ranging from 1 to 5 days. Using a haematocytometer, an equal number of cells (1000 cells) were counted and subsequently plated on SC-Ura media supplemented with 2% galactose, with or without Tunicamycin. The plating process was conducted under sterile conditions using an autoclaved L-rod. The plated cells were allowed to grow for 48 hours at 30 °C. Cell viability of cells overexpressing *LAC1* and *LAG1* was determined by comparing their CFUs to the CFU count on day 1 of the WT vector control cells. Viability was expressed as a percentage, with 100% representing the CFU count on day 1 of WT-Vec. The data represent the mean ± SD (*P<0.05) from three separate experiments.

A cell survival assay further demonstrated a dramatic decrease in viability among cells overexpressing *LAG1* (Fig. 9C.1). The viability of WT+*LAG1* cells was further diminished under higher concentrations of tunicamycin (2 and 4 µg/ml) when compared to cells with *LAC1* overexpression and the vector control (Fig. 9C.2 and C.3). Although *LAC1* overexpression did result in a significant reduction in cell viability, it was less impactful than the decline observed in *LAG1*-overexpressing cells. Overall, these results demonstrate that overexpression of *LAG1* exerts more detrimental effects on cell growth and viability than *LAC1* overexpression, emphasizing the similar yet distinct roles of these two genes in cellular stress responses.

#### 3.10. Overexpression of *LAG1* triggered a pronounced ER stress response

Overexpression of *LAG1* markedly intensified the ER stress response, as evidenced by enhanced expression of ER stress response genes, *HAC1* splicing increased Kar2p protein levels, and robust UPRE-GFP fluorescence signals (Fig. 10A to D). This indicates a heightened ER stress induction in WT+*LAG1* cells compared to WT+*LAC1*. While *LAC1* overexpression also triggered ER stress, the ER stress response genes, and Kar2p upregulation levels were significantly lower in *LAG1*-overexpressing cells.

**Figure 10.**
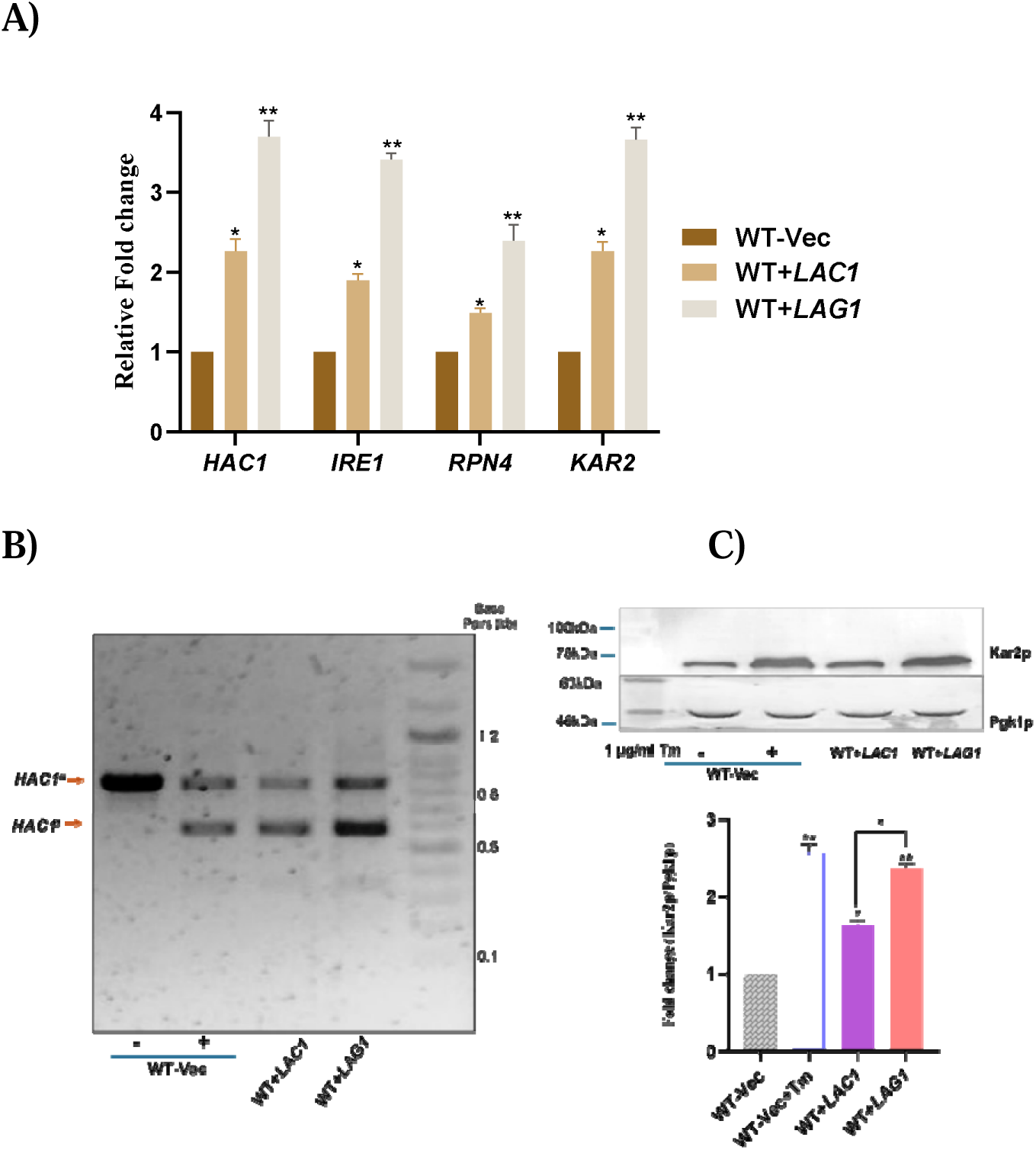
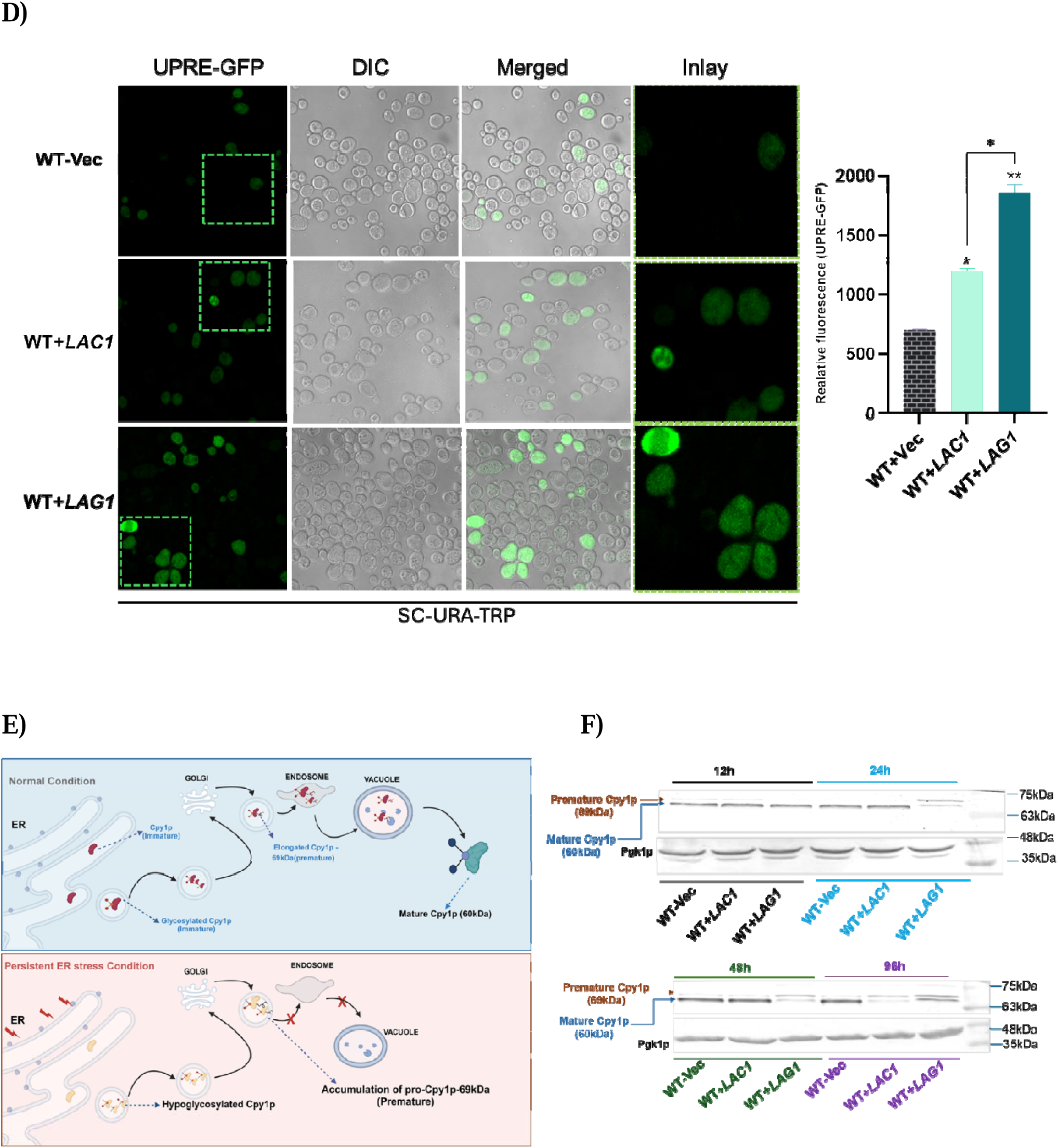
Overexpression of *LAG1* in wild-type cells induced the ER stress response. (A) Q-RT PCR was used to quantify the gene expression. (B) 1% agarose gel was used to separate the PCR products produced by the HAC1 mRNA splicing experiment. Records were made of the gel electrophoresis data. (C) A 12% SDS-PAGE gel was loaded with identical protein concentrations. The primary antibody used was anti-Kar2, while Pgk1p was the loading control. (D) The pRS314 vector expressing UPRE-GFP was used to transform yeast cells, and then the cells were cultured till they reached the mid-log phase on SC-Trp-Ura media with the transformed cells; the expression of UPRE-GFP was measured with an F-4500 fluorescence spectrophotometer. Fluorescence measurements were recorded and plotted at 510 nm for the emission and 488 nm for the excitation. (E) Graphical illustration of Cpy1p maturation (F) The target genes overexpressed strains and the wild-type cells transformed with vector control were cultivated in SC-Ura + 2% galactose medium. The cells were collected at specified time points as indicated. Protein was extracted in equal quantities from the cells and then put onto a 10% SDS-PAGE gel. Anti-Cpy1p was the primary antibody utilized for this study, while Pgk1p was the loading control. The data represents the mean ± SD (*P<0.05) from three separate experiments.

To further evaluate and compare ER stress levels between *LAC1* and *LAG1* overexpression, cells were treated with Aureobasidin A (AbA), an inhibitor of inositol phosphoryl ceramide synthase *AUR1* [38], [39](Fig. 13A). By inhibiting the conversion of ceramides to complex ceramides, AbA treatment induces ceramide accumulation, leading to cell death (Fig. S3A). Notably, ceramide synthesis-deficient strains, such as the *lac1*Δ *lag1*Δ double mutants, could grow even at lethal AbA doses due to their inability to accumulate ceramides[38]. Overexpression of *LAG1* led to a marked reduction in growth, while *LAC1* overexpression caused a more moderate growth impairment, underscoring the higher ER stress burden from *LAG1* overexpression (Fig. S3B and C). Additionally, *HAC1* splicing was observed with AbA treatment to vector control itself (Fig. S3D), and Kar2p expression levels were more elevated in *LAG1*-overexpressing cells treated with AbA compared to *LAC1*-overexpressing cells under the same conditions (Fig. S3E).

To track unfolded protein response element (UPRE) gene activation in response to ER stress, cells overexpressing *LAG1* or *LAC1* and a control vector were transformed with a GFP-tagged UPRE plasmid (pRS314), allowing GFP fluorescence to act as an indicator of UPRE activation. UPRE induction was notably more pronounced in *LAG1*-overexpressing cells than those overexpressing *LAC1*, though both showed greater UPRE activation than WT-Vec (Fig.10D).

Carboxypeptidase Y (CPY), an enzyme predominantly found in the vacuole, plays a key role in removing amino acids from the carboxy-terminal of proteins. CPY is initially synthesized in an inactive, N-glycosylated form within the ER lumen during its maturation. It is then transported to the Golgi apparatus, where further modifications yield a premature form, pro-CPY, with a molecular weight of 69 kDa. Pro-CPY is subsequently directed to the vacuole via late endosomes, where vacuolar peptidase B cleaves its polypeptide chain, activating CPY into its mature 61 kDa form. However, persistent ER stress can lead to CPY hypo-glycosylation and accumulation of the precursor form [40](Fig. 10E). In cells overexpressing *LAG1*, the accumulation of premature CPY was observed after 24 hours and persisted through 96 hours. In contrast, *LAC1*-overexpressing cells exhibited accumulation of premature CPY only at the 96-hour mark (Fig. 10F). These findings indicate that *LAG1* overexpression severely disrupts ER homeostasis, impairing CPY maturation and eliciting a more robust ER stress response than *LAC1* overexpression.

#### 3.11. *LAG1* overexpression elevates phospholipid biosynthesis and alters membrane morphology

Phospholipid (PL) levels and the expression of their biosynthetic genes were elevated in WT+*LAG1* and WT+*LAC1* cells compared to the vector control (Fig. 11A and B). However, the increase was particularly pronounced in WT+*LAG1* cells, which exhibited a robust elevation in overall PL content and the expression of key genes involved in PL biosynthesis, including *CHO2, OPI3*, and *PSD2*. This suggests that *LAG1* may play a stronger role in promoting PL metabolism than *LAC1*. Furthermore, *LAG1* overexpression led to a marked increase in abnormal membrane structures, an effect that was more severe than that observed with *LAC1* overexpression and the vector control, as highlighted by the yellow arrow (Fig. 11D). This proliferation of atypical membrane formations underscores the structural impact of WT+*LAC1* on cellular membranes.

**Figure 11.**
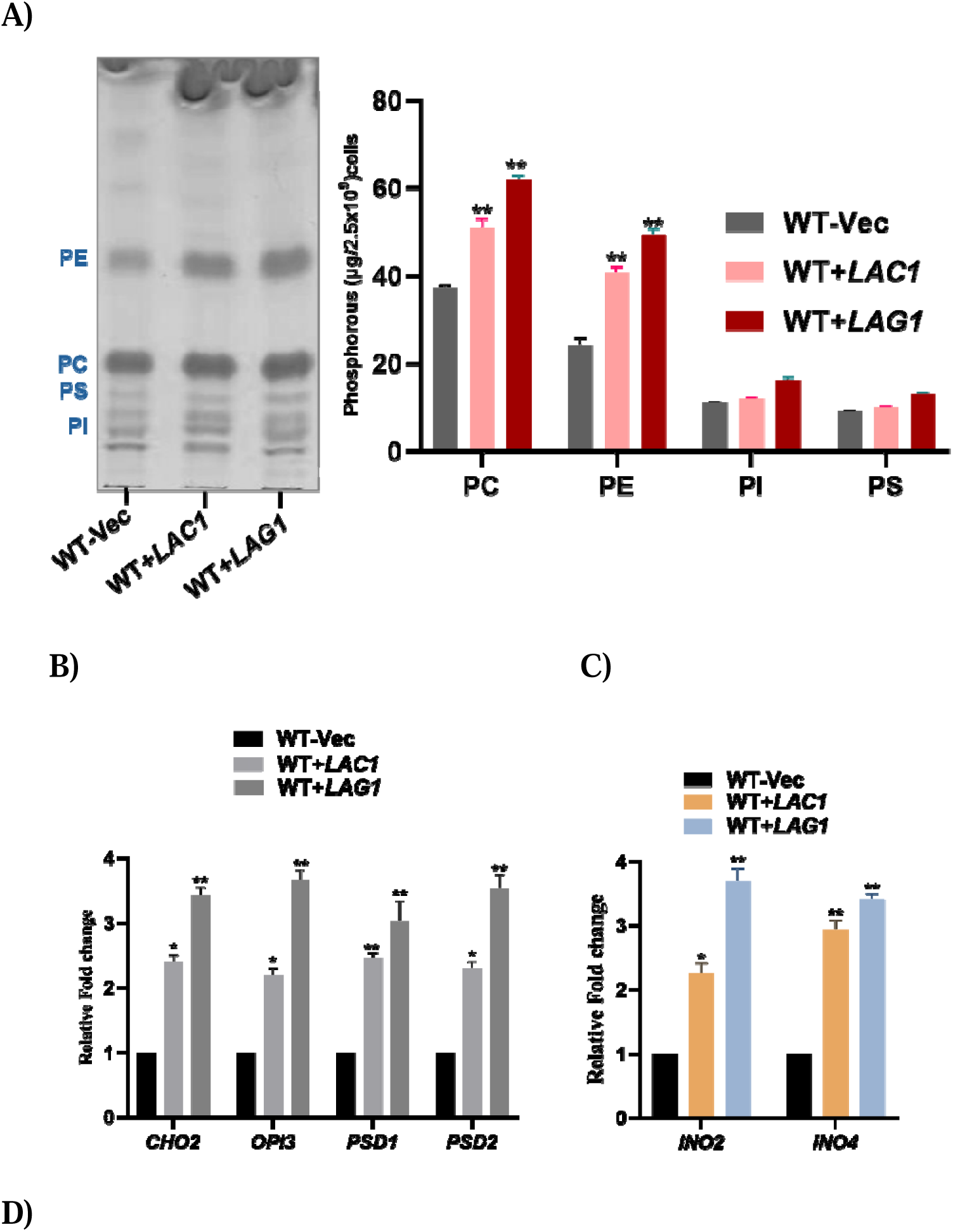
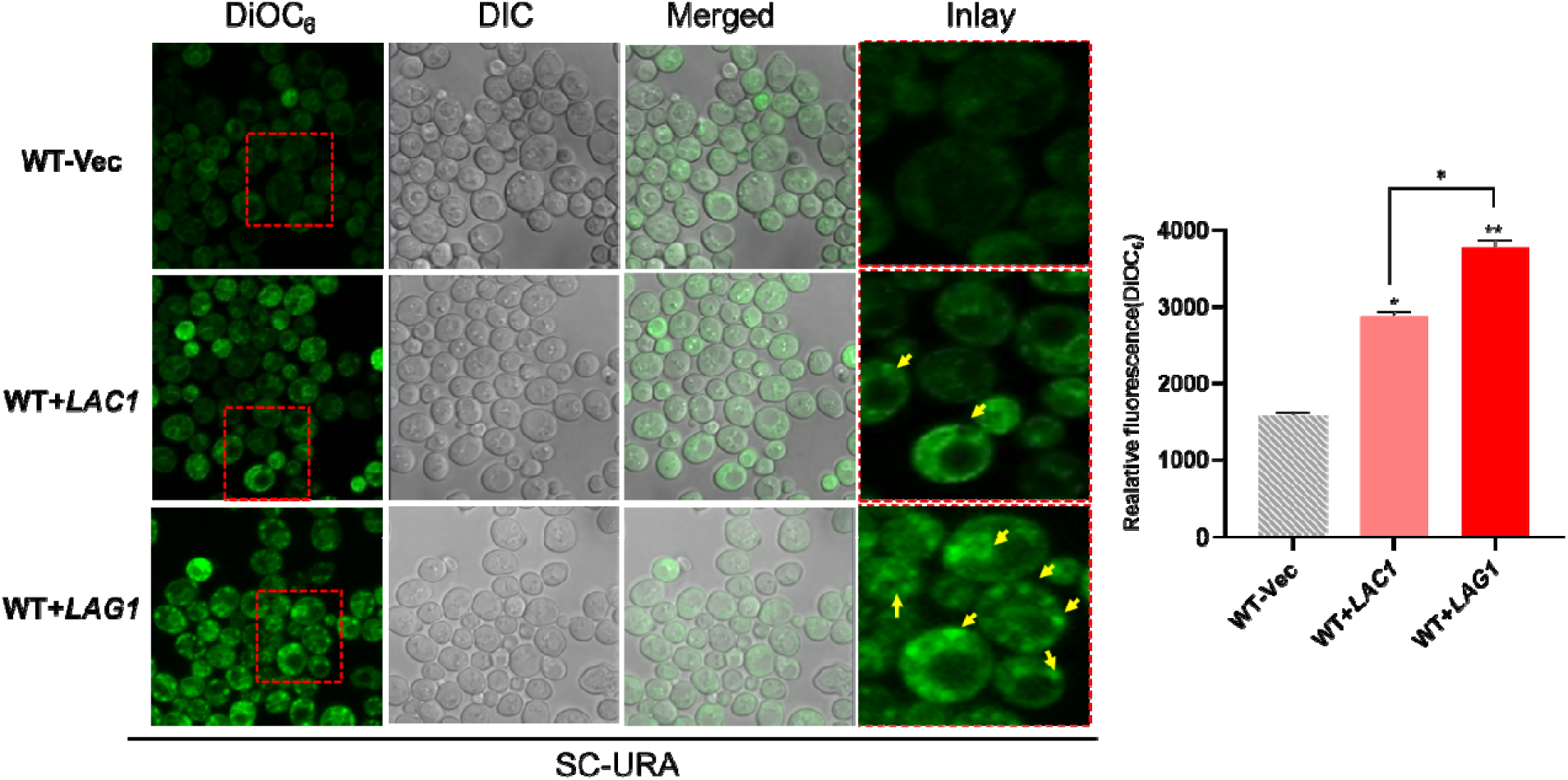
Wild-type cells overexpressed with *LAG1* exhibited elevated levels of phospholipids and showed an accumulation of membrane structures. (A) TLC was performed to analyze the composition of phospholipids, following the previously described protocol. (B and C) Q-PCR was employed to assess the expression of the genes of interest. (D) The staining of membrane structures was conducted using DiOC_6_, as explained previously. Data represents the mean ± SD (*P<0.05) in three separate experiments.

The regulation of PL biosynthesis is mediated by the *INO2-INO4* heterodimer, which binds to inositol/choline response elements (ICREs) to activate phospholipid synthesis via the CDP-diacylglycerol (CDP-DAG) pathway[41], [42]. Additionally, the unfolded protein response (UPR) pathway can stimulate the *INO2-INO4* complex, further enhancing the biosynthesis of phospholipids[36], [43]. Notably, *LAG1* overexpression induced a significant upregulation of *INO2* and *INO4* transcription compared to both *LAC1* overexpression and the vector control (Fig. 11C), pointing to a potential mechanism whereby *LAG1* activates the UPR pathway to support the increased demand for membrane lipid synthesis.

#### 3.12. *LAG1* overexpression reduced the Neutral lipid synthesis and lipid droplets

In WT+*LAG1* cells, levels of triacylglycerols (TAG) and free fatty acids (FFA) were significantly reduced compared to WT+LAC1 and vector control cells (Fig. 12A), while steryl esters (SE) showed a more moderate decrease in WT+*LAG1* cells compared to WT+*LAC1* and control cells. This decline in storage lipids was further corroborated by the marked downregulation of the genes *LRO1* and *DGA1*, which are essential for TAG biosynthesis. Similarly, the expression of *ARE1* and *ARE2*, involved in SE synthesis, was moderately reduced in WT+*LAG1* and WT+*LAC1* cells (Fig. 12B). Interestingly, sterol levels remained largely unchanged across all conditions (Fig. 12A).

**Figure 12.**
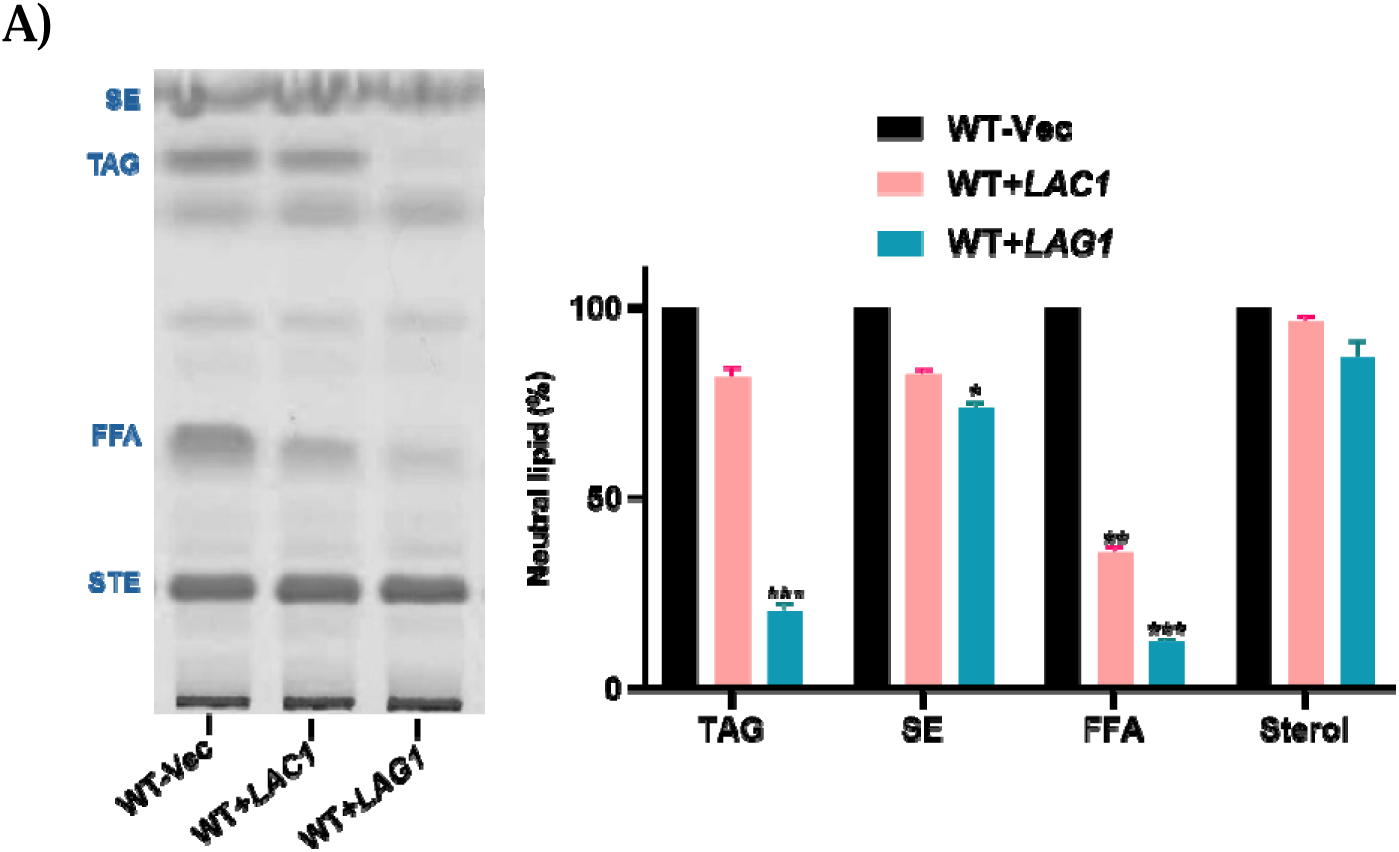
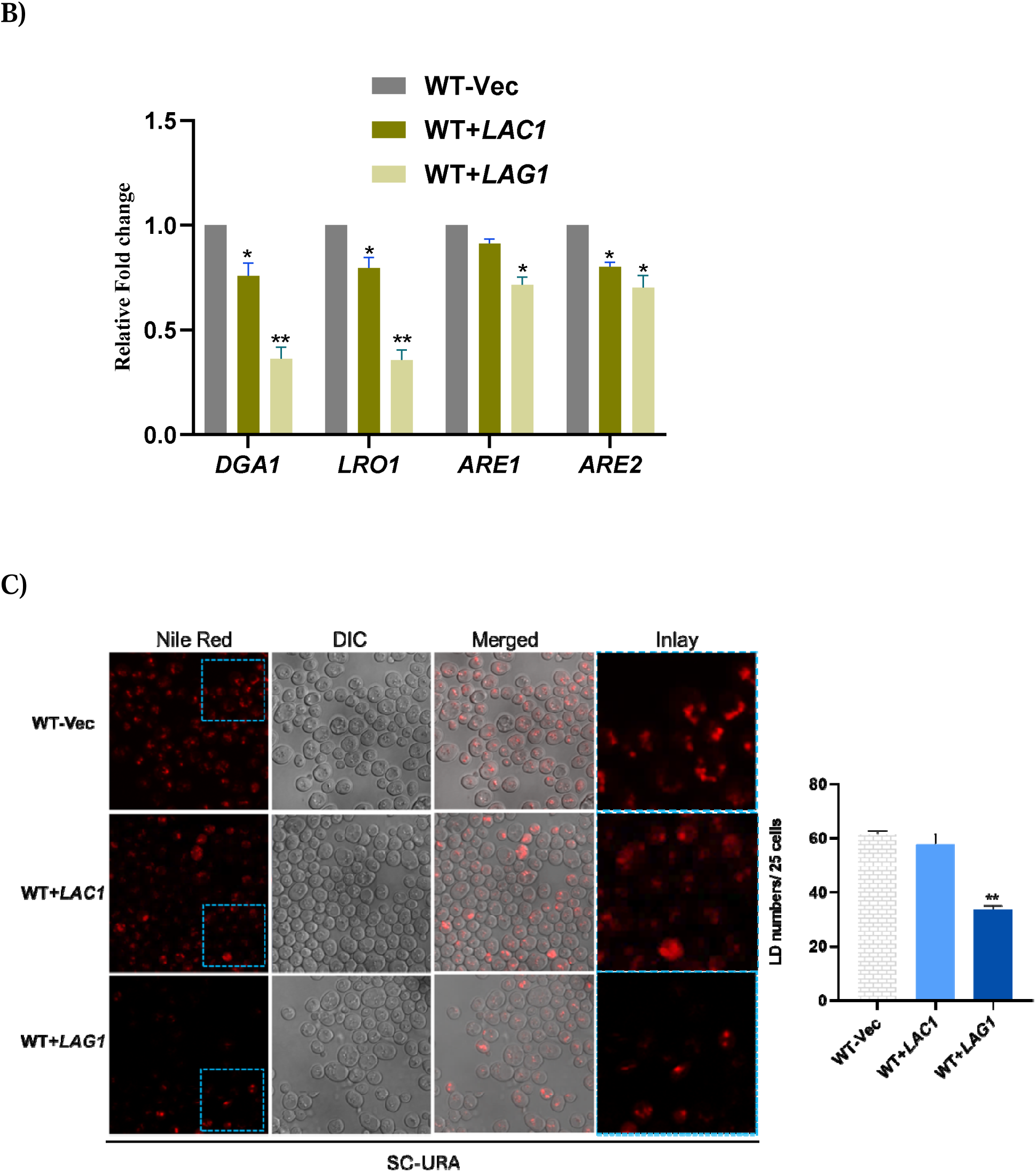
Neutral lipid levels dropped in cells that overexpressed *LAG1*. (A) TLC was used to separate the neutral lipids, as described in the methods chapter. Applying ImageJ software, the separated NL was evaluated, and the results were plotted. (B) Q-RT PCR was used to analyze the gene expression of neutral lipid-producing genes. (C) Nile Red staining was used to visualize LD, as explained before, using ImageJ software, lipid droplets were counted in 25 randomly chosen cells, and the findings were graphed. The data represents the mean ± SD (*P<0.05) from three separate experiments.

Furthermore, Nile Red staining highlighted a substantial reduction in lipid droplet (LD) numbers in cells overexpressing *LAG1* compared to those overexpressing *LAC1* and the control (Fig. 12C). These observations imply that *LAG1* overexpression may trigger a shift in lipid metabolism, leading to decreased storage lipids and a reduction in lipid droplet formation. The altered lipid balance may reflect a reallocation of resources from storage forms like TAG and SE toward membrane lipid synthesis and restructuring. Such a shift could support membrane expansion or specialized cellular functions induced by LAG1 overexpression.

### Regulation of *LAG1* overexpression mediated ER stress

#### 3.13. Overexpression of *LAG1* in *ORM1 ORM2* double deletion cells severely impaired cell growth

*ORM1* and *ORM2* are critical regulators of sphingolipid biosynthesis, primarily by inhibiting serine palmitoyl transferase (SPT), which catalyzes the rate-limiting step in ceramide production [44], [45](Fig.13A). This inhibition is essential for maintaining sphingolipid homeostasis and preventing excess sphingolipid accumulation, which could otherwise trigger ER stress and activate the UPR[45], [46]. The regulatory control exerted by ORM proteins involves phosphorylation events mediated by the TOR signaling pathway, which can modulate ORM activity in response to cellular conditions, finely tuning SPT activity and sphingolipid synthesis[47].

Upon *LAG1* overexpression, *ORM2* expression notably increases, likely as a compensatory response to heightened ceramide synthesis (Fig.13B). The loss of ORM proteins (e.g., in *orm1*Δ*orm2*Δ mutants) eliminates this regulatory control, leading to unrestrained SPT activity, sphingolipid buildup, and pronounced ER stress (Fig.13C). This suggests that *ORM1* and *ORM2* help prevent cellular stress by regulating sphingolipid levels under normal conditions and that their deletion or dysregulation alongside *LAG1* overexpression disrupts this balance, significantly impairing cell growth compared to control vector cells with the same deletions (Fig. 13C). Under ER stress conditions, *orm1*Δ*orm2*Δ cells overexpressing *LAG1* showed no growth.

**Figure 13.**
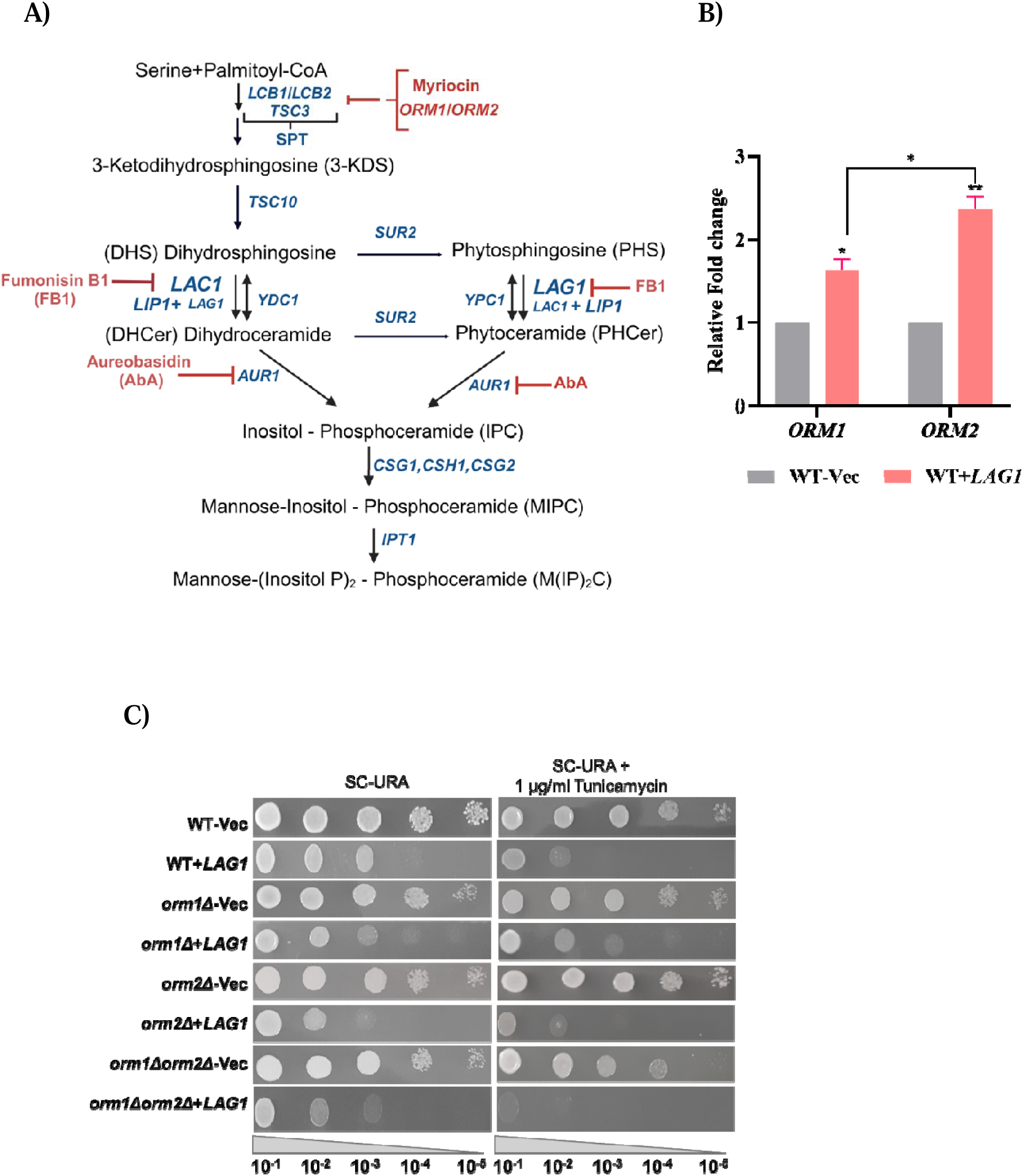
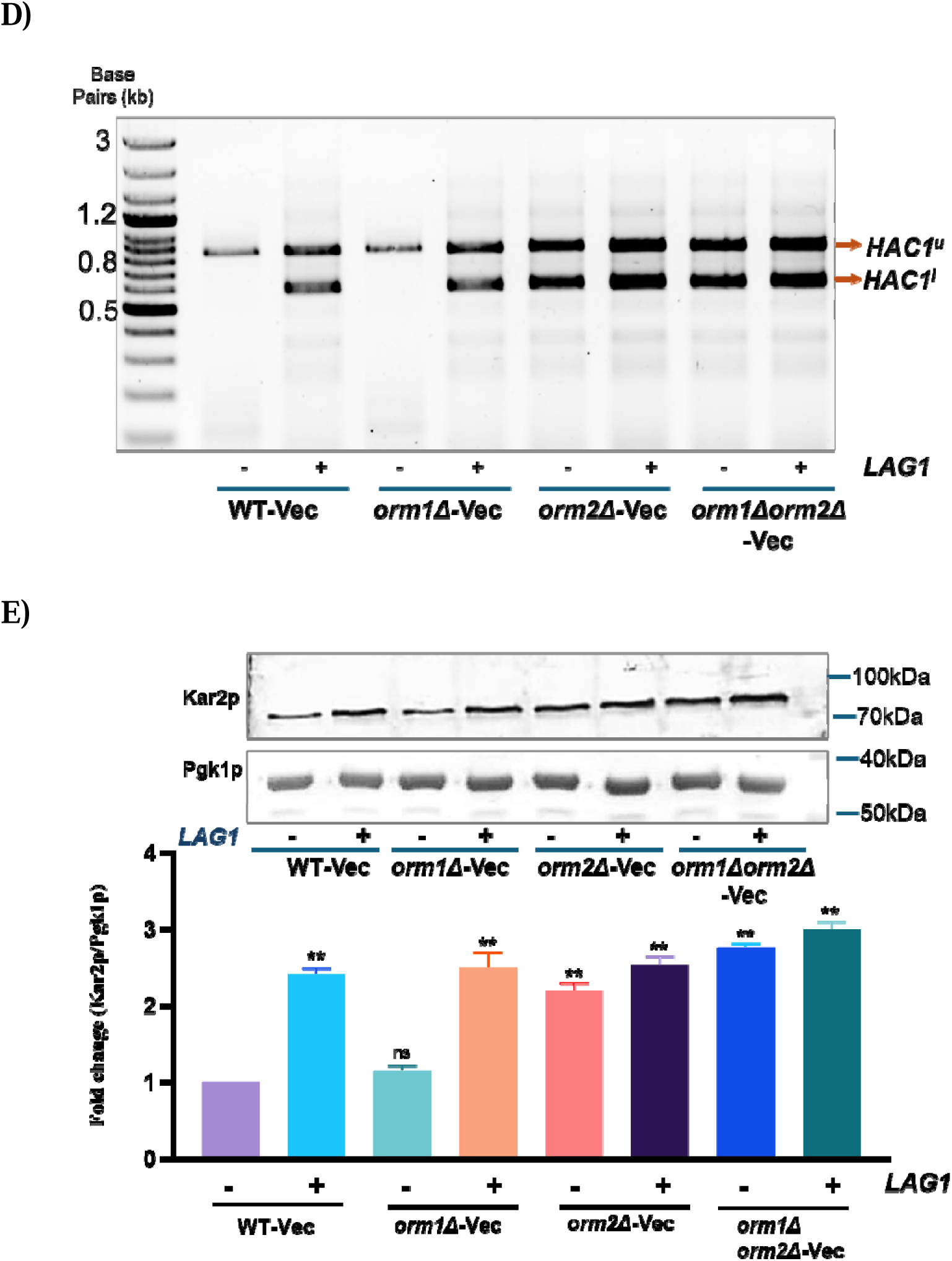
ORM genes tightly regulate the expression of ceramide synthesizing genes, including *LAG1*. (A) overview of yeast sphingolipid metabolism and its regulators. (B) Q-RT PCR was used to analyze the gene expression of *ORM* genes. (C)The cells were cultured in SC media at 30°C till the mid-log phase. Following cell harvesting, an equal optical density (OD of 3.0 at A600 nm) was used to create a 10-fold serial dilution. These diluted cells were subsequently spotted onto SC-Ura media supplemented with 2% galactose. (D) 1% agarose gel was used to separate the PCR products produced by the HAC1 mRNA splicing experiment (E)A 12% SDS-PAGE gel was loaded with identical protein concentrations. The primary antibody used was anti-Kar2p, while Pgk1p was the loading control. The data represents the mean ± SD (*P<0.05) of three replicates from three separate experiments.

*HAC1* splicing was observed in *orm2*Δ and *orm1*Δ*orm2*Δ deletion strains. Still, not in *orm1*Δ cells unless co-expressed with *LAG1* (Fig. 13D). Additionally, analysis of Kar2p levels revealed substantial increases in *orm2*Δ and *orm1*Δ*orm2*Δ cells, with an even higher rise in cells co-expressing LAG1, particularly in *orm1*Δ*orm2*Δ strains (Fig. 13E). These results suggest that the dual impact of *orm1*Δ*orm2*Δ deletions and LAG1 overexpression disrupts sphingolipid homeostasis, exacerbating ER stress, impairing growth, and potentially jeopardizing cell survival.

#### 3.14. Myriocin was ineffective in alleviating the consequences of *LAG1* overexpression

Myriocin, a potent SPT inhibitor, significantly impacts sphingolipid metabolism and its regulatory feedback mechanisms[48]. By inhibiting SPT activity, myriocin reduces sphingolipid synthesis, which often leads to adaptive changes, such as increased phosphorylation of *ORM1* and modifications in *ORM2* expression [45], [49](Fig.13A). Interestingly, while *ORM1* is responsive to phosphorylation alterations triggered by myriocin, *ORM2* remains unaffected, suggesting that these proteins play distinct roles in maintaining sphingolipid balance during stress conditions[10].

To evaluate the potential protective effects of myriocin, a sub-lethal dose was applied to cells overexpressing *LAG1*; it was found that it did not yield a significant enhancement in cell growth (Fig. 14A and B). Additionally, the splicing status of *HAC1* and the expression levels of Kar2p remained unchanged compared to those in cells solely overexpressing *LAG1* (Fig.14C and D). This indicates that a sub-lethal dose of myriocin was ineffective in alleviating the ER stress associated with *LAG1* overexpression. These findings imply that inhibiting SPT activity with myriocin is inadequate to counteract the stress induced by elevated ceramide levels resulting from *LAG1* overexpression.

**Figure 14.**
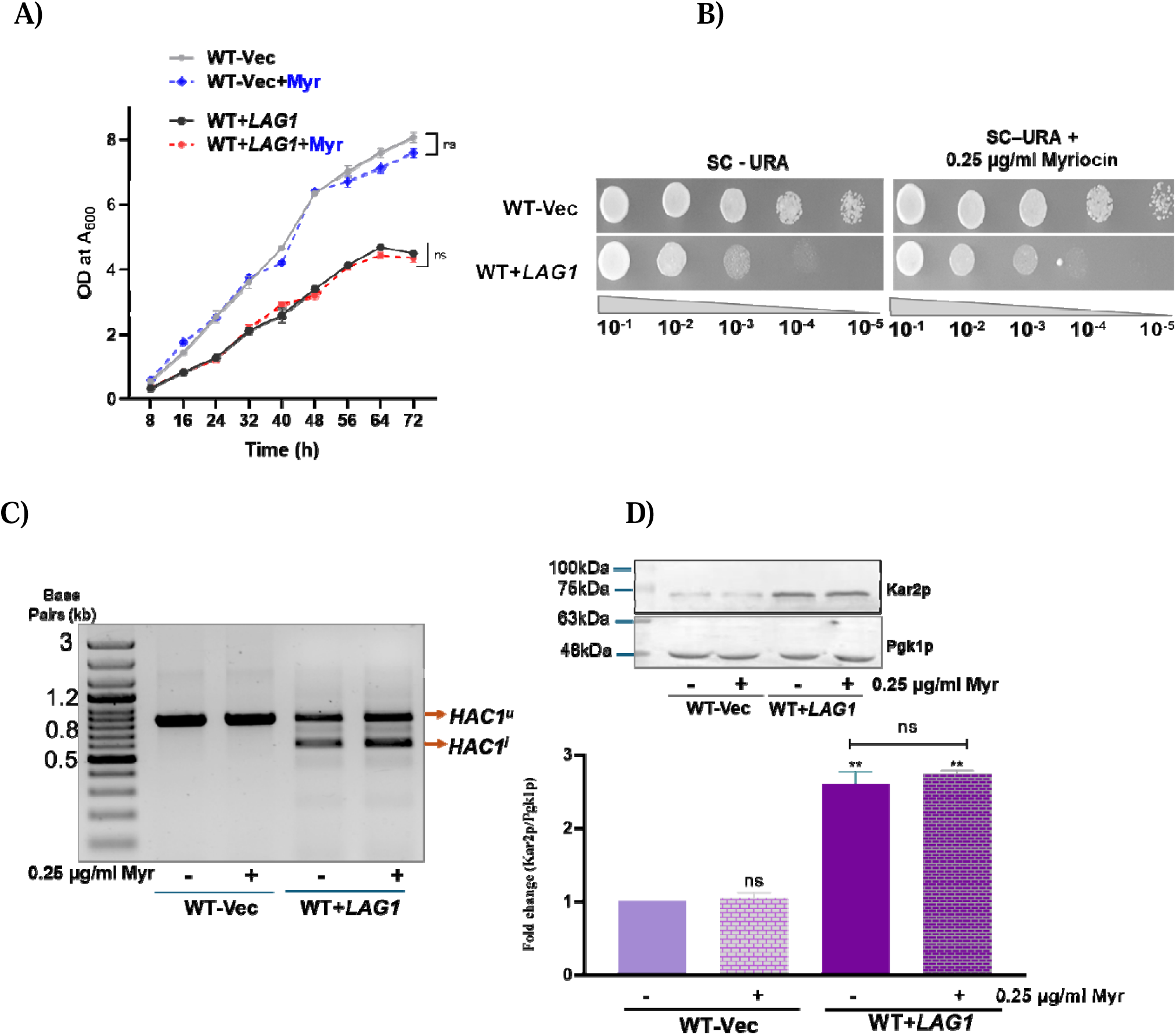
Myriocin did not alleviate ER stress caused by LAG1 overexpression. (A) Growth of WT-Vec, WT+*LAG1* strains in SC-URA+ Galactose medium without and with 0.25 µg/ml Myriocin. The graphs show absorbance at 600 nm measured every 8 hours, with time points displayed on the x-axis using a plate reader. (B) Cells with the same OD (3.0 at A_600 nm_) were serially diluted (10-fold) and grown as described in the methods section. These cells were then spotted on SC agar plates, with or without myriocin, and incubated for 48 hours at 30°C. (C) 1% agarose gel was used to separate the PCR products produced by the HAC1 mRNA splicing experiment). (D) A 12% SDS-PAGE gel was loaded with identical protein concentrations. The primary antibody used was anti-Kar2p, while Pgk1p was the loading control. The data represent the mean ± SD (*P<0.05) of three replicates from three separate experiments.

#### 3.15. Ceramide synthase inhibitor Fumonisin B1 rescued growth in cells overexpressed with LAG1 to a certain extent

Fumonisin B1 (FB1), a mycotoxin that structurally resembles sphingoid bases, allows it to interfere with ceramide synthases, specifically *LAC1* and *LAG1*, which facilitate the transfer of fatty acids from fatty acyl-CoAs to sphinganine and sphingosine[50] (Fig. 15A). The expression of *LAG1* in *orm1*Δ*orm2*Δ cells results in marked growth impairment, highlighting the lethal effects associated with elevated sphingolipid levels in conjunction with *LAG1* overexpression (Fig. 13C).

**Figure 15.**
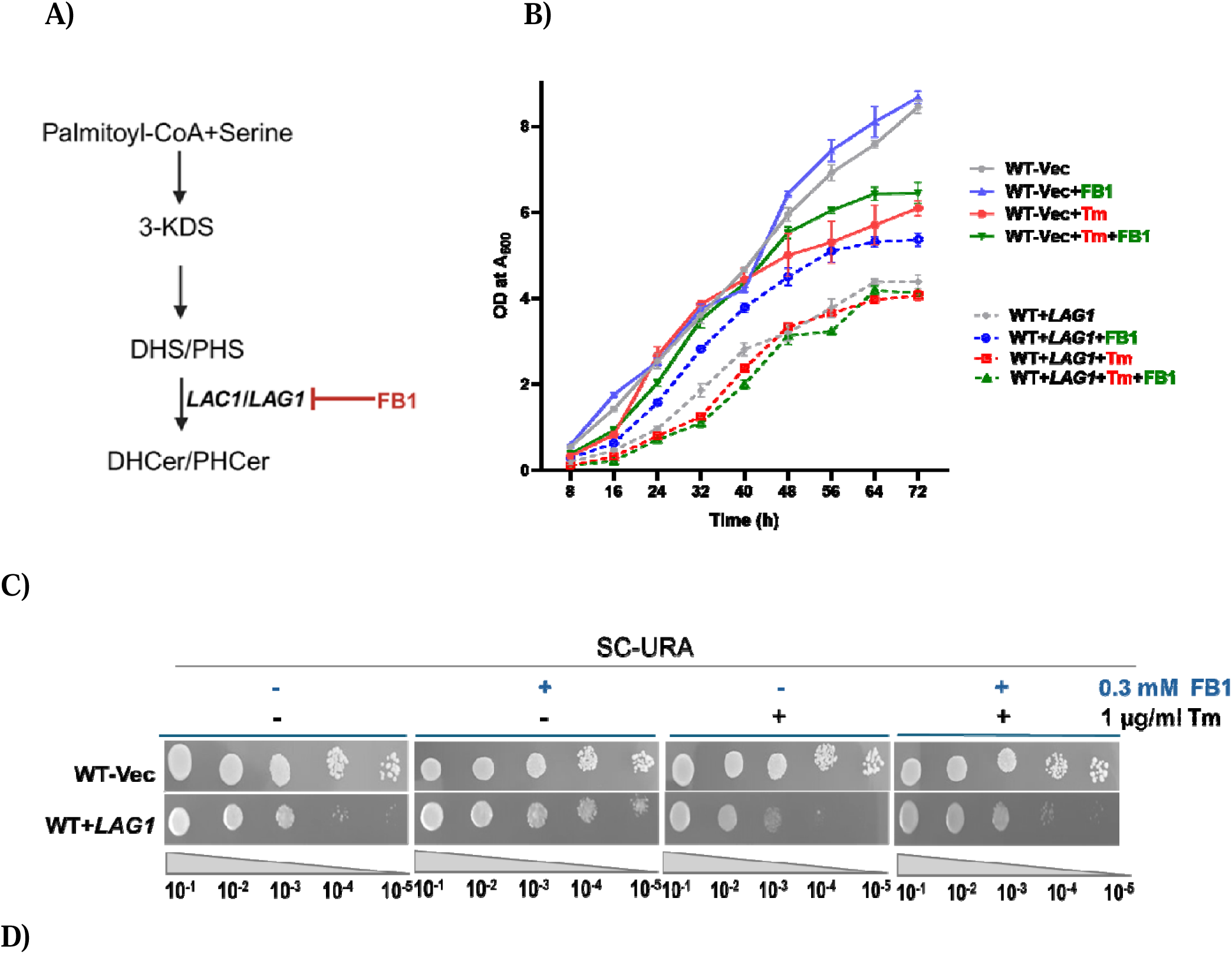
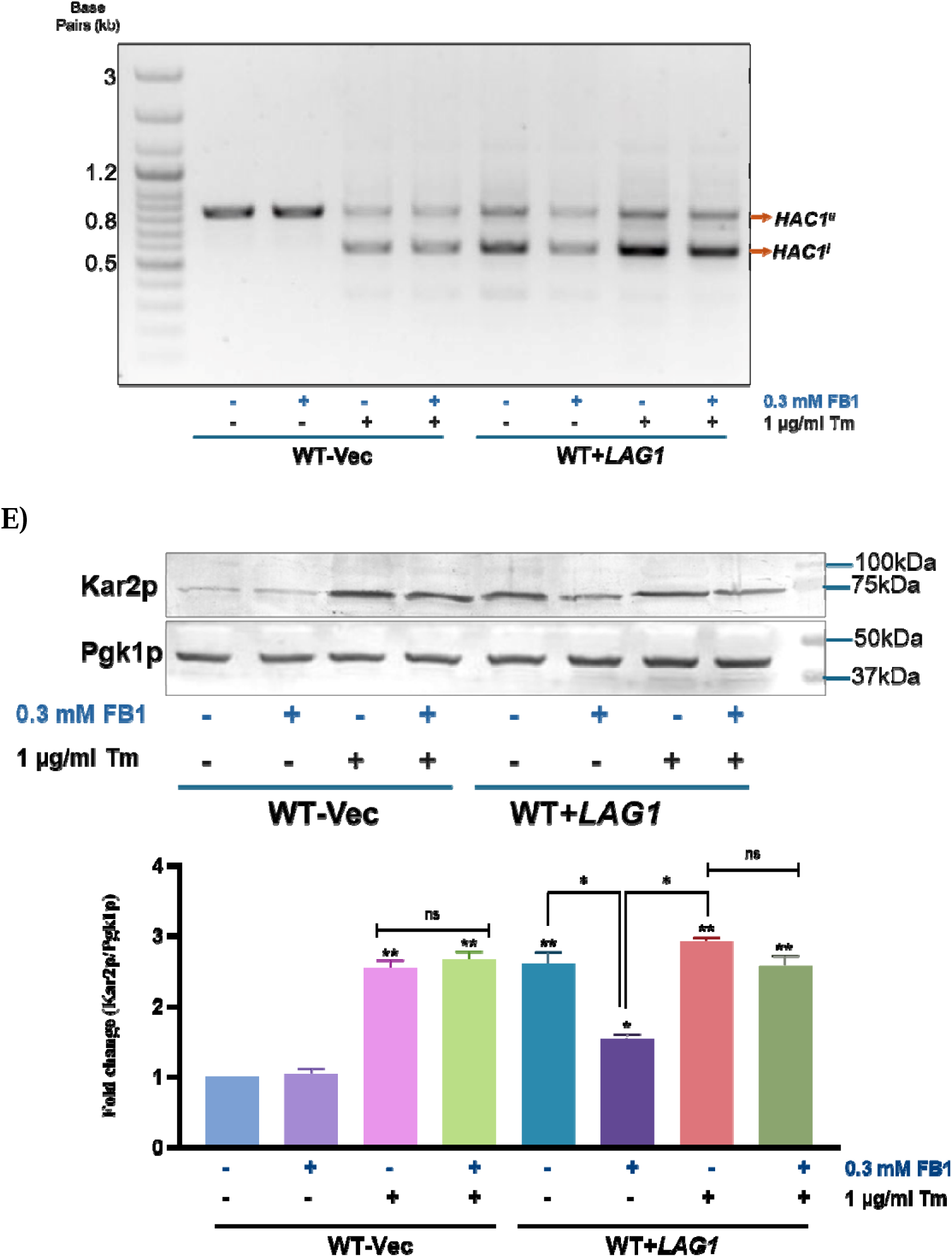
Fumonisin B1 ameliorated the ER stress observed in the *LAG1* overexpression cells. (A) overview of the role of FB1 in regulating ceramide biosynthesis. (B)Spot test was performed as described before. (C) The growth of WT-Vec and WT+*LAG1* strains was evaluated in SC-URA + galactose medium, with and without 1 µg/ml tunicamycin and/or 0.3 mM Fumonisin B1 (FB1). Absorbance at 600 nm was measured every 8 hours, with time points displayed on the x-axis using a plate reader (D) 1% agarose gel was used to separate the PCR products produced by the HAC1 mRNA splicing experiment and records of the gel electrophoresis data were made. (E) Equal concentrations of respective proteins were loaded in 12% SDS-PAGE gel. The primary antibody was anti-Kar2p, while the Pgk1p antibody was the loading control. The data represents the mean ± SD (*P<0.05) of three replicates from three separate experiments.

A series of tests involving various concentrations of FB1 (data not shown) showed that a sublethal concentration of 0.3 mM was optimal for promoting cell growth in WT+*LAG1* cells. While treatment with this concentration improved growth (Fig. 15B and C), it failed to reverse *HAC1* splicing (Fig.15D) and was linked to further reductions in Kar2 protein levels in WT+*LAG1* cells compared to untreated controls (Fig. 15E). Additionally, the introduction of tunicamycin alongside FB1 negated the growth-promoting effects of FB1, resulting in elevated Kar2 protein expression in WT+*LAG1* cells (Fig. 15E). This dynamic illustrates the complex interactions between sphingolipid metabolism and cellular stress responses in the context of *LAG1* overexpression, indicating the necessity for further research into the underlying mechanisms involved.

## 4.) DISCUSSION AND CONCLUSION

The interplay between ceramides and cellular quality control networks and the interconnectedness of their metabolic pathways warrants further exploration. Defective proteostasis significantly contributes to various human diseases, including cancer, highlighting the need to investigate strategies for alleviating endoplasmic reticulum (ER) stress with potential therapeutic implications[51]. The human tumor metastasis suppressor gene TMSG1, homologous to the yeast gene *LAG1*, has been linked to cancer progression and metastasis when silenced[52], [53]. Conversely, knockout studies of the mouse homolog LASS2 have shown protective effects against hepatic steatosis, while its overexpression is associated with cell cycle arrest, increased oxidative stress, and enhanced apoptosis, ultimately inhibiting cancer progression[52]. Emerging evidence indicates that ceramides play vital roles in various cellular processes, including actin cytoskeleton polarization, endosomal transport, autophagy, and apoptosis[51]. Notably, the ER stress response observed in yeast mirrors inflammatory responses in higher organisms, making the investigation of yeast ceramide synthases *LAG1* and *LAC1* in the context of ER stress crucial for understanding pathophysiological processes in more complex systems.

Traditionally, the LAG genes *LAG1* and *LAC1* have been recognized primarily for their roles in synthesizing dihydroceramide and phytoceramide[19] (Fig.13A). However, recent research has uncovered their additional contributions to maintaining the ER stress response and lipid homeostasis. Sphingolipids, which comprise long-chain bases (LCBs) and fatty acids, utilize serine as a backbone instead of glycerol[54]. These molecules are vital for structural integrity and signaling in various biological processes, with functional diversity arising from modifications such as phosphorylation, hydroxylation, and desaturation. The enzyme *SUR2* hydroxylates LCBs in ceramide, influencing their physical properties and interactions[55], [56].

*SUR2* catalyzes the conversion of dihydrosphingosine (DHS) and dihydroceramide (DHCer) into phytosphingosine (PHS) and phytoceramide (PHCer), respectively, affecting substrate availability for ceramide synthases *LAG1* and *LAC1* [19](Fig.S4A). Recent findings show that *LAG1* preferentially utilizes PHS to produce phytoceramide PHCer, while *LAC1* favors DHS for DHCer synthesis, denoted in bold font [19](Fig.S4A). The *LAG1* and *SUR2* double deletion exhibited increased resistance to ER stress, similar to the resistance observed with *LAG1* deletion alone, while the *LAC1* and *SUR2* double deletion demonstrated growth patterns comparable to WT cells (Fig.S4B). Furthermore, *HAC1* splicing and elevated levels of Kar2p were absent in *sur2*Δ*, sur2*Δ*lac1*Δ, and *sur2*Δ*lag1*Δ cells, yielding results akin to WT cells (Fig.S4C and D). Recent studies indicate that PHCer is critical for maintaining diffusion barriers in the ER; LAG1, primarily localized in the nuclear ER, prefers PHS as its substrate and converts it into PHCer. Sphingolipids are key components of the ER stress surveillance pathway (ERSU), ensuring that only healthy ER is transmitted to daughter cells[57], [58]. Thus, the role of PHCer in relation to ER stress warrants further examination.

In the context of ER stress, the non-redundant roles of *LAC1* and *LAG1* become apparent. The deletion of *LAG1* did not compromise the ER stress response; *lag1*Δ cells exhibited expression levels of ER stress response genes comparable to those of WT and *lac1*Δ cells, underscoring the necessity for a functional ER stress response. Further analysis reveals that the unfolded protein response (UPR) is essential for *lag1*Δ cells to buffer against ER stress. Notably, phosphatidylcholine (PC) and phosphatidylethanolamine (PE) levels are crucial for the role of *LAG1* in alleviating ER stress. The disruption in lipid homeostasis and the protective role against ER stress were restored to normal when *LAG1* deletion cells were complemented with pRS416-*LAG1* (data not shown). In cells overexpressing *LAG1*, an intrinsic ER stress phenotype emerged, characterized by reduced growth and survival. Under external ER stress conditions, growth declined further, with no growth observed at higher doses of tunicamycin. *LAG1* overexpression triggered the UPR, increased the expression of ER stress response genes, and led to the accumulation of immature carboxypeptidase Y (CPY). Remarkably, *LAG1* overexpression resulted in elevated levels of *INO2* and *INO4*, enhancing phospholipid (PL) synthesis while concurrently decreasing neutral lipid synthesis and lipid droplet numbers, diverging from typical ER stress responses (Fig.16). In *orm1*Δ*orm2*Δ cells, *LAG1* expression severely impaired growth; however, treatment with Myriocin, an SPT inhibitor, did not ameliorate the growth defects associated with *LAG1* overexpression. In contrast, Fumonisin B1, a ceramide/sphingolipid synthesis inhibitor, provided slight growth rescue in *LAG1*-overexpressing cells, suggesting that *LAG1* plays a unique and critical role in maintaining ER quality control beyond ceramid synthesis.

**Figure 16.**
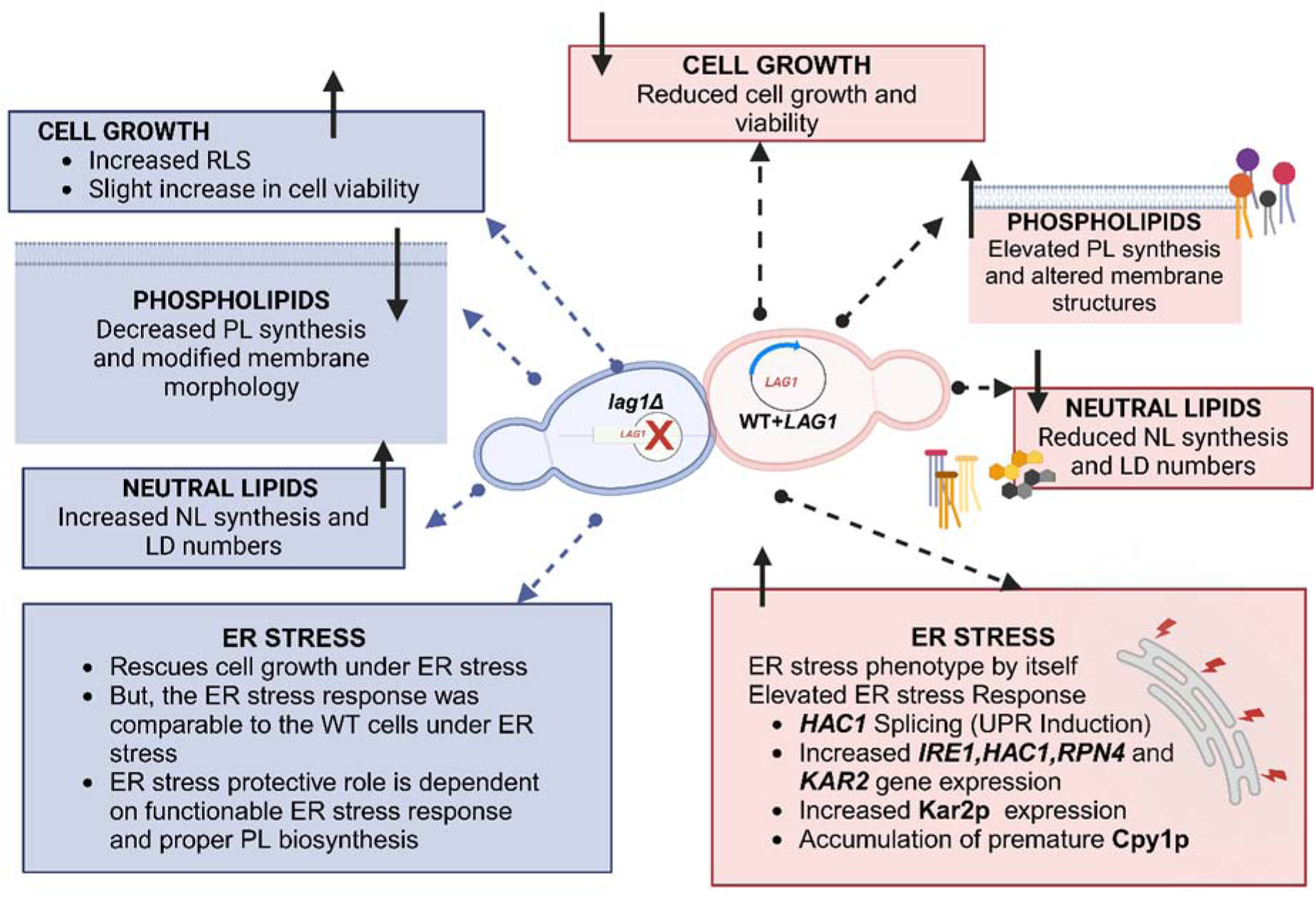
An illustration showcasing the effects of *LAG1* loss and overexpression on various cellular functions. *LIP1*, known as the *LAC1/LAG1* Interacting Protein, is a crucial regulatory subunit that forms a complex with *LAG1* and *LAC1*[59] (Fig.S5A). The loss of *LIP1* has been shown to reduce ceramide synthesis and may affect the localization and phosphorylation status of *LAG1*[59]. While *LAC1* has a distinct role in responding to ER stress compared to *LAG1*, we overexpressed *LAG1* in cells lacking *LIP1* and *LAC1* to investigate their functions further. As reported earlier, deletion of *LIP1* conferred resistance to cells treated with AbA by decreasing ceramide synthesis[38]. In *LAG1*-overexpressing cells, the deletion of *LIP1* slightly improved cell growth, suggesting a partial compensatory effect (Fig.S5B). In contrast, no changes in cell growth were observed when LAG1 was overexpressed in cells lacking LAC1(Fig.S5B), underscoring the unique contributions of these proteins to cell growth and stress response. Nevertheless, despite the partial rescue of cell growth in *LAG1*-overexpressing cells with *LIP1* deletion, no alterations in *HAC1* splicing or elevated Kar2p expression were noted (Fig.S5C and D).

Despite reduced growth in cells lacking both *LAG1* and *LAC1*, ceramide synthesis was sustained by alkaline ceramidases *YPC1* and *YDC1*, which condense serine with long-chain bases through a reverse reaction[60] (Fig.13A) (Fig.S6A). Reports indicate that even with the deletion of *LAG1, LAC1, YPC1,* and *YDC1*, cells continue to produce low levels of sphingolipids. In this study, *LAG1* was overexpressed in *ypc1*Δ*, ydc1*Δ, and *ypc1*Δ *ydc1*Δ double deletion cells, revealing no significant change in cell growth [5](Fig.S6B); growth levels were comparable to those seen with *LAG1* overexpression in WT cells. Recent findings suggest that *LAG1* primarily acts on PHS to convert it into PHCer. Intriguingly, the deletion of *YPC1*, which functions as the yeast phytoceramidase, did not impact cell growth when *LAG1* was overexpressed. *HAC1* splicing and elevated Kar2p expression were observed only in *YDC1* and/or *YPC1* deletion cells co-expressing *LAG1* (Fig.S6C and D).

In conclusion, this research confirms the essential roles of LAG genes in modulating ER stress and lipid metabolism. The distinct functions of *LAG1* and *LAC1*, along with the regulatory influences of various factors on *LAG1* gene expression, have been elucidated, enhancing our understanding of the additional roles of *LAG1* in cellular processes. These findings contribute to our fundamental knowledge of ceramide biology and open avenues for potential therapeutic strategies targeting ER stress-related diseases.

