## Supplementary information for "Differential Roles of Longevity Assurance Genes *LAG1* and *LAC1* in Regulating Endoplasmic Reticulum stress and Lipid Homeostasis in *Saccharomyces cerevisiae*"

**Supplementary Data**

**Table S1. The strains of yeast and E. coli and plasmids utilized in this research**

| **Name of the strains** | **Descriptive genotype** | **Origin** |
| --- | --- | --- |
| Wild-Type (WT) | BY4741; Mat a; his3Δ 1; leu2Δ 0; met15Δ 0; ura3Δ 0 | Euroscarf |
| *lac1∆* | BY4741; Mat a; his3Δ 1; leu2Δ 0; met15Δ 0; ura3Δ 0; YKL008c:: kanMX4 | Prof. Maya Schuldiner |
| *lag1∆* | BY4741; Mat a; his3Δ 1; leu2Δ 0; met15Δ 0; ura3Δ 0; YHL003c:: kanMX4 | Prof. Maya Schuldiner |
| *rpn4∆* | BY4741; Mat a; his3Δ 1; leu2Δ 0; met15Δ 0; ura3Δ 0; YDL020c:: kanMX4 | Prof. Shantanu Sengupta |
| *ire1∆* | BY4741; Mat a; his3Δ 1; leu2Δ 0; met15Δ 0; ura3Δ 0; YHR079c:: kanMX4 |  |
| *hac1∆* | BY4741; Mat a; his3Δ 1; leu2Δ 0; met15Δ 0; ura3Δ 0; YFL031w:: kanMX4 | ” |
| *cho2∆* | BY4741; Mat a; his3Δ 1; leu2Δ 0; met15Δ 0; ura3Δ 0; YGR157w:: kanMX4 | Euroscarf |
| *opi3∆* | BY4741; Mat a; his3Δ 1; leu2Δ 0; met15Δ 0; ura3Δ 0; YJR073c:: kanMX4 | Euroscarf |
| *psd1∆* | BY4741; Mat a; his3Δ 1; leu2Δ 0; met15Δ 0; ura3Δ 0; YNL169c:: kanMX4 | Euroscarf |
| *psd2Δ* | BY4741; Mat a; his3Δ 1; leu2Δ 0; met15Δ 0; ura3Δ 0; YGR170w:: kanMX4 | Euroscarf |
| *ino2∆* | BY4741; Mat a; his3Δ 1; leu2Δ 0; met15Δ 0; ura3Δ 0; YDR123c:: kanMX4 | Euroscarf |
| *ino4∆* | BY4741; Mat a; his3Δ 1; leu2Δ 0; met15Δ 0; ura3Δ 0; YOL108c:: kanMX4 | Euroscarf |
| *tor1∆* | BY4741; Mat a; his3Δ 1; leu2Δ 0; met15Δ 0; ura3Δ 0; YJR066w:: kanMX4 | Euroscarf |
| *lro1Δ* | BY4741; Mat a; his3Δ 1; leu2Δ 0; met15Δ 0; ura3Δ 0; YNR008w:: kanMX4 | ” |
| *dga1Δ* | BY4741; Mat a; his3Δ 1; leu2Δ 0; met15Δ 0; ura3Δ 0; YOR245c:: kanMX4 |  |
| *are1Δ* | BY4741; Mat a; his3Δ 1; leu2Δ 0; met15Δ 0; ura3Δ 0; YCR048w:: kanMX4 | ” |
| *are2Δ* | BY4741; Mat a; his3Δ 1; leu2Δ 0; met15Δ 0; ura3Δ 0; YNR019w:: kanMX4 |  |
| *orm1Δ* | BY4741; Mat a; his3Δ 1; leu2Δ 0; met15Δ 0; ura3Δ 0; YGR038w:: kanMX4 | Prof. Maya Schuldiner |
| *orm2Δ* | BY4741; Mat a; his3Δ 1; leu2Δ 0; met15Δ 0; ura3Δ 0; YLR350w:: kanMX4 | Prof. Maya Schuldiner |
| *orm1∆orm2∆* | BY4741; Mat a; his3Δ 1; leu2Δ 0; met15Δ 0; ura3Δ 0; YGR038W::kanMX4;orm2Δ::LEU2 | This study |
| *lac1∆lag1∆* | BY4741; Mat a; his3Δ 1; leu2Δ 0; met15Δ 0; ura3Δ 0; YKL008c:: kanMX4;lag1Δ::LEU2 | This study |
| *cho2∆lac1∆* | BY4741; Mat a; his3Δ 1; leu2Δ 0; met15Δ 0; ura3Δ 0; YGR157w:: kanMX4; lac1Δ::LEU2 | This study |
| *cho2∆lag1∆* | BY4741; Mat a; his3Δ 1; leu2Δ 0; met15Δ 0; ura3Δ 0; YGR157w:: kanMX4; lag1Δ::LEU2 | This study |
| *opi3Δ lac1Δ* | BY4741; Mat a; his3Δ 1; leu2Δ 0; met15Δ 0; ura3Δ 0; YJR073c:: kanMX4;lac1Δ::LEU2 | This study |
| *opi3Δ lag1Δ* | BY4741; Mat a; his3Δ 1; leu2Δ 0; met15Δ 0; ura3Δ 0; YJR073c:: kanMX4; lag1Δ::LEU2 | This study |
| *psd1Δlac1Δ* | BY4741; Mat a; his3Δ 1; leu2Δ 0; met15Δ 0; ura3Δ 0; YNL169c:: kanMX4;lac1Δ::LEU2 | This study |
| *psd1Δlag1Δ* | BY4741; Mat a; his3Δ 1; leu2Δ 0; met15Δ 0; ura3Δ 0; YNL169c:: kanMX4;lag1Δ::LEU2 | This study |
| *psd2Δlac1Δ* | BY4741; Mat a; his3Δ 1; leu2Δ 0; met15Δ 0; ura3Δ 0; YGR170w:: kanMX4; lac1Δ::LEU2 | This study |
| *psd2Δlag1Δ* | BY4741; Mat a; his3Δ 1; leu2Δ 0; met15Δ 0; ura3Δ 0; YGR170w:: kanMX4; lag1Δ::LEU2 | This study |
| *lro1Δlac1Δ* | BY4741; Mat a; his3Δ 1; leu2Δ 0; met15Δ 0; ura3Δ 0; YNR008w:: kanMX4; lac1Δ::LEU2 | This study |
| *lro1Δlag1Δ* | BY4741; Mat a; his3Δ 1; leu2Δ 0; met15Δ 0; ura3Δ 0; YNR008w:: kanMX4; lag1Δ::LEU2 | This study |
| *dga1Δlac1Δ* | BY4741; Mat a; his3Δ 1; leu2Δ 0; met15Δ 0; ura3Δ 0; YOR245c:: kanMX4; lac1Δ::LEU2 | This study |
| *dga1Δlag1Δ* | BY4741; Mat a; his3Δ 1; leu2Δ 0; met15Δ 0; ura3Δ 0; YOR245c:: kanMX4; lag1Δ::LEU2 | This study |
| *are1Δlac1Δ* | BY4741; Mat a; his3Δ 1; leu2Δ 0; met15Δ 0; ura3Δ 0; YCR048w:: kanMX4; lac1Δ::LEU2 | This study |
| *are1Δlag1Δ* | BY4741; Mat a; his3Δ 1; leu2Δ 0; met15Δ 0; ura3Δ 0; YCR048w:: kanMX4; lag1Δ::LEU2 | This study |
| *are2Δlac1Δ* | BY4741; Mat a; his3Δ 1; leu2Δ 0; met15Δ 0; ura3Δ 0; YNR019w:: kanMX4; lac1Δ::LEU2 | This study |
| *are2Δlag1Δ* | BY4741; Mat a; his3Δ 1; leu2Δ 0; met15Δ 0; ura3Δ 0; YNR019w:: kanMX4; lag1Δ::LEU2 | This study |
| *ire1Δlac1Δ* | BY4741; Mat a; his3Δ 1; leu2Δ 0; met15Δ 0; ura3Δ 0; YHR079c:: kanMX4; lac1Δ::LEU2 | This study |
| *ire1Δlag1Δ* | BY4741; Mat a; his3Δ 1; leu2Δ 0; met15Δ 0; ura3Δ 0; YHR079c:: kanMX4; lag1Δ::LEU2 | This study |
| *hac1Δlac1Δ* | BY4741; Mat a; his3Δ 1; leu2Δ 0; met15Δ 0; ura3Δ 0; YFL031w:: kanMX4; lac1Δ::LEU2 | This study |
| *hac1Δlag1Δ* | BY4741; Mat a; his3Δ 1; leu2Δ 0; met15Δ 0; ura3Δ 0; YFL031w:: kanMX4; lag1Δ::LEU2 | This study |
| *rpn4Δlac1Δ* | BY4741; Mat a; his3Δ 1; leu2Δ 0; met15Δ 0; ura3Δ 0; YDL020c:: kanMX4; lac1Δ::LEU2 | This study |
| *rpn4Δlag1Δ* | BY4741; Mat a; his3Δ 1; leu2Δ 0; met15Δ 0; ura3Δ 0; YDL020c:: kanMX4; lag1Δ::LEU2 | This study |
| *sur2Δ* | BY4741; Mat a; his3Δ 1; leu2Δ 0; met15Δ 0; ura3Δ 0; YDR297w:: kanMX4 | Prof. Maya Schuldiner |
| *sur2Δlac1Δ* | BY4741; Mat a; his3Δ 1; leu2Δ 0; met15Δ 0; ura3Δ 0; YDR297w::kanMX4; lac1Δ::LEU2 | Prof. Maya Schuldiner |
| *sur2Δlag1Δ* | BY4741; Mat a; his3Δ 1; leu2Δ 0; met15Δ 0; ura3Δ 0; YDR297w:: kanMX4; lag1Δ::LEU2 | Prof. Maya Schuldiner |
| *ydc1Δ* | BY4741; Mat a; his3Δ 1; leu2Δ 0; met15Δ 0; ura3Δ 0; YDR297w:: kanMX4 | Prof. Maya Schuldiner |
| *ypc1Δ* | BY4741; Mat a; his3Δ 1; leu2Δ 0; met15Δ 0; ura3Δ 0; YBR183w:: kanMX4 | Prof. Maya Schuldiner |
| *lip1Δ* | BY4741; Mat a; his3Δ 1; leu2Δ 0; met15Δ 0; ura3Δ 0; YMR298w:: kanMX4 | Prof. Maya Schuldiner |
| WT-Lag1p+GFP | gfp was tagged endogenously and regulated by its promoter | Prof. Maya Schuldiner |
| WT-Lac1p+GFP | gfp was tagged endogenously and regulated by its promoter | Prof. Maya Schuldiner |
| *DH5α* | F‐ϕ80dlacZΔM15Δ (lacZYA‐argF)U169 deoR recA1 endA1 hsdR17 (rk‐mk‐) phoA supE44λthi‐1 gyrA96 relA1 | Invitrogen |
| pRS416 | Empty vector | Addgene |
| pRS416-LAC1 | LAC1 clone constructed with URA3 promoter | This study |
| pRS416-LAG1 | LAG1 clone constructed with URA3 promoter | This study |
| pYES2 | Empty vector |  |
| pYES2-LAC1 | LAC1 clone was constructed with GAL1 promoter | This study |
| pYES2-LAG1 | LAG1 clone was constructed with GAL1 promoter | This study |
| pRS314+UPRE-GFP | GFP is driven by unfolded Protein Response Element (UPRE)- TRP1 selectable marker | Addgene |

**Table S2. Primers used in this research**

| **Real-Time PCR Primers** | | |
| --- | --- | --- |
| **Primer Name** | **Forward Oligonucleotide sequence (5′→3′)** | **Reverse** |
| LAG1 | GTTTCAGGGCCCTTTGGTCT | ATGCCGCTTGACCCAAGTAA |
| HAC1 | GAAGACGCGTTGACTTGCAG | ACGTATCAGACGACGAGTGC |
| IRE1 | AAGGCATCCGTTGTTTTGGC | AGTCAGAACCGGCGTCAAAT |
| RPN4 | GCTTCGATACCCCCACAACA | TCTATCGTTGGCCGTTGCTT |
| KAR2 | ACCAAGTTGCTGCCAATCCT | CAGCGGGCTTCCCATCTTTA |
| CHO2 | TATTTGCCTATCCAGAAGAGATCAAC | TGTAAGTCACAAAATCTTGCATCA AG |
| OPI3 | GGGCCAGAAAGGGCTGTTAC | ACACGTAGGCTGTTCACGA |
| PSD1 | CACTCCCACCCCTTGATGTC | CCCTGGCGAATTCTCTATGC |
| PSD2 | GCCACAAGATTATCACCGGTTT | GAACGGCCATTGGATTTACAG |
| DGA1 | TGACTATCGCAACCAGGAATGT | AACGCACCAAGTGCTCCTATG |
| LRO1 | CGTACAACCCTGCCGCCGGAAT | GTCTACGTGTTCGGCGCTTT |
| ARE1 | TGTTCCCCGTTCCTCGTGTA | CGCACACCTTCTCCAACACA |
| ARE2 | GCAACTCACCAGCCAATGAA | ATGCGACGTCTCCGTTTGA |
| INO2 | TCAACCAAGCATGGGTTTTG | AAAACTGTTCAATGGCATTCGA |
| INO4 | AGCTAAGCATGAGGCAAAAACC | CCCAAATTAACTCCTTCGGTACTAA |
| ORM1 | ATCAAGGAACTGCTGAGGCG | TCTTTCACAGGTTCCGTCGT |
| ORM2 | CCAGTACTACCGCAACGACA | GCCCGTAATACCAGGGATGG |
| ACT1 | ACTTTCAACGTTCCAGCCTTCT | ACACCATCACCGGAATCCAA |
| **Deletion Primers** | | |
| **Primer Name** | **Forward** | **Reverse** |
| LEU2  Cassette | AACTGTGGGAATACTCAGGTATCGTAAG | CAAATTAGGGATTCGTAGTTTCATGATTTTCTG |
| LEU2 Screening | CTCTTTGCCAGACAAGAACACCGCATTT | CCAAATGCGGTGTTCTTGTCTGGCAAAGAG |
| LAC1 deletion | CATACCTCCGGTAAACATTTAGATAGACACAGTATCAATAAACAAGAGCTAACTGTGGGAATACT | GTAAAGAATTAATGTGTAATGGTTATACTACTTAAAAACACCGTTTTCCTCAAATTAGGGATTCG |
| LAG1 deletion | TGTTGAGAGTGAACTCCAAGATACAGAGAAACTGAAGAAATAACGACAACAACTGTGGGAATACT | TCATACAGGGGGGAAATCATATGATGATACGTATTCTCCTTAAGATACGTCAAATTAGGGATTCG |
| ORM2 deletion | ATTAACGCAAGACTATACCATTATAAAAACGCATAAGAAACAGTTTCATCAACTGTGGGAATACT | ATACATATATATATATATATATATACATATATGCGTATAGGCAGAGCCAACAAATTAGGGATTCG |
| YDC1 deletion | TCATAGAACCTAGTGAATTTTTAAGAAAGTAAGATAAAGAAAAAAATCAAAACTGTGGGAATACT | TATACGTACATATATTTTGAAGATTCAAATGGATGGCACAAAATCACTCCCAAATTAGGGATTCG |
| **Splicing Primer** | | |
| HAC1 Splicing | AGGAAAGGCAGCGAAGG | GAATTCAAACCTGACTGCGC |
| **Cloning Primers** | | |
| Primer Name | Forward Primer (Restriction enzyme used) | Reverse primer (Restriction enzyme used) |
| LAC1 pYES2 | AATAGAAGCTTATGTCGACAATAAAGCCAA (*HindIII*) | ATAAACTCGAGAATATCCTTTTTCGTTGGA (*XhoI)* |
| LAG1 pYES2 | ATAAAGCTTATGACATCAGCTACGGACAAATC (*HindIII*) | ATACTCGAGTTCACACTTTTCCTTAGATTCTT (*XhoI*) |
| LAC1 pRS416 | GATGGTACCATGTCGACAATAAAGCCAA *(KpnI)* | GGTGGATCCAATATCCTTTTTCGTTGGA *(BamHI)* |
| LAG1 pRS416 | GATGGTACCATGACATCAGCTACGGACAAATC*(KpnI)* | GGTGGATCCTTCACACTTTTCCTTAGATTCTT*(BamHI)* |

Underlining indicates restriction enzyme sites.

**Supporting Results:**

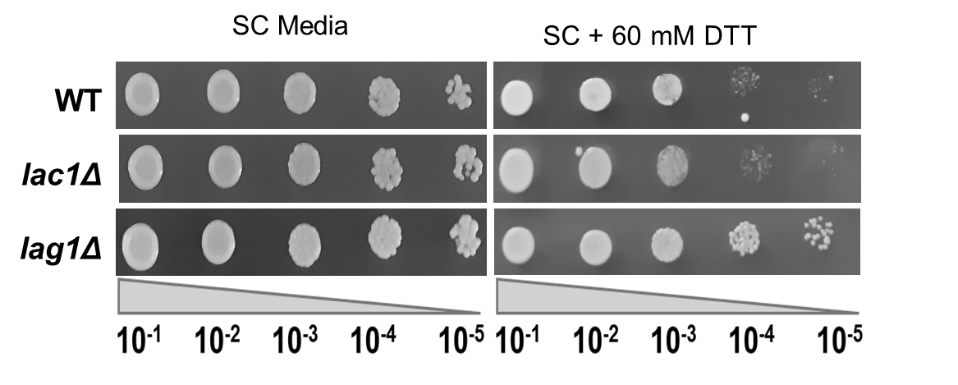

**Figure S1. *LAG1* deletion mitigates DTT-induced cell growth inhibition.** Cells with the same OD (3.0 at A_600 nm_) were serially diluted (10-fold) and grown as described in the methods section. These cells were spotted on SC agar plates, with or without DTT, and incubated for 48 hours at 30^o^C.

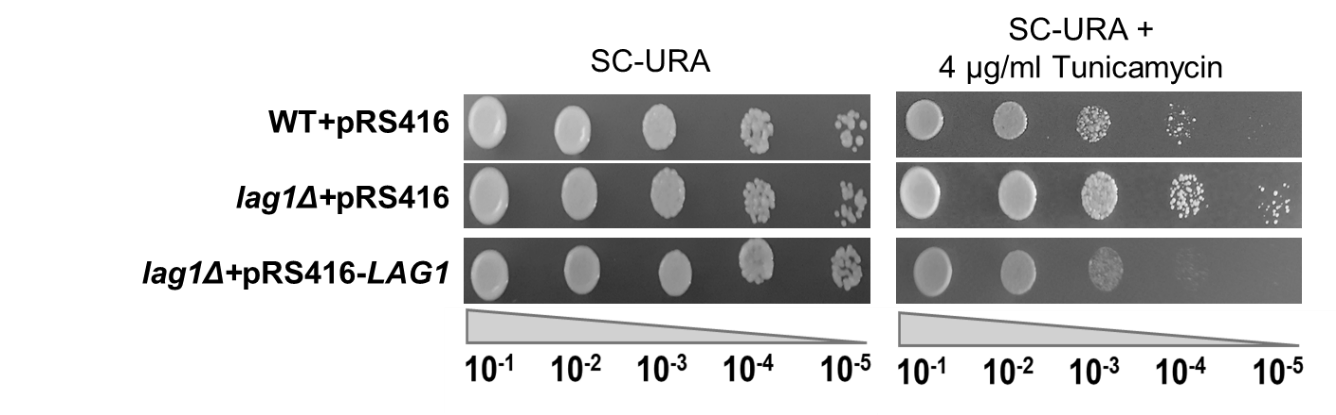

**Figure S2. *LAG1* complementation in *lag1Δ* cells reversed its ER stress-protective effects.** Cells with the same OD (3.0 at A_600 nm_) were serially diluted (10-fold) and grown as described in the methods section. These cells were then spotted on SC agar plates, with or without tunicamycin, and incubated for 48 hours at 30^o^C.

A) **B)**

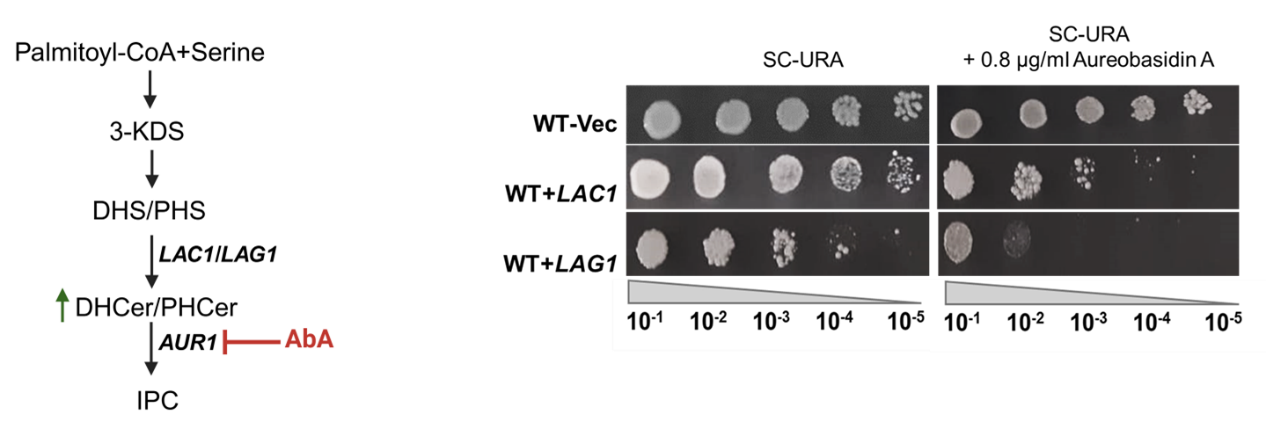

**C)**

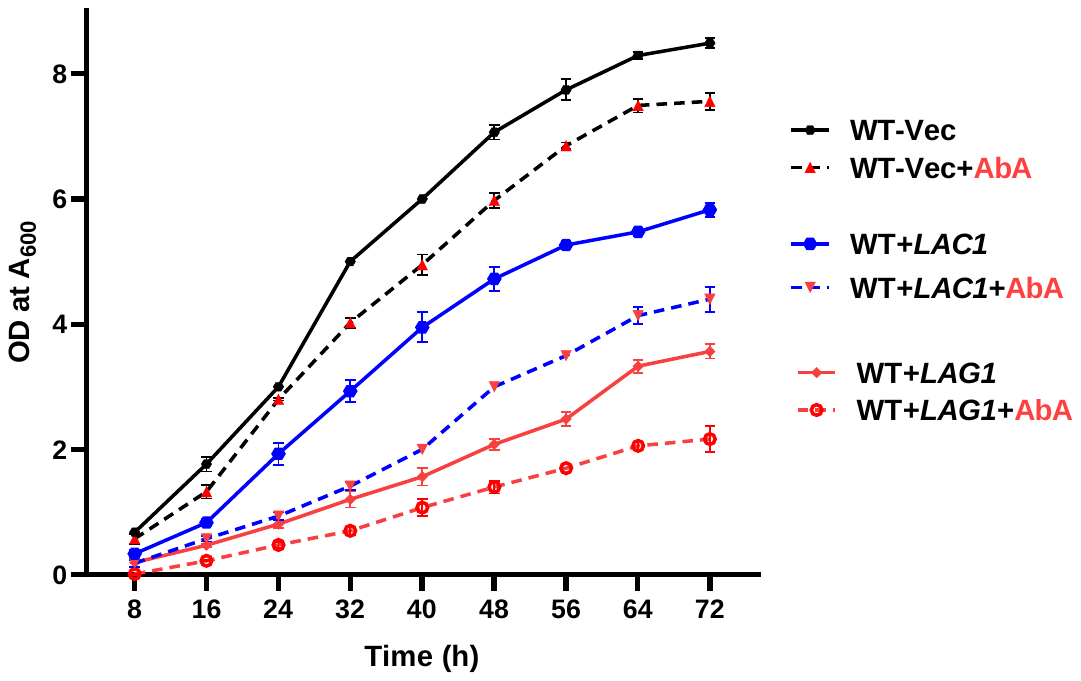

**D)	E)**

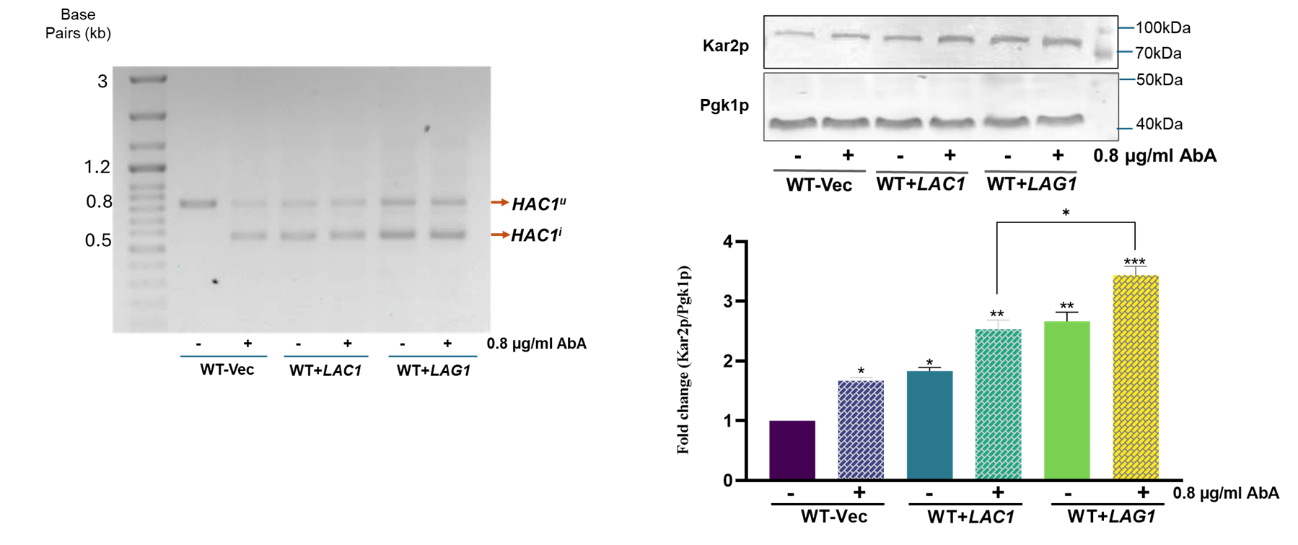

**Figure S3. Aureobasidin A treatment intensified the growth defects and ER stress in *LAG1*-overexpressing cells. (**A) Overview of the role of AbA in ceramide synthesis. (B) Cells with the same OD (3.0 at A_600 nm_) were serially diluted (10-fold) and grown as described in the methods section. These cells were spotted on SC-URA+ Galactose agar plates, with or without AbA, and incubated for 48 hours at 30^o^C. (C) Growth of WT-Vec, WT+*LAC1*, and WT+*LAG1* strains in SC-URA+ Galactose medium without and with 0.8 µg/ml AbA. The graphs show absorbance at 600 nm measured every 8 hours, with time points displayed on the x-axis using a plate reader. (D) The *HAC1* mRNA splicing experiment was carried out, and the resulting PCR products were separated on a 1% agarose gel. (E) A 12% SDS-PAGE gel was loaded with identical protein concentrations. The primary antibody used was anti-Kar2p, while Pgk1p was the loading control. The data represents the mean ± SD (*P<0.05) of three replicates from three separate experiments.

**A)**

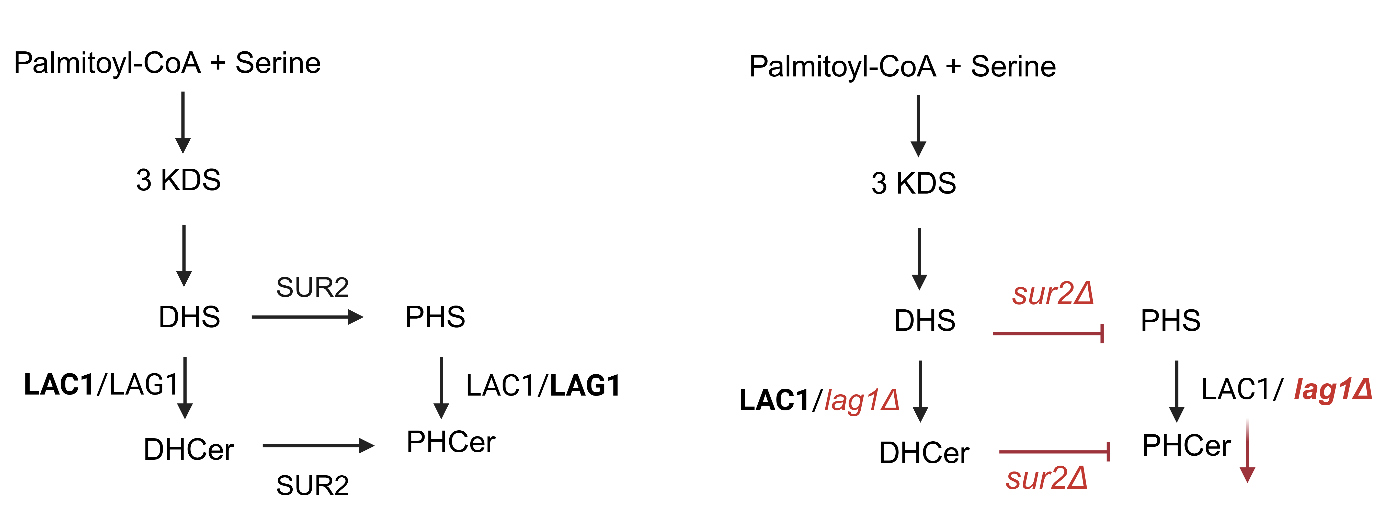

**B)**

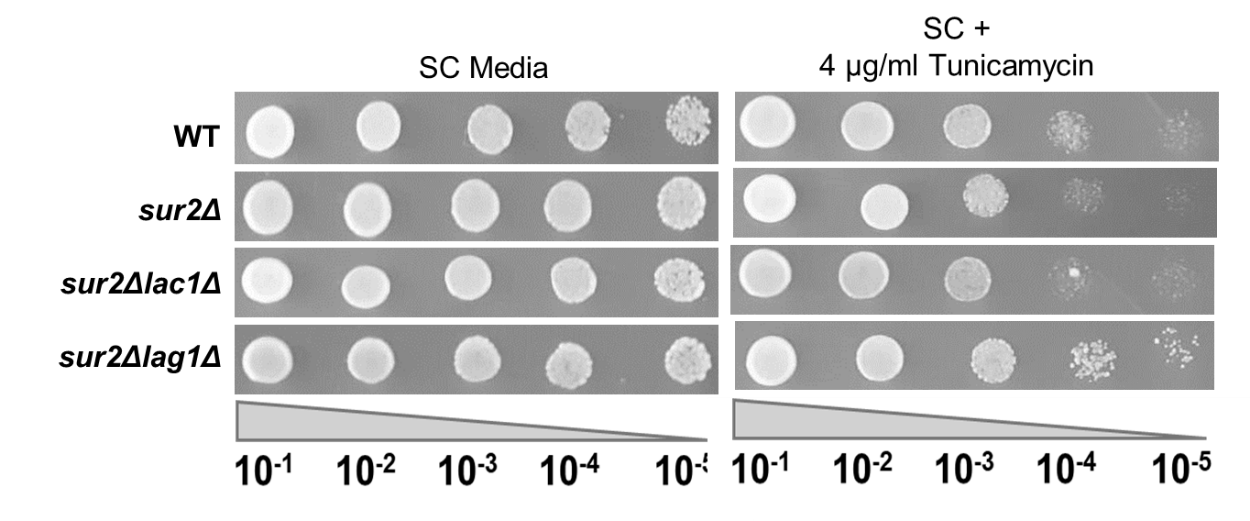

**C)	D)**

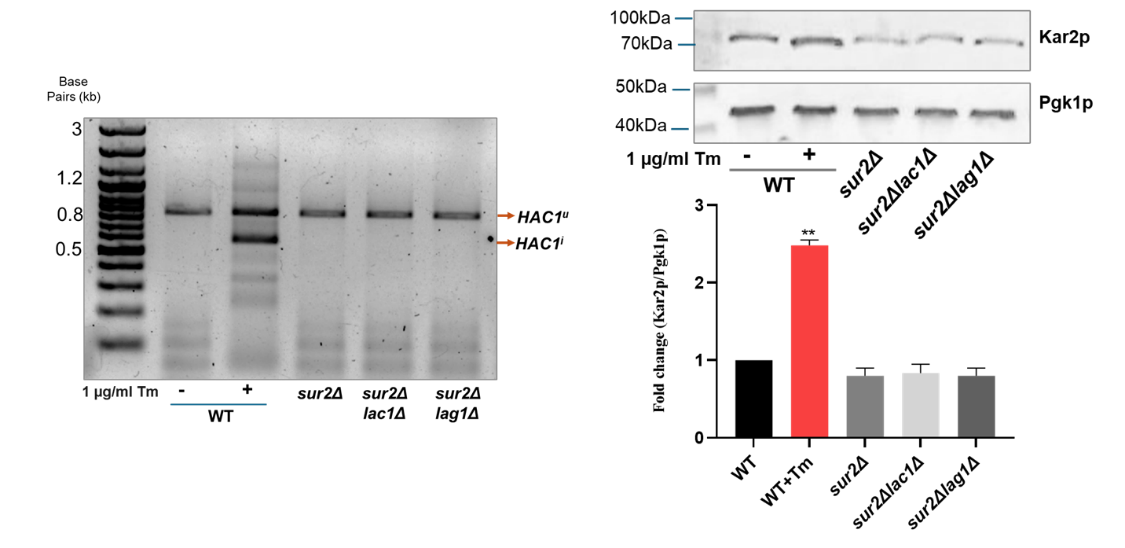

**Figure S4.** **Double deletion of *SUR2* and *LAG1* remains to confer ER stress resistance**: (A) overview of *SUR2* in ceramide biosynthesis. (B) Cells with the same OD (3.0 at A_600 nm_) were serially diluted (10-fold) and grown as described in the methods section. These cells were then spotted on SC agar plates, with or without tunicamycin, and incubated for 48 hours at 30 degrees Celsius. (C) The *HAC1* mRNA splicing experiment was carried out, and the resulting PCR products were separated on a 1% agarose gel. (D) A 12% SDS-PAGE gel was loaded with identical protein concentrations. The primary antibody used was anti-Kar2p, while Pgk1p was the loading control. The data represents the mean ± SD (*P<0.05) of three replicates from three separate experiments.

**A)**

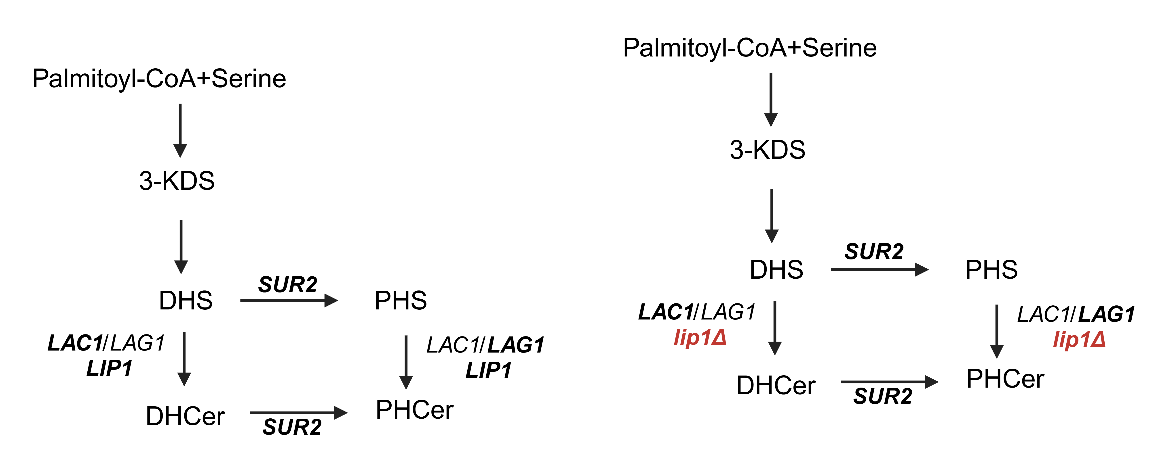

**B)**

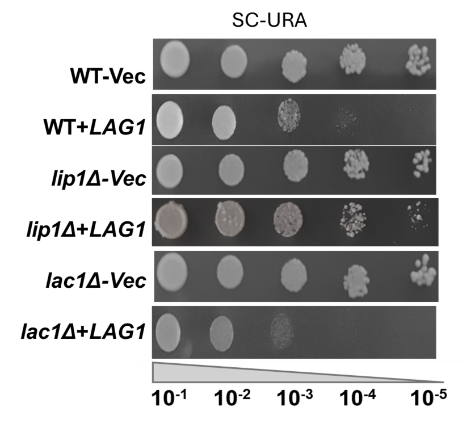

**C)**

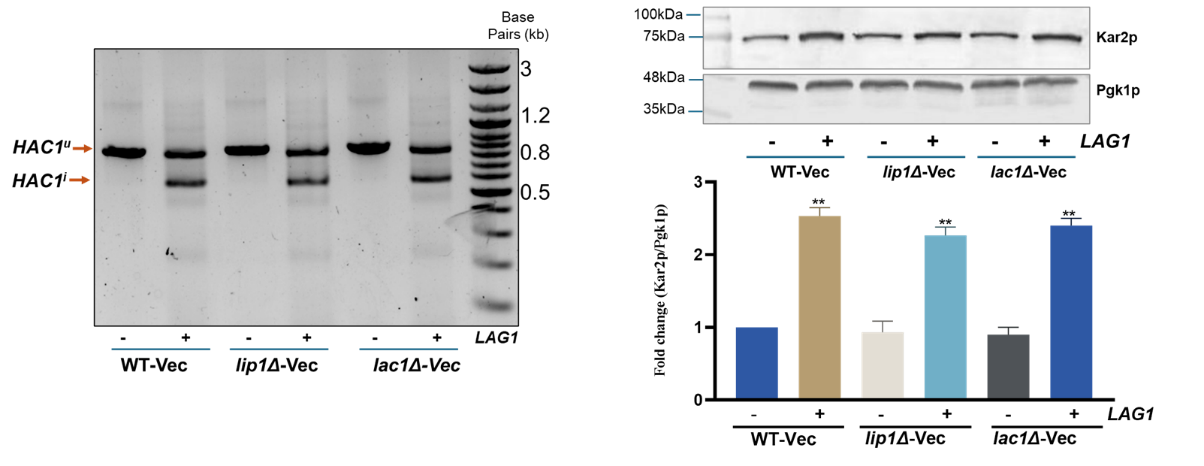

**Figure S5: Deleting *LIP1*, but not *LAC1*, slightly rescued the growth of cells overexpressing *LAG1*.** (A) Overview of the role of *LIP1* and *LAC1* in ceramide synthesis. (B) Cells with the same OD (3.0 at A_600 nm_) were serially diluted (10-fold) and grown as described in the methods section. These cells were spotted on SC agar plates, with or without AbA, and incubated for 48 hours at 30^o^C. (C) The *HAC1* mRNA splicing experiment was carried out, and the resulting PCR products were separated on a 1% agarose gel. (D) A 12% SDS-PAGE gel was loaded with identical protein concentrations. The primary antibody used was anti-Kar2p, while Pgk1p was the loading control. The data represents the mean ± SD (*P<0.05) of three replicates from three separate experiments.

**A)**

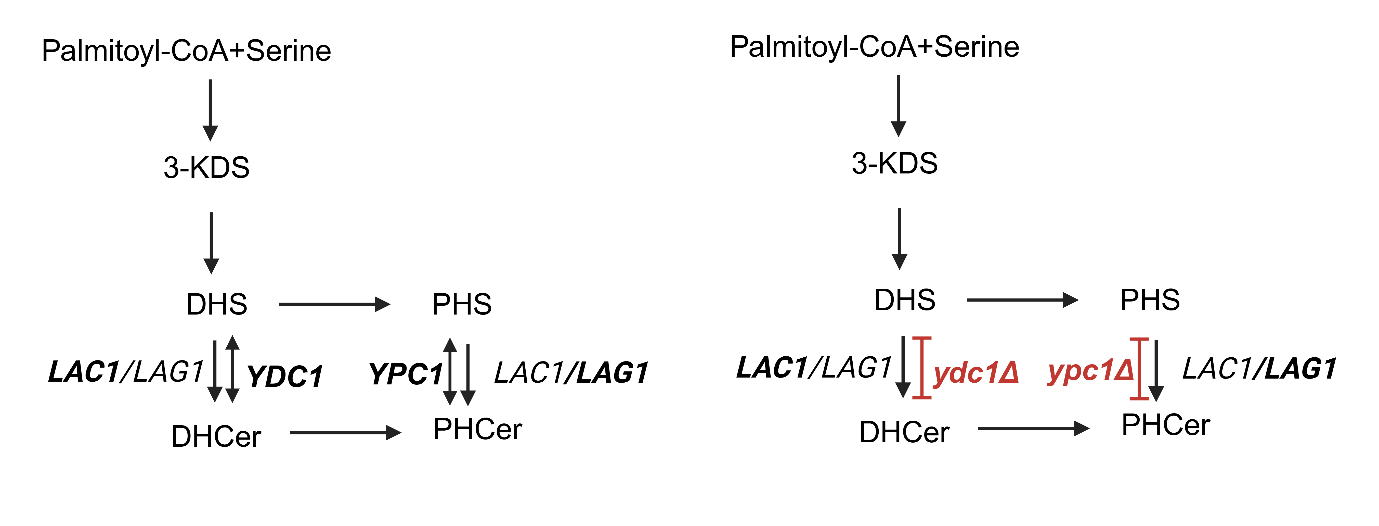

**B)**

`
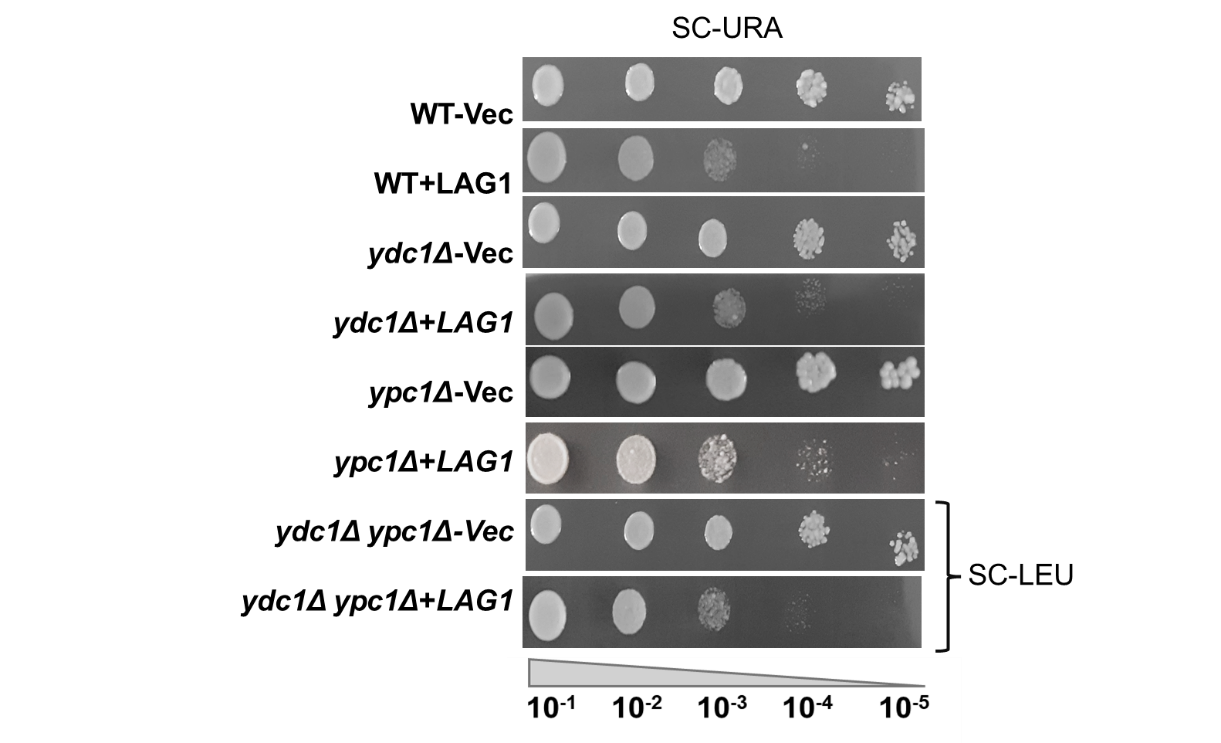

**C)	D)**

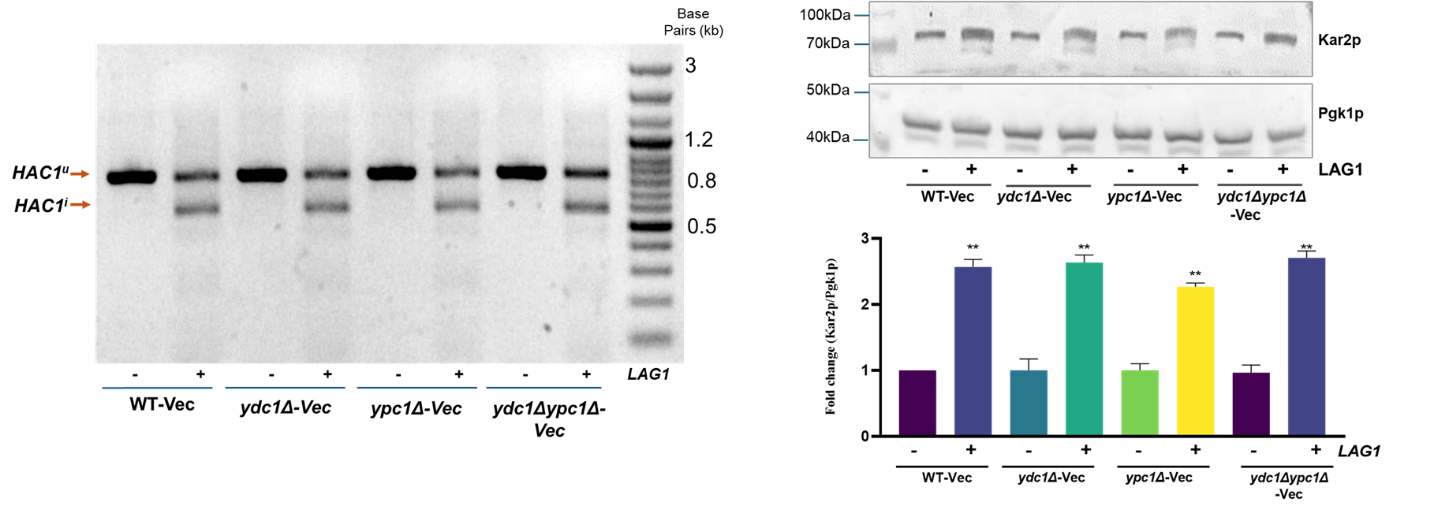

**Figure S6: Deleting ceramidases *YDC1* and *YPC1* has not changed the growth pattern and ER stress response in *LAG1* overexpression cells. (**A) Overview of the role of *YDC1* and *YPC1* in ceramide synthesis. (B) Cells with the same OD (3.0 at A_600 nm_) were serially diluted (10-fold) and grown as described in the methods section. These cells were spotted on SC agar plates, with or without AbA, and incubated for 48 hours at 30 degrees Celsius. (C) The *HAC1* mRNA splicing experiment was carried out, and the resulting PCR products were separated on a 1% agarose gel. (D) A 12% SDS-PAGE gel was loaded with identical protein concentrations. The primary antibody used was anti-Kar2p, while Pgk1p was the loading control. The data represents the mean ± SD (*P<0.05) of three replicates from three separate experiments.
